# Characterization and automatic classification of preterm and term uterine records

**DOI:** 10.1101/349266

**Authors:** Franc Jager, Sonja Libenšek, Ksenija Geršak

## Abstract

Predicting preterm birth is uncertain, and numerous scientists are searching for non-invasive methods to improve its predictability. Current researches are based on the analysis of ElectroHysteroGram (EHG) records, which contain information about the electrophysiological properties of the uterine muscle and uterine contractions. Since pregnancy is a long process, we decided to also characterize, for the first time, non-contraction intervals (dummy intervals) of the uterine records, i.e., EHG signals accompanied by a simultaneously recorded external tocogram measuring mechanical uterine activity (TOCO signal). For this purpose, we developed a new set of uterine records, TPEHGT DS, containing preterm and term uterine records of pregnant women, and uterine records of non-pregnant women. We quantitatively characterized contraction intervals (contractions) and dummy intervals of the uterine records of the TPEHGT DS in terms of the normalized power spectra of the EHG and TOCO signals, and developed a new method for predicting preterm birth. The results on the characterization revealed that the peak amplitudes of the normalized power spectra of the EHG and TOCO signals of the contraction and dummy intervals in the frequency band 1.0-2.2 Hz, describing the electrical and mechanical activity of the uterus due to the maternal heart (maternal heart rate), are high only during term pregnancies, when the delivery is still far away; and they are low when the delivery is close. However, these peak amplitudes are also low during preterm pregnancies, when the delivery is still supposed to be far away (thus suggesting the danger of preterm birth); and they are also low or barely present for non-pregnant women. We propose the values of the peak amplitudes of the normalized power spectra due to the influence of the maternal heart, in an electro-mechanical sense, in the frequency band 1.0-2.2 Hz as a new biophysical marker for the preliminary, or early, assessment of the danger of preterm birth. The classification of preterm and term, contraction and dummy intervals of the TPEHGT DS, for the task of the automatic prediction of preterm birth, using sample entropy, the median frequency of the power spectra, and the peak amplitude of the normalized power spectra, revealed that the dummy intervals provide quite comparable and slightly higher classification performances than these features obtained from the contraction intervals. This result suggests a novel and simple clinical technique, not necessarily to seek contraction intervals but using the dummy intervals, for the early assessment of the danger of preterm birth. Using the publicly available TPEHG DB database to predict preterm birth in terms of classifying between preterm and term EHG records, the proposed method outperformed all currently existing methods. The achieved classification accuracy was 100% for early records, recorded around the 23rd week of pregnancy; and 96.33%, the area under the curve of 99.44%, for all records of the database. Since the proposed method is capable of using the dummy intervals with high classification accuracy, it is also suitable for clinical use very early during pregnancy, around the 23rd week of pregnancy, when contractions may or may not be present.

## Introduction

Preterm birth, also referred to as premature birth or premature delivery, is defined by the World Health Organization (WHO) as the live delivery of babies which occur before 37 weeks of gestation [1]. Term births are the live delivery occurring at 37-42 weeks. According to WHO, the prevalence of preterm birth is 1 in 10 babies, or 15 million babies every year. This presents a serious problem since preterm delivery is the leading cause of morbidity of babies, and accounts for approximately 50% of all perinatal deaths [2]. Up to 40% of survivors of preterm delivery may develop numerous health defects as well as long-term disabilities in the neurological sense. An early prediction of an impending preterm labor could improve general newborn health. Appropriate medical intervention can be taken early to postpone the labor as long as possible.

Besides medically indicated or induced preterm births [2], and Preterm Premature Rupture of Membranes (PPROM) [2], evidence suggests that different pathological processes that might be involved in initiating preterm labor, such as uterine ischemia, burst blood vessels, intrauterine infection or inflammation, uterine over-distention [2], and numerous of other risk factors, such as diabetes, conization, hypertension, uterine abnormalities, smoking, alcohol and drug use, and life style, have also been identified [3].

Predicting preterm birth based on these factors alone is far from certain. Other techniques are needed for better prediction. One such promising technique is the analysis of an electromyogram (EMG) of the uterus recorded from the abdominal wall of a pregnant woman, i.e. ElectroHysteroGram (EHG), which allows the non-invasive quantitative assessment of mechanical uterine contractions present during pregnancy, which are the result of discontinuous bursts of action potentials due to spontaneous electrical discharges from the uterine muscle [4–
9].

Applying different prediction methods showed that EHG records seem to provide adequate data to predict preterm labor [7], and can diagnose labor more accurately than other traditional clinical methods [5] [7,10–12].

Measuring mechanical uterine pressure using an external tocodynamometer (external tocography) is another way of monitoring mechanical uterine contractions during pregnancy. Initially, it was thought that tocography, monitoring mechanical uterine contractions, would be a promising approach for predicting the risk of preterm birth, however later studies on monitoring, and quantitative analysis of uterine activity [10,13], have not confirmed this. However, tocography signals were successfully used for detection [14] and classification [15] of uterine contractions.

The task of predicting preterm birth on the basis of EHG uterine records is usually performed by two approaches: to distinguish between pregnancy and labor, either in preterm or term cases, or, to distinguish between preterm and term delivery. Both approaches may be divided into two categories: approaches dealing with individual contraction bursts corresponding to uterine contractions, and approaches dealing with the entire EHG records or signals. The uterus is composed of billions of intricately interconnected cells whose responses are non-linear, therefore it is regarded as a complex, non-linear dynamic system. A variety of linear and non-linear signal processing techniques have been used for predicting preterm birth. The selected parameters (or features) for classification were time domain parameters, parameters based on signal power spectrum, entropy parameters, parameters estimating non-linearity, and EHG propagation parameters.

Approaches to classifying individual pregnancy and labor contraction bursts incorporated the following features: Root Mean Square (RMS) value [6,16]; amplitude and area under contraction curve [16]; contraction power [16]; peak frequency of power spectrum [10,17–19]; mean frequency, peak frequency, and median frequency of power spectrum [16,19–21]; mean power frequency [22]; EHG propagation velocity [10,17]; wavelets [11] [20,23,24]; autoregressive (AR) coefficients [23]; time reversibility [20–22]; sample entropy and Lyapunov exponent [19–21]; variance entropy [20]; delay vector variance [21]; approximate entropy [22]; non-linear correlation coefficient [25]; and intrinsic mode functions using Empirical Mode Decomposition (EMD) [26].

The appearance of the Term-Preterm EHG Database (TPEHG DB) [27,28] resulted in a number of signal processing approaches dealing with entire EHG records or signals and non-linear features. These include: RMS, peak frequency and median frequency of power spectrum, and sample entropy [27,29–33]; autocorrelation zero crossing, maximal Lyapunov exponent, and correlation dimension [27]; entropy of intrinsic mode functions using EMD [34]; fractal dimension, fuzzy entropy, interquartile range, mean absolute deviation, mean energy, mean Teager-Kaiser energy, sample entropy, and standard deviation after EMD combined with wavelet packet decomposition [35]; and Multivariate Multiscale Fuzzy Entropy (MMFE) [36]. For successful classification, standard classifiers, e.g., *k* nearest neighbour [36], support vector machine [35], quadratic discriminant analysis [33], as well as advanced classifiers, such as variety of neural networks [32], random forests [31], and AdaBoost [34], were used.

Despite extensive research and some excellent achievements, the task of predicting preterm birth is still not as sufficiently solved as we would like. It is not quite clear what would be the “best” frequency content of EHG signals to extract features for classification, nor what would be the “best” features in order to achieve as high a classification accuracy as possible. These two questions are closely related.

Many methods to assess the danger of preterm birth use the frequency band 0.34-1.0 Hz of EHG signals [10,17,18,29,31,32]. This is done to avoid maternal respiration in the frequency band 0.2-0.34 Hz (assuming a respiratory rate from 12 to 20.4 breaths per min), and to avoid the influence of maternal electrocardiogram (ECG) in terms of maternal heart rate, a strong component, above 1.0 Hz (assuming a maternal heart rate of 60 beats per min [bpm] and higher) together with high frequency harmonics [37]. On the other hand, there is evidence that useful information regarding electrical activity of uterine bursts lies in the frequency band 0.1-4.7 Hz divided into two frequency bands: 0. 1-1.2 Hz, Fast Wave Low (F*W*_L_), and 1.2-4.7 Hz, Fast Wave High (F*W*_H_) [11,38]. The FWL is assumed to be related to the propagation of the electrical activity along the uterus, while the F*W*_H_ is assumed to be related to the excitability of the uterus [11,12,38,39]. The shift of the spectrum of uterine bursts towards higher frequencies as labor approaches was reported [4, 11, 12]. Therefore, many other methods used a wider frequency band expanding above 1.0 Hz, 0.3-3.0 Hz [27,30,34–36], 0.1-3.0 Hz [20,24,26], 0.3-4.0 Hz [27,33], 0.2-8.0 Hz [23], 0.05-16.0 Hz [11].

To address the problem of the non-stationarity of the EHG signals, and using the frequency band expanding above 1.0 Hz, several multifrequency band decomposition approaches were employed. The wavelet packet transform multiresolution signal decomposition technique, splitting the frequency region 0.0-3.125 Hz of the input signals into eight packets of equal bandwidths, was used to investigate the way the energy distribution of the uterine EHG signal is modified during pregnancy and in labor [40], and to improve the classification accuracy of complex uterine EHG signals [24]. The EMD method decomposing the EHG signals into a number of intrinsic mode functions which were sequentially ranked from the high to the low frequency components of the frequency region 0.3-3.0 Hz [34], or in combination with the wavelet packet decomposition technique [35], was used to improve the classification accuracy of preterm and term records of the TPEHG DB.

In order to select the relevant features to classify pregnancy and labor contractions, the Jeffrey divergence distance, sequential forward selection method, and Binary Particle Swarm Optimization (BPSO) technique, using standard classifiers (*k* nearest neighbour, linear discriminant analysis, and quadratic discriminant analysis) were investigated and compared [20]. The identified common features relevant for the classification of contractions were: the wavelet related features and variance entropy. Using the BPSO technique and support vector machine classifier, the proposed features for classification of entire preterm and term EHG records were fractal dimension, fuzzy entropy, interquartile range, mean absolute deviation, mean energy, mean Teager-Kaiser energy, sample entropy, and standard deviation [35].

Pregnancy is a long process, and all the underlying physiological mechanisms of the uterus present during pregnancy, and their evolution, are still poorly understood. We believe that the indications of excitability of the uterus are not restrained to efficient contractile events, which represent only a small fraction of total duration of pregnancy, but important physiological mechanisms may also be present outside of the contraction intervals, and at higher frequencies. None of the studies using the EHG signals were dedicated to non-contraction intervals, nor separately investigated the frequency band above 1.0 Hz, where permanent maternal heart activity (maternal heart rate with higher harmonics) in the electrical sense is expected. The ECG activity is of strong potential, about 1 mV, while the potential of EHG bursts is about 50 μV; the ratio is about 20. The influence of the maternal ECG [5], as well as the influence of the maternal heart rate with higher harmonics [37], on the uterine EHG activity are known. Besides, none of the studies were dedicated to frequency analysis of the external tocogram (TOCO signal) in the frequency band above 1.0 Hz where permanent maternal heart activity (maternal heart rate) in the mechanical sense, i.e., mechanical “vibrating” of the uterus due to the heart beating, is expected. To better understand the behavior of uterine physiological processes, we also decided to “listen” to the uterus, for the first time during non-contraction intervals, through the entire spectrum of the EHG and TOCO signals, up to 5.0 Hz, and to pay special attention to the frequency band above 1.0 Hz, which carries information of the electrical and mechanical activity of the uterus due to the maternal heart. Inclusion and frequency analysis of the TOCO signal, the characteristics of the power spectra of EHG and TOCO signals above 1.0 Hz, and signal features, help to better understand and describe the physiological mechanisms involved during pregnancy.

The aims of this study were:

1. to develop a new set of uterine records (EHG signals accompanied by a simultaneously recorded TOCO signal) of pregnant women (preterm, term) and of non-pregnant women;
2. to characterize the uterine records of the new set in terms of normalized power spectra and spectrograms;
3. to test the hypothesis that the frequency region of the EHG and TOCO signals above 1.0 Hz containing the frequency components due to the influence of the maternal heart (maternal heart rate with higher harmonics) provides important features for the efficient prediction of preterm birth;
4. to test another hypothesis that the non-contraction intervals of uterine records are equally, or even more, important for the accurate prediction of preterm birth than the contraction intervals;
5. to develop a new and improved method for the automatic prediction of preterm birth;
6. to evaluate the classification performance of the new method using the newly developed set, and using the publicly available TPEHG DB.

First, we quantitatively characterized, in addition to the power spectra of uterine contraction intervals, the power spectra of non-contraction intervals, using the EHG and TOCO signals of pregnant and non-pregnant women of the newly developed set of uterine records. We showed that the influence of the maternal heart on the uterus in the electro-mechanical sense is measurable via frequency domain analysis of the EHG, and TOCO, signals in the frequency region above 1.0 Hz. We found that the variability of the normalized peak amplitude of the power spectra of the EHG and TOCO signals in the frequency band 1.0-2.2 Hz, reflecting the electro-mechanical influence of the maternal heart activity, is very important for predicting preterm birth, and propose it as a new biophysical marker. Using the newly developed set, we showed that the features (sample entropies, median frequencies of power spectra, and peak amplitudes of the normalized power spectra) obtained from non-contraction intervals provide quite comparable and slightly higher classification accuracy in classifying preterm and term deliveries than these features obtained from contraction intervals. In addition, we verified the extent of the influence of the maternal heart on a pregnant uterus in the electro-mechanical sense via classification accuracies obtained for the records of non-pregnant women versus the records of pregnant women. Finally, the classification performance of the proposed method to classify preterm and term deliveries on the basis of the entire EHG records of the TPEHG DB was evaluated.

The study’s design was approved by the National Medical Ethics Committee of the Republic Slovenia (No. 32/01/97, No. 108/09/09).

## Materials and methods

With the aim to develop a useful and improved automatic method for predicting preterm birth, we followed a general and widely accepted development process [29–36]:

1. select or construct a valid batabase for training and testing the model;
2. characterize the data and use effective mathematical expressions to formulate the features that reflect their correlation with the target classes;
3. develop or introduce an algorithm or method to operate the analysis;
4. evaluate the performance of the classification using cross-validation tests for the objective prediction of real-world performance.

### Term-preterm electrohysterogram databases

#### Term-Preterm ElectroHysteroGram DataSet with Tocogram

The newly developed Term-Preterm ElectroHysteroGram DataSet with Tocogram (TPEHGT DS) contains 26 three-signal 30-min uterine EHG records with the fourth signal of a simultaneously recorded external TOCO signal of pregnant women, and another five 30-min uterine records (EHG and TOCO signals) of non-pregnant women. The records were collected at the University Medical Centre Ljubljana, Department of Obstetrics and Gynecology. (All women gave their written, informed consent.) The recording equipment and the recording protocol (including position of the electrodes) were those which were also used during collecting the records of the TPEHG DB [27]. The records of the pregnant women belong to pregnancies that resulted in spontaneous preterm delivery (13 *preterm* records from eight pregnancies), and to pregnancies that resulted in spontaneous term delivery (13 *term* records from ten pregnancies). The mean recording time and standard deviation of the records of pregnant women was 30.2 (± 2.76) weeks of pregnancy. The mean delivery times of *preterm* and *term* records, were 33.7 (± 1.97) weeks and 38.1 (± 1.04), respectively. Fig 1 shows the position of electrodes to measure EHG signals. The EHG signals of the records were collected from the abdominal surface using four AgCl_2_ electrodes. The first acquired EHG signal (S1) was measured between the topmost electrodes (E2-E1), the second (S2) between the leftmost electrodes (E2-E3), and the third (S3) between the lower electrodes (E4-E3). The reference electrode was attached to the woman’s thigh. The fourth simultaneous signal was the external mechanical uterine pressure. It was acquired using a cardiotocograph (model HP8030) of which sensor measuring the mechanical uterine pressure was attached at the top of the fundus. This signal is also known as an external tocogram or TOCO signal. The analog TOCO signal was lead to one of the amplifiers of the A/D converter (the value of 150 *μ*V corresponds to a pressure of 1 Pa). Prior to sampling, the EHG signals and TOCO signal were filtered using an analog three-pole Butterworth filter with the bandwidth 0.0-5.0 Hz. The sampling frequency, *f*s, for the EHG and TOCO signals was 20 samples per second per signal.

**Fig 1.**
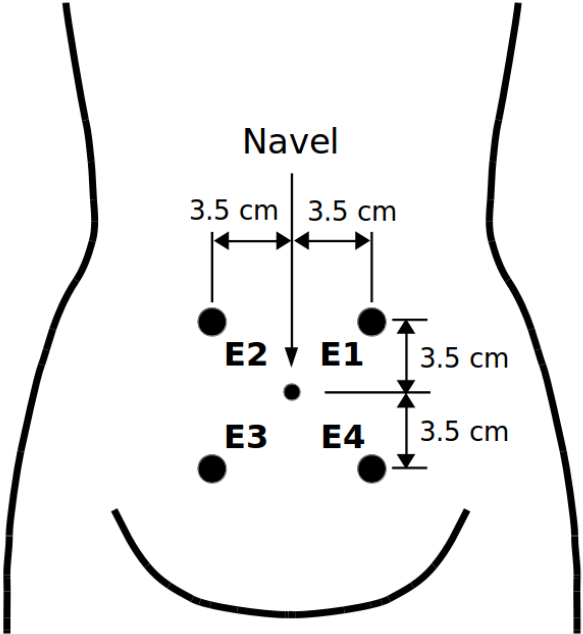
Position of electrodes. The electrodes were placed in two horizontal rows, symmetrically above and under the navel, spaced 7 cm apart.

There are 47 annotated intervals related to uterine contractions (*contraction* intervals) and 47 annotated non-contraction intervals (we named them *dummy* intervals) in the *preterm* records of the TPEHGT DS, another 53 annotated *contraction* and 53 *dummy* intervals in *term* records, and another 53 annotated *dummy* intervals in the records of non-pregnant women (*non-pregnant dummy* intervals). For the manual annotating procedure, we used our own graphic user interface and annotation editor. Besides visualizing original signals and annotation editing, the graphic user interface also allows calculating and visualizing spectrograms of the signals. Consensus about the annotated intervals was reached by two annotators. Fig 2 shows annotations, original signals, and spectrograms of original signals, of a *preterm* record of the TPEHGT DS after the procedure of manual annotating. The beginnings and ends of *contraction* intervals were set according to onsets and offsets of the deflections visible in the TOCO signal, which had to be accompanied by simultaneous severe bursts in the EHG signals. Ambiguous cases and cases contaminated with motion artefacts were not annotated. After that, in each record the same number of *dummy* intervals as the number of already annotated *contraction* intervals were annotated. The beginnings and ends of 6/50 *dummy* intervals were set in the signal intervals with no visible deflection in the TOCO signal, and with no simultaneous activity in the EHG signals, between, or next/prior to, *contraction* intervals, again avoiding motion artefacts. The lengths of *dummy* intervals were decided to be approximately of the same lengths as were the lengths of already annotated neighbouring *contraction* intervals, thus providing as much as possible comparable frequency resolution during Fourier decomposition. The lengths of *non-pregnant dummy* intervals in the records of non-pregnant women were decided again to be approximately of the same lengths as were the lengths obtained for annotated *contraction* intervals. Fig 3 shows annotations, original signals, and spectrograms of original signals, of a record of non-pregnant woman of the TPEHGT DS. The average lengths and standard deviations of *preterm* and *term contraction* intervals were 82 (± 48) sec and 88 (± 36) sec, of *preterm* and *term dummy* intervals were 83 (± 46) sec and 89 (± 46) sec, while of *non-pregnant dummy* intervals were 92 (± 37) sec.

**Fig 2.**
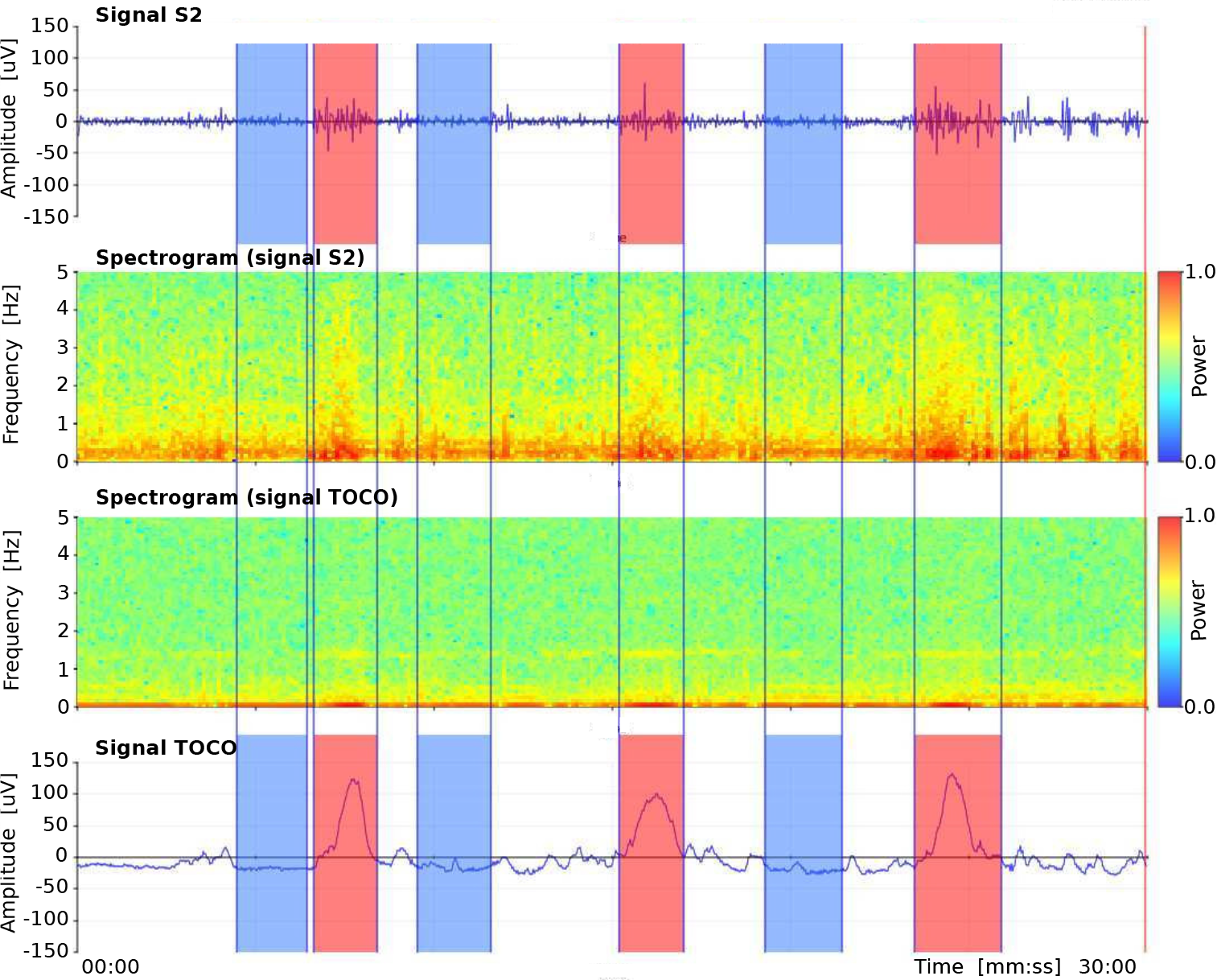
Manual annotations of three *contraction* and three *dummy* intervals in the record *tpehgt_p008* (*preterm*, recorded in the 26th week, delivery in the 32nd week) of the TPEHGT DS. From top to bottom: EHG signal S2, spectrogram (0.0-5.0 Hz) of EHG signal S2, spectrogram (0.0-5.0 Hz) of TOCO signal, and TOCO signal. Red: *contraction* intervals, blue: *dummy* intervals.

#### Term-Preterm EHG Database

The Term-Preterm EHG Database (TPEHG DB) [27], which is also publicly available on Physionet web site (https://physionet.org/physiobank/database/tpehgdb/) [28], contains 162 three-signal EHG records recorded around the 23rd week of gestation (*early* records), of which 19 ended in *preterm* delivery and 143 in *term* delivery, and another 138 EHG records recorded around the 31st week of gestation (*later* records), of which 19 ended in *preterm* delivery and 119 in *term* delivery. In total, there are 300 EHG records (all records), of which 38 ended in *preterm* delivery and 262 in *term* delivery. The records of the TPEHG DB do not contain the TOCO signal. The EHG signals S1, S2, and S3 were measured according to the positions of the electrodes as shown in Fig 1.

**Fig 3.**
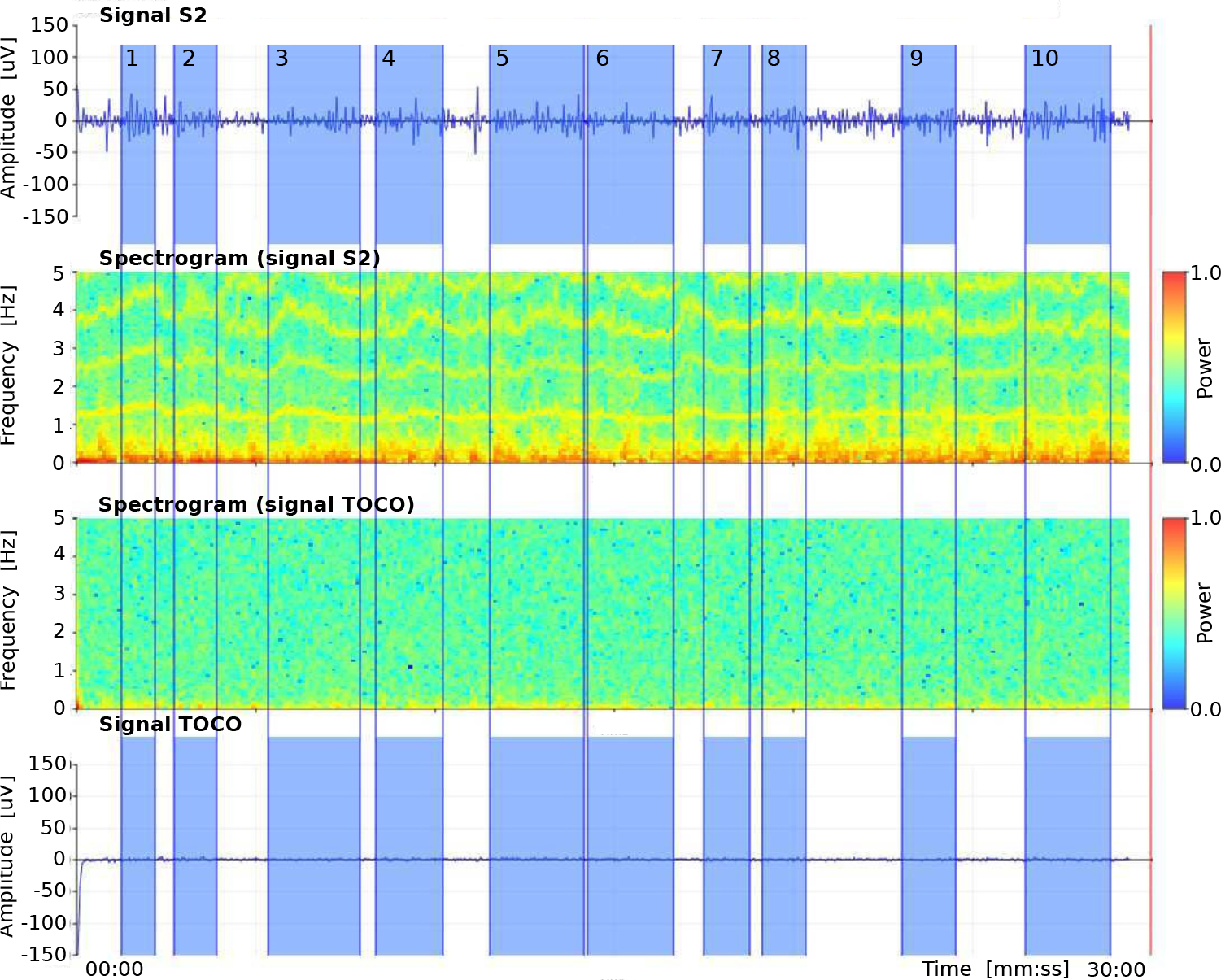
Manual annotations of *non-pregnant dummy* intervals in the record *tpehgt_n002* (non-pregnant) of the TPEHGT DS. From top to bottom: EHG signal S2, spectrogram (0.0-0.5 Hz) of EHG signal S2, spectrogram (0.0-0.5 Hz) of TOCO signal, and TOCO signal. Blue: *dummy* intervals.

#### Feature extraction technique and features

The EHG signals are non-stationary and time-varying. They carry important information about the non-linear processes of the underlying physiological mechanisms of the uterus present during pregnancy. In general, physiological mechanisms likely reside in different locally stationary frequency bands and with different intensities, whereas the corresponding frequency content and the intensities of the mechanisms in the bands vary as the pregnancy progresses. Activity connected to contractions in the EHG and TOCO signals of uterine records is expected below 1.0 Hz, while above 1.0 Hz separate and strong activity connected to another mechanism, the frequency component of the maternal ECG (heart rate) together with higher frequency harmonics, is expected. Therefore, the approach for the characterization and classification of *contraction* and *dummy* intervals of uterine records (composed from the EHG and TOCO signals), and of entire EHG records, is based on four strictly separated frequency bands:

- Band B0: *f*_low_ = 0.08 Hz, *f*_high_ = 1.0 Hz;
- Band B1: *f*_low_ = 1.0 Hz, *f*_high_ = 2.2 Hz;
- Band B2: *f*_low_ = 2.2 Hz, *f*_high_ = 3.5 Hz;
- Band B3: *f*_low_ = 3.5 Hz, *f*_high_ = 5.0 Hz.

The chosen frequency bands to analyze the EHG and TOCO signals allow: 1) characterization of the influence of the maternal heart in terms of heart frequency (B1), and its second (B2) and third (B3) harmonic, to electrical (EHG) and mechanical (TOCO) activity of the uterus separately from the influence of contractions (B0); 2) testing of the classification performance to predict preterm birth when the features are related to strictly separated frequency bands that correspond to different physiological mechanisms. The selected features for characterization, and classification of *contraction* or *dummy* intervals, or entire records, the median frequency of the power spectrum, and the peak amplitude of the normalized power spectrum. These features have the capability to directly estimate the presence and extent of the underlying physiological mechanisms in separate frequency bands.

#### Sample entropy

Sample entropy, *SE*, estimates the regularity or predictability of signals [41,42]. Less regular or less predictable signals exhibit a higher sample entropy. Let y(n) denote the input time series of length N and let *b*_*l*_ [0,…, *m* − 1] denote patterns of length *m, m* < *N*, where patterns *b*_*l*_ are taken from the time series *y*(*n*), *b_l_*(*i*) = *y*(*l* + *i*), 0 ≤ *i* ≤ *m* − 1, 0 ≤ *l* ≤ *N* − *m*. The part of the time series *y*(*n*) at index *n* = *n*_*s*_, *y*[*n_s_*,…, *n*_*s*_ + *m* — 1], is considered a match for a given pattern *b*_*l*_, *n*_*s*_ ≠ *l*, if max {|*y*(*n*_*s*_ + *i*) − *b*_*l*_(*i*)|: 0 ≤ *i* ≤ *m* − 1} ≤ *r*. The number of pattern matches, *c*_*m*_, is constructed for each *m*. The sample entropy, *SE*_*m,r*_(*y*), is then calculated following:

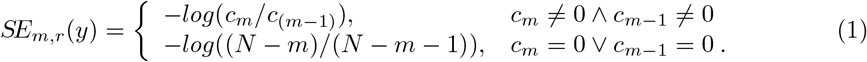

The following values of parameters were used, *m* = 3 and *r* = 0.15. These two values were adopted from the previous study assessing the separability between *preterm* and *term* EHG records of the TPEHG DB [27].

#### Median frequency of power spectrum

The median frequency, *MF*, of the power spectrum, *P*(*k*), in the selected frequency band between the low, *f*_low_, and high, *f*_high_, frequencies of interest was defined following:

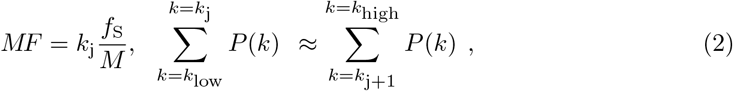

where 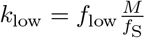 and 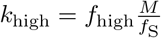 are the indexes of the power spectrum components at the low, *f*_low_, and high, *f*_high_, frequencies, *f*s is the sampling frequency, and *M* denotes the number of components of the power spectrum *P*(*k*).

#### Peak amplitude of normalized power spectrum

The peak amplitude of the normalized power spectrum, *PA*, in the selected frequency band between the low, *f*_low_, and high, *f*_high_, frequencies of interest was defined following:

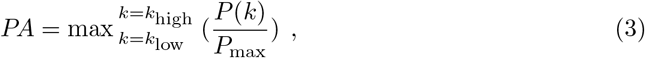

where 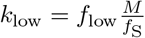 and 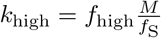 are the indexes of the power spectrum components at the low, *f*_low_, and high, *f*_high_, frequencies, and *P_max_* is the maximum component of the power spectrum in the frequency interval from 0 to 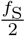.

The power spectra of signals of *contraction* or *dummy* intervals, or of entire signals of records, were calculated using the Fourier transform preceded by Hanning weighted windowing to attenuate spectral leakage. In order to reduce the fluctuations (small amplitude frequency spikes) in the power spectra, smoothing of the power spectra was performed using moving average over the frequency interval of 0.1 Hz, i.e., 1.0% of the interval from 0.0 Hz to 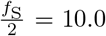. Normalization of the power spectrum with respect to the spectral peak in the entire spectrum allows the estimating of relative proportions of the *PA* in the selected frequency bands. For example, normalization allows the assessment of the intensity, *PA*, of the influence of the maternal heart in the frequency band B1 in comparison to the *PA* in other frequency bands, e.g., B0 where the influence of maternal respiration and contractions is expected. Moreover, higher and frequent uterine activity in signals of *term* records results in slightly higher EHG signal amplitude in an absolute sense than in signals of *preterm* records [5]. Normalization provides comparable estimates of relative proportions of peak amplitudes in separate frequency bands for both types of records. In addition, normalization of the power spectra of the EHG and TOCO signals allows comparison of relative proportions of peak amplitudes in separate frequency bands for both types of signals. The *f*_low_ of the frequency band B0 was intentionaly set to 0.08 Hz thus retaining the respiration component (0.2-0.34 Hz) in the power spectra of the EHG and TOCO signals of *dummy* intervals and *non-pregnant dummy* intervals, and preserveing at least one high spectral peak for normalization purposes.

The *SE* has been successfully applied for analysis of many biological signals such as ECG, blood pressure, electroencephalogram, and electromyogram [43]. The *SE* has the ability to estimate the level of regularity or predictability of time series. A low value of the *SE* suggests the presence of a physiologic mechanism with periodic behavior, while a high value suggests the absence of a mechanism. The *SE* and *MF* have been successfully used to classify individual pregnancy and labor contractions [19–21,44], and to classify entire *preterm* and *term* EHG records [27,29–33], which are actually sequences of *contraction* and non-contraction (*dummy*) intervals. The *MF* and *PA* are suitable features for assessing shifts and intensity of the frequency content in any biological signal and in separate frequency bands. The selected features are transparent and suitable for explaining the behaviour of the underlying physiological mechanisms present. They directly indicate and describe the mechanisms. While the *SE* estimates the presence or absence of a physiologic mechanism, given frequency band, the *MF* and *PA* estimate its median frequency and its intensity. Considering all these facts, the choice for *SE*, *MF*, and *PA* also seems suitable for analyzing *dummy* intervals of EHG and TOCO signals. The same features (*SE*, *MF*, *PA*) were used to analyze *contraction* and *dummy* intervals. If using the same features, it is more straightforward to compare characterization and classification results for different types of signal intervals.

### Spectrograms

A spectrogram allows for the visualization of the changes of the power spectrum of a signal over time. In order to derive spectrograms, the EHG and TOCO signals of the uterine records were initially preprocessed using the four-pole band-pass digital Butterworth filter, with cut-off frequencies at 0.08 Hz and 5.0 Hz, applied bi-directionally to yield zero-phase shift. The sliding Hanning window of duration of 12.8 sec (256 signal samples) and short-time Fourier transform were then used to derive the power spectra of the spectrograms.

#### Assessing separability, feature selection, and feature ranking

In order to estimate the ability of individual features to separate between *preterm* and *term*, *contraction* and *dummy* intervals, to separate between the entire *preterm* and *term* EHG records, and to assess the rank of their ability to classify preterm and term deliveries, we used the two-sample *t*-test with a pooled variance estimate [45], the Bhattacaryya criterion, i.e., the minimum attainable classification error or Chernoff bound [35, 46, 47] and the relative entropy criterion, also known as Kullback-Leibler distance or divergence [48].

We dealt with a large number of potential features that could be used for classification. Using all the features for classification may have negative impact on the classification performance due to the correlation between the features (redundant information), and overfitting may occur. Even though the entire set of features has explainable physiological interpretation or meaning, it is necessary to reduce the number of discriminative features to avoid overfitting, to simplify the classifier, and to improve classification accuracy. To select the best subsets of features for a variety of classification tasks in this study, we used the wrapper feature selection search strategy with the Sequential Forward Selection (SFS) method [49]. As a preprocessing step to select features, the features are sorted according to the selected criterion estimating the ability of individual features to separate groups. The available features of each of the classification tasks of this study were initially sorted according to their descending values of the Bhattacaryya criterion. The SFS method selects a subset of features by sequentially adding the features until a certain stopping condition is satisfied. Using the selected learning algorithm and the misclassification error (MCE), i.e., the number of misclassified observations divided by the total number of observations, as the performance indicator on each candidate features subset, the SFS sequentialy searches for features. Data are divided into training and test sets. The training set is used to select the features and to fit the selected model, while the test set is used to evaluate the performance of the selected features. In order to evaluate and compare the performance of each candidate feature subset, cross-validation is applied to the training set. The SFS algorithm stops when the first local minimum of the cross-validation MCE on the training set, as a function of the number of features, is found. However, the algorithm may stop prematurely. A smaller MCE may be found by looking for the minimum of the MCE function over a wider range of number of features. Therefore, the cross-validation MCE can be derived over the entire set of features available. When the curve of the MCE as a function of the number of features goes up, overfitting occurs. The minimum of the cross-validation MCE on the training set, as a function of the number of selected features, defines the number of needed features and their performance. In order to evaluate the performance of the selected features, 20% holdout dividing the data into training and test sets, 10-fold cross-validation applied to the training set, and the Quadratic Discriminant Analysis (QDA) learning algorithm, were used for each of the classification tasks of this study.

However, different runs of the SFS algorithm result in different selected feature subsets. To avoid this instability of the SFS method, and to form a stable subset of predictive features, we applied a procedure based on the frequency-based aggregation of the selected features by running the SFS algorithm a significant number of times and recording the selected feature subset after each run [50]. The aggregated frequency histogram of feature occurences was then used to form the final feature subset by selecting the most frequent features in rank order as indicated by histogram peaks [50]. The number of final selected features using this frequency-based feature-aggregation and feature-selection procedure was defined by the minimum of the average cross-validation MCE function as obtained on the training sets over repeated runs of the SFS algorithm. In this study, 200 runs were used in each case.

#### Proposed method for predicting preterm birth

Fig 4 shows the signal processing flow chart of the proposed method for predicting preterm birth. After manual annotating of *contraction* or *dummy* intervals, or neither, in a uterine record, **R**, the entire selected (or available) signals (EHG signals, or EHG signals and TOCO signal) of the record are preprocessed using a band-pass linear-phase filter with cut-off frequencies at 0.08 Hz and 5.0 Hz (to reject signal baseline wander and higher frequencies above 5.0 Hz) producing the record **R0**-**3**; and with a bank of band-pass linear-phase filters with cut-off frequencies corresponding to the frequency bands B0, B1, B2, and B3, producing subrecords **R0**, **R1**, **R2**, **R3**. Note that the TOCO signal is processed in the same manner as the EHG signals in all procedures. We propose using the fourth-order band-pass digital Butterworth filters for the strict separation of the four frequency bands, which are applied bi-directionally to each signal of the original record, **R**, to yield zero-phase shift. The Butterworth filters have a smooth, monotonically changing frequency response, are maximally flat (no ripples) in the passband and stopband, and are computationally non-intensive. The frequency response of the filters rolls off at −80 dB per decade, and after bi-directional use even at −160 dB per decade. After that, the sample entropy, *SE*, is derived for each signal of an annotated data interval (*contraction* or *dummy)*, or for each signal of the entire record, for each of the subrecords **R0**, **R1**, **R2**, **R3**, which correspond to the frequency bands B0, B1, B2, and B3. Next, the Hanning weighting window is applied to each signal of the annotated data interval, or to each signal of the entire record, of the record **R0**-**3** to attenuate spectral leakage, and the power spectra are calculated using the Fourier transform. The power spectra are then smoothed using a moving average over the frequency interval of 0.1 Hz, and normalized by their maximum components. Extraction of the median frequency, *MF*, and peak amplitude, *PA*, of the normalized power spectrum follows for each signal of the annotated data interval, or for each signal of the entire record, for each of the frequency bands: B0, B1, B2, and B3. Finally, the extracted features of the annotated data interval, or of entire record, lead to a classifier judging the danger of preterm birth.

**Fig 4.**
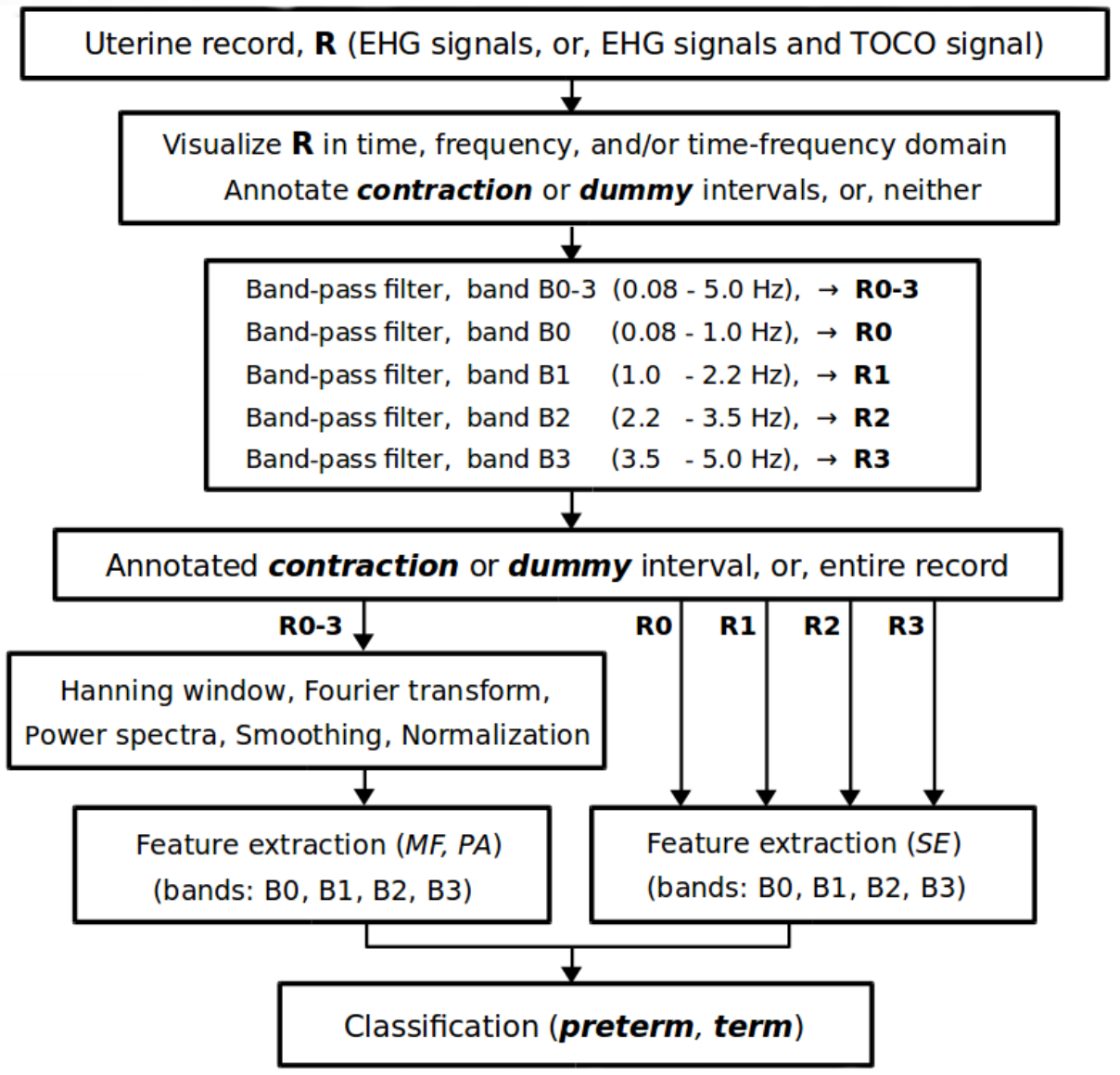
Signal processing flow chart of the proposed method for predicting preterm birth.

### Classification

The main task of this study was to justify the importance of the influence of the maternal heart on the uterus for the accurate prediction of preterm birth via variety of the classification tasks, and not necessarily to seek for the best classifier. To assess the classification performance of the variety of classification tasks, and for the easier and consistent comparison of the performance results obtained, the same classifier, i.e., the QDA classifier, was used in each case. In comparison to a few other standard classifiers, the QDA classifier already reliably classified between pregnancy and labor contractions [20], and between *preterm* and *term*, *early* and *later* records of the TPEHG DB [33], therefore the QDA classifier seems suitable for the domain of predicting preterm birth. In the scope of this study, we have also preliminarily tested a few other standard classifiers: naive Bayes, *k* nearest neighbour, linear discriminant analysis, decision tree, and support vector machine. Among the tested classifiers, if using the TPEHG DB, the highest classification accuracy obtained was for the QDA classifier. Moreover, the proposed method using the QDA classifier and the TPEHG DB outperformed all currently existing methods for predicting preterm birth. For these reasons, the QDA classifier seems suitable choice for this study.

Classification performance results were summarized in terms of Sensitivity, *Se* = *TP*/(*TP+FN*), Specificity, *Sp* = *TN*/(*TN+FP*), and Classification Accuracy, *CA* = (*TP+TN*)/(*TP+FN+TN+FP*), where *TP* denotes the number of true positives, *FN* the number of false negatives, *TN* the number of true negatives, and *FP* the number of false positives; and in terms of the Area Under the ROC Curve (*AUC*) [51] to assess the predictive confidence of discrimination accuracy. Cross-validation with five-folds (TPEHG DB), ten-folds (TPEHGT DS, TPEHG DB), and with 30 repetitions was used in each case.

To avoid an overfitting problem due to the unequal prior probabilities of the classes of datasets (a classifier would be more sensitive in detecting the majority class than the minority class), and to provide more accurate results in terms of the predictive confidence of discrimination accuracy, data balancing using the over-sampling approach was employed. The Standard Synthetic Minority Over-sampling Technique (SMOTE) [52] was used to balance the *contraction* and *dummy* intervals of the TPEHGT DS, while the Adaptive Synthetic Sampling Approach (ADASYN) [53] was used to balance *preterm* and *term* records of the TPEHG DB. The SMOTE technique forces the decision region of the minority class to become more general and provides an equal number of samples in both classes, while the ADASYN approach provides a balanced representation of data distribution in the resulting datasets.

## Results

### Characterization of *preterm* and *term, contraction* and *dummy* intervals

Records of the TPEHGT DS were separated into two groups, nonlabor and labor. There is little consensus regarding the definitions of labor onset in the research literature [54]. The boundary was set intentionally early, to three weeks, likely to the latent phase of labor or even earlier. Initially, several *preterm* and *term* records of the nonlabor and labor groups were characterized in terms of spectrogram time-frequency representation. Figs 5 and 6 show the spectrograms of the EHG signal S3 and TOCO signal of a *preterm* nonlabor record and of a *term* nonlabor record. For both records, the time interval between recording and delivery is ten weeks. In the spectrograms of EHG signals maternal heart activity above 1.0 Hz and with higher harmonics can be observed. The activity is stronger for the *term* record. Similarly, strong maternal heart activity above 1.0 Hz is present in the spectrogram of the TOCO signal for the *term* record, but it is absent in the spectrogram of the TOCO signal for the *preterm* record. (Note that the TOCO signal carries information about the mechanical activity of the uterus.) The absence of the maternal heart activity in the spectrogram of the TOCO signal can also be observed for *preterm* nonlabor record from Fig 2. These observations suggest a stronger presence of a maternal heart activity for *term* nonlabor records.

**Fig 5.**
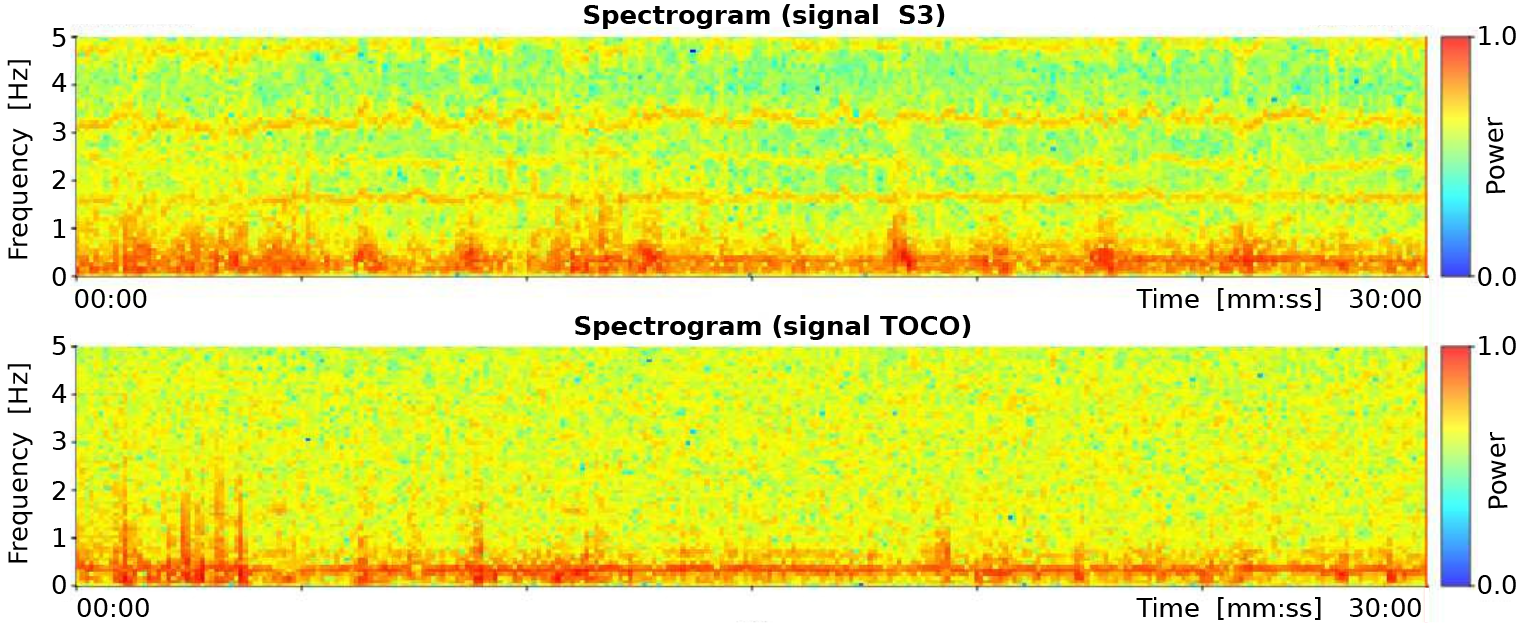
Spectrograms of the record tpehgt p006 (preterm nonlabor record, recorded in the 26th week, delivery in the 36th week) of the TPEHGT DS.

**Fig 6.**
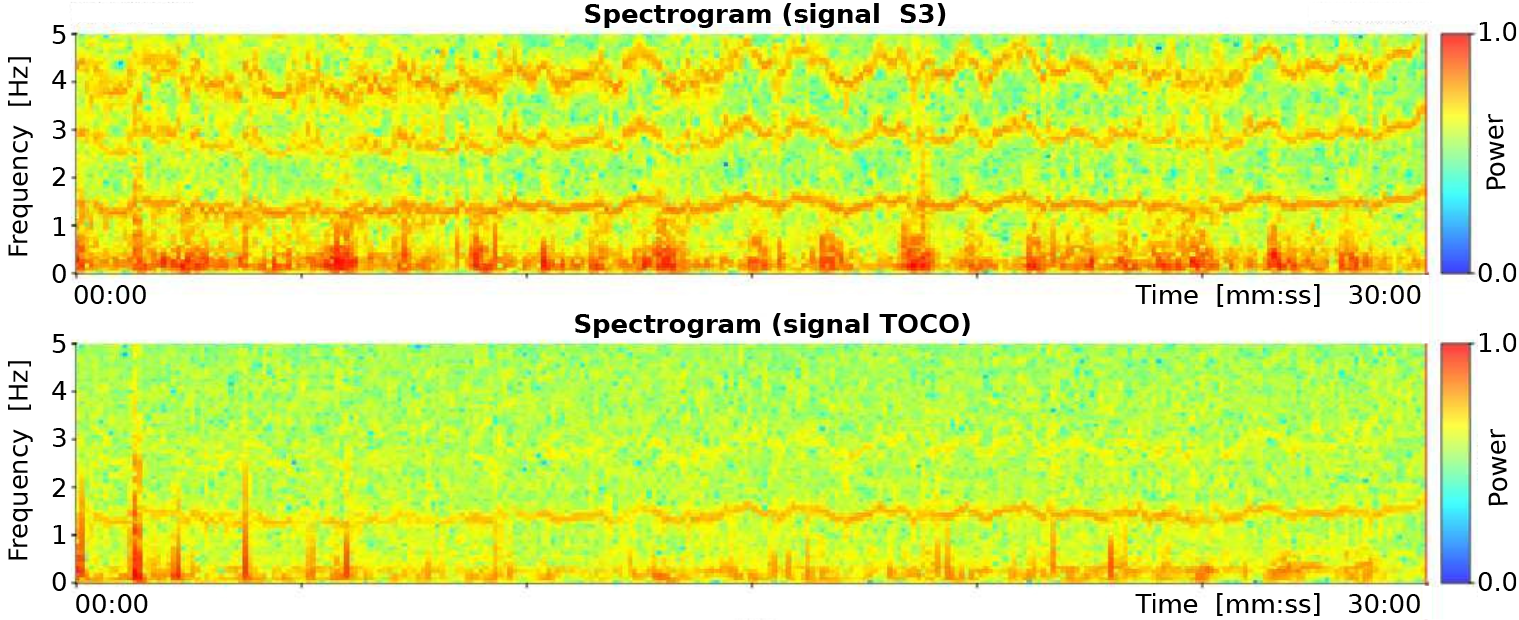
Spectrograms of the record *tpehgt_t011* (*term* nonlabor record, recorded in the 29th week, delivery in the 39th week) of the TPEHGT DS.

*Contraction* and *dummy* intervals of *preterm* and *term* records of the TPEHGT DS were characterized in terms of normalized power spectra. The separation of the records into nonlabor and labor groups, with boundary of three weeks, resulted in 35 *preterm* nonlabor (*contraction* or *dummy*) intervals, 12 *preterm* labor intervals, 41 *term* nonlabor intervals, and 12 *term* labor intervals. Figs 7-9 show the overlaid normalized power spectra of the EHG signals S2, S3, and TOCO signal for *contraction* intervals. Besides the content in the frequency band B0 due to uterine contractions and maternal respiration, the most obvious and strong component in the spectra of *contraction* intervals for *term* nonlabor group of records is activity due the maternal heart in the frequency band B1 (Figs 7, 8, and 9), with higher harmonics in the frequency bands B2 and B3 (Figs 7 and 8). The frequency of the main component in the frequency band B1 ranges from approximately 1.2 Hz (72 bpm) to 1.5 Hz (90 bpm). The frequencies of the peaks in the frequency bands B2 and B3 were additionally verified for a few significant examples. The frequencies were the exact second and third harmonics of the frequency of the peak in the frequency band B1. The presence of the maternal heart activity is expected since the ratio between the ECG activity and EHG bursts is about 20. This activity in the frequency band B1 is actually absent, or weak, in all signals for *preterm* nonlabor or *preterm* labor intervals, it is stronger in all signals for *term* nonlabor intervals, it is especially strong in the TOCO signals (Fig 9), and it is again weak in the EHG signal S3 (Fig 8), or absent in the TOCO signal (Fig 9), for *term* labor intervals. (S1 Fig also shows the overlaid normalized power spectra of the EHG signal S1 for *contraction* intervals.)

**Fig 7.**
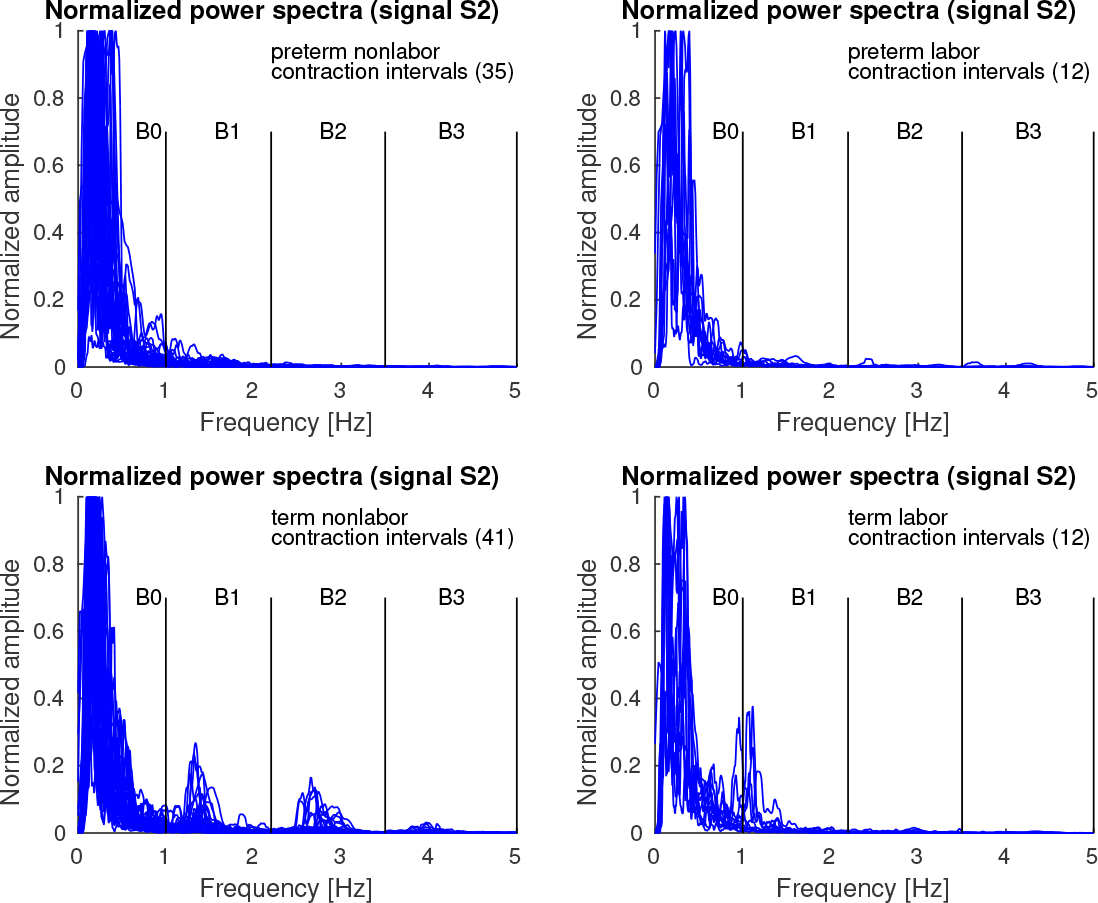
Normalized power spectra of *contraction* intervals of signal S2 of the records of the TPEHGT DS.

**Fig 8.**
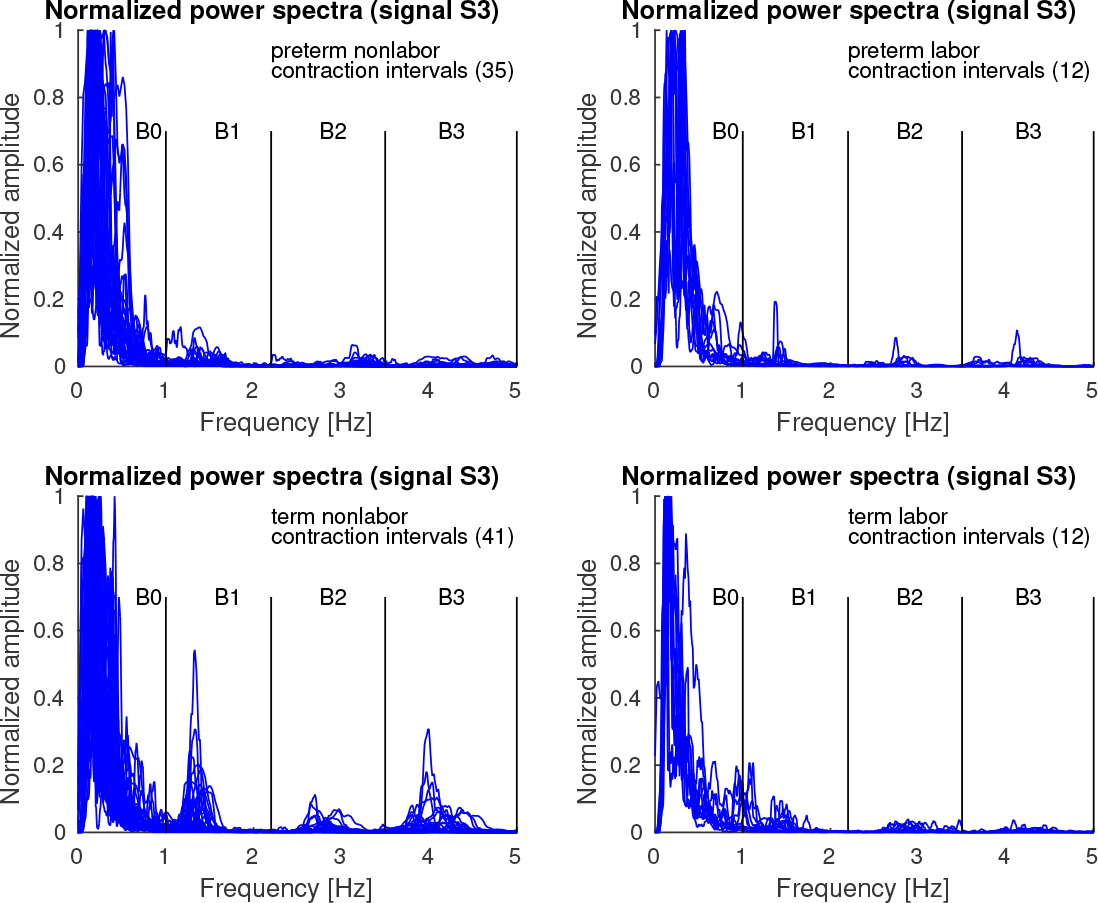
Normalized power spectra of *contraction* intervals of signal S3 of the records of the TPEHGT DS.

**Fig 9.**
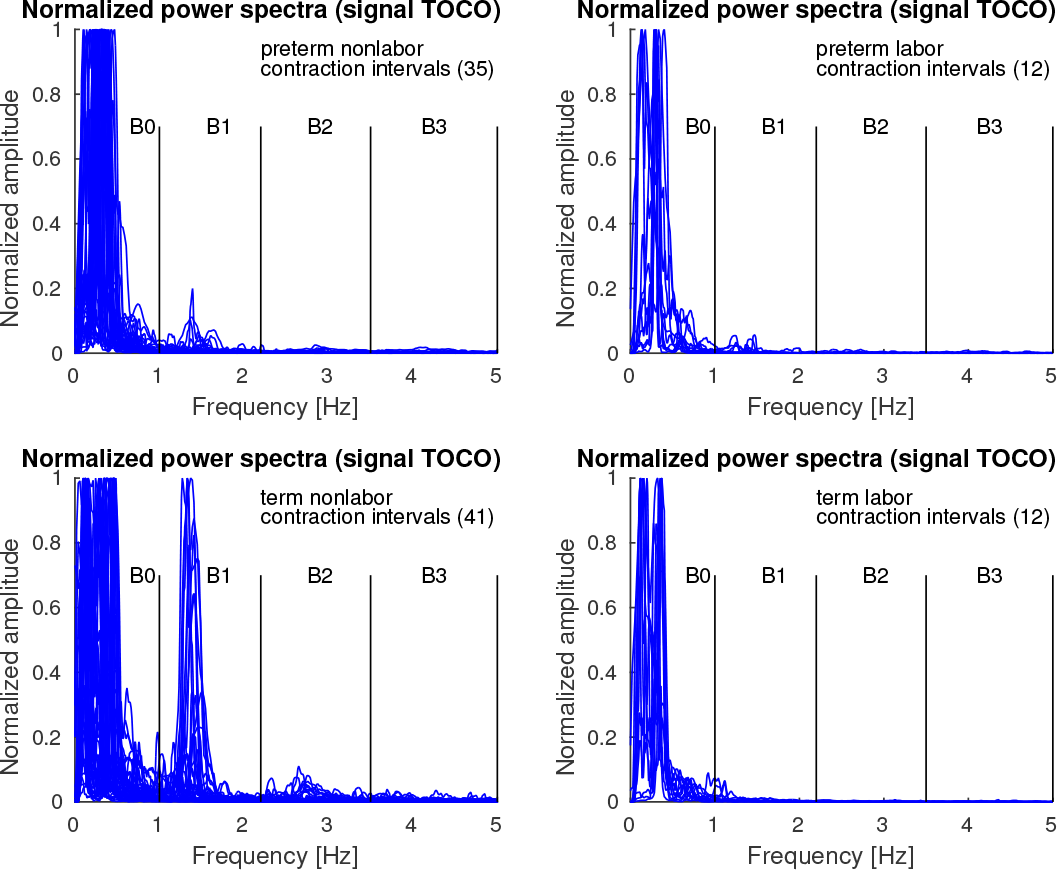
Normalized power spectra of *contraction* intervals of TOCO signal of the records of the TPEHGT DS.

Figs 10 - 12 show the overlaid normalized power spectra of the EHG signals S2, S3, and TOCO signal for *dummy* intervals. Activity due to the maternal heart in the frequency bands B1, B2, and B3, is again present and even stronger. This is reasonable since during *dummy* intervals there are no contractions present. The major component in the frequency band B0 is the maternal respiration component. As for *contraction* intervals, the same pattern regarding presence and intensity of maternal heart activity in the frequency bands B1, B2, and B3, in all signals for *preterm* and *term*, nonlabor and labor, *dummy* intervals may be observed. The activity is again especially strong in the frequency band B1 in *term* nonlabor group for the TOCO signal (Fig 12). (S2 Fig also shows the overlaid normalized power spectra of the EHG signal S1 for *dummy* intervals.)

Fig 13 shows box plots of normalized peak amplitudes, *PA*, in the frequency band B1 of signals S1, S2, S3, and TOCO, for *preterm* and *term*, nonlabor and labor, groups of *contraction* and *dummy* intervals. The box plots reveal that the influence of the maternal heart is the strongest in signal S1 for *dummy* intervals. The maternal heart activity is much stronger for *term contraction* and *dummy* intervals than for *preterm contraction* and *dummy* intervals. The activity does not change much from the *preterm* nonlabor to *preterm* labor group, it is the highest for the *term* nonlabor group, and drops in signals S2, S3, and TOCO for the *term* labor group. The activity in the EHG signals is also stronger for *dummy* intervals than for *contraction* intervals in all four groups. Table 1 summarizes the values of the median of the normalized peak amplitudes, *PA*, in the frequency band B1 for the groups of records, and their relations for signals. *Contraction* intervals, in comparison to *dummy* intervals, show lower median of the *PA* for all EHG signals in all four groups of records, and comparable median of the *PA* for the TOCO signal in all four groups of records. For the EHG signals, this is likely due to contractions yielding higher *PA* in the frequency band B0, and consequently low normalized peak amplitude, *PA*, in the frequency band B1. Maternal heart activity in the frequency band B1 is high for *term* nonlabor and for *term* labor groups, but not for *preterm* nonlabor and *preterm* labor groups for both types of intervals. If the labor is close, or if there is a danger of preterm birth, the activity is low. Moreover, the highest ratios between the medians of the *PA* for *term* nonlabor and *preterm* nonlabor groups, or between the medians of the *PA* for *term* nonlabor and maximum of *preterm* groups, were found in the TOCO signal, and then in the EHG signal S2, for both, *contraction* and *dummy* intervals.

**Fig 10.**
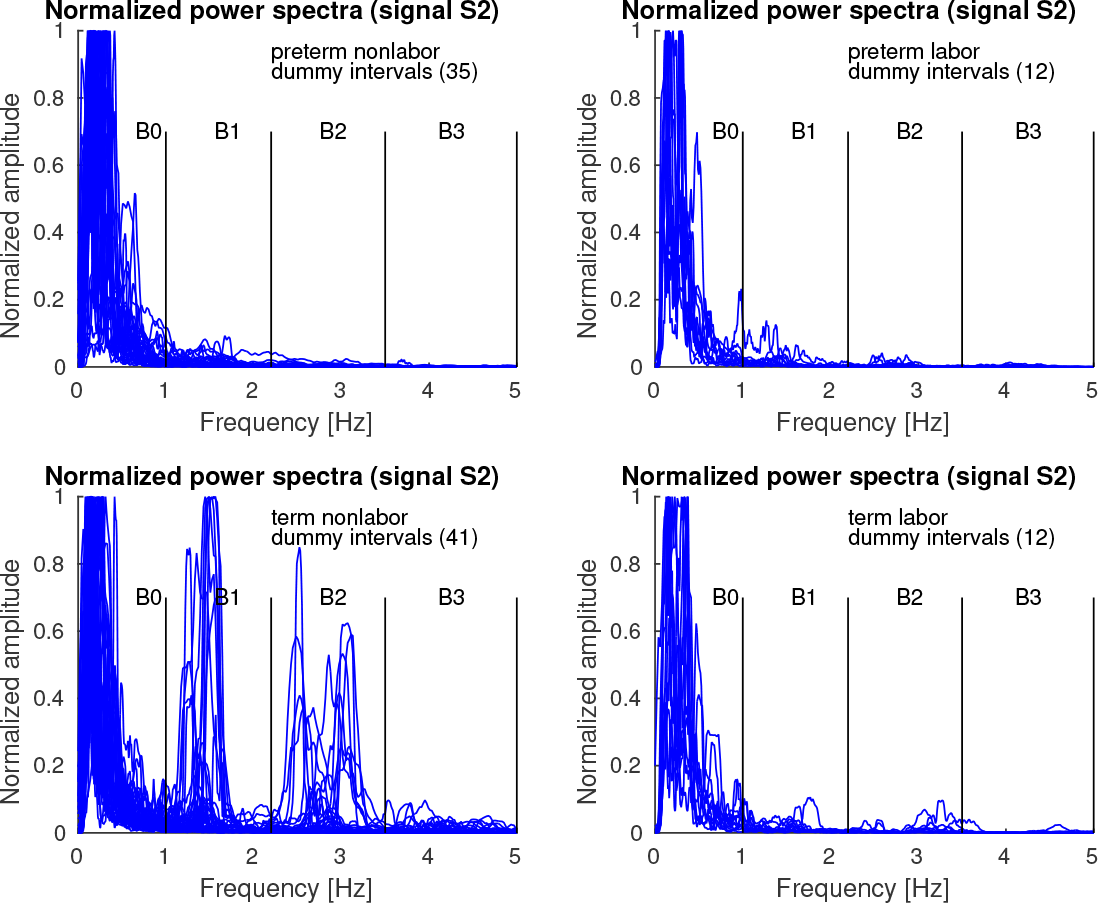
Normalized power spectra of *dummy* intervals of signal S2 of the records of the TPEHGT DS.

**Fig 11.**
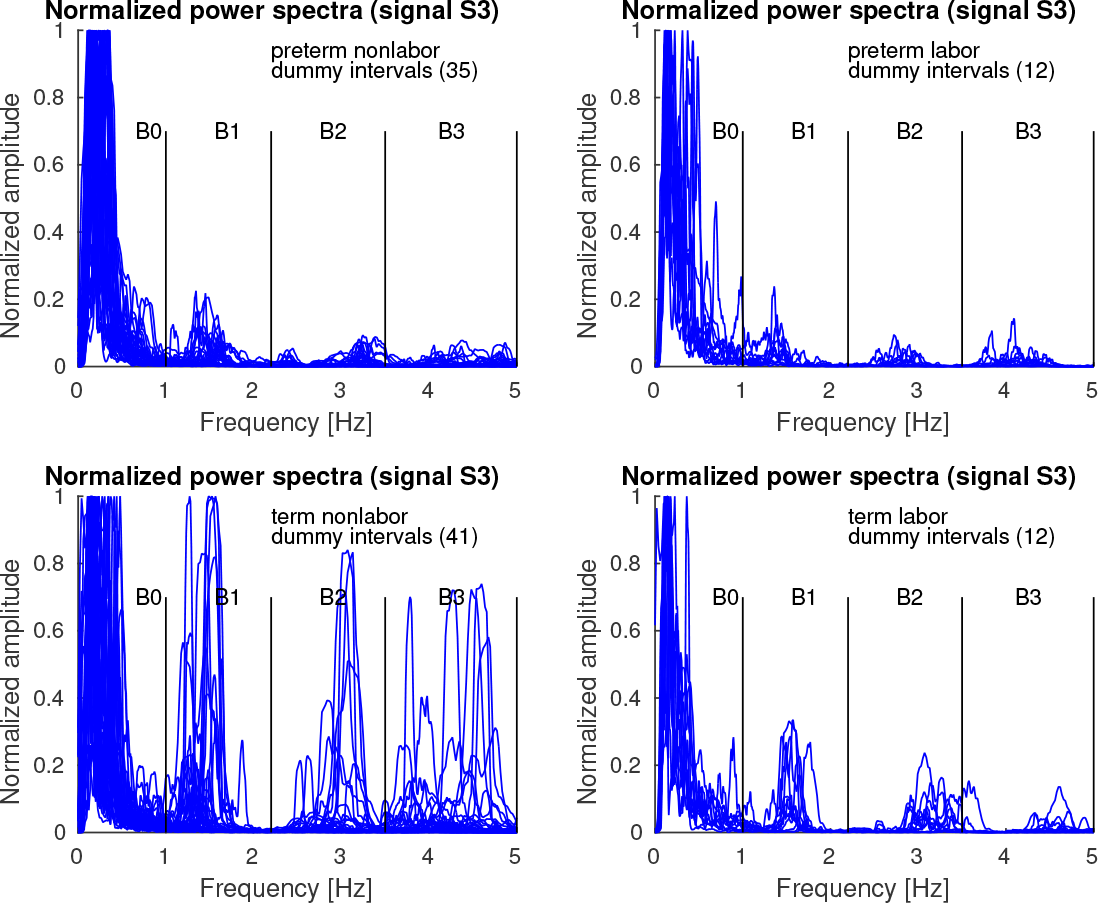
Normalized power spectra of *dummy* intervals of signal S3 of the records of the TPEHGT DS.

**Fig 12.**
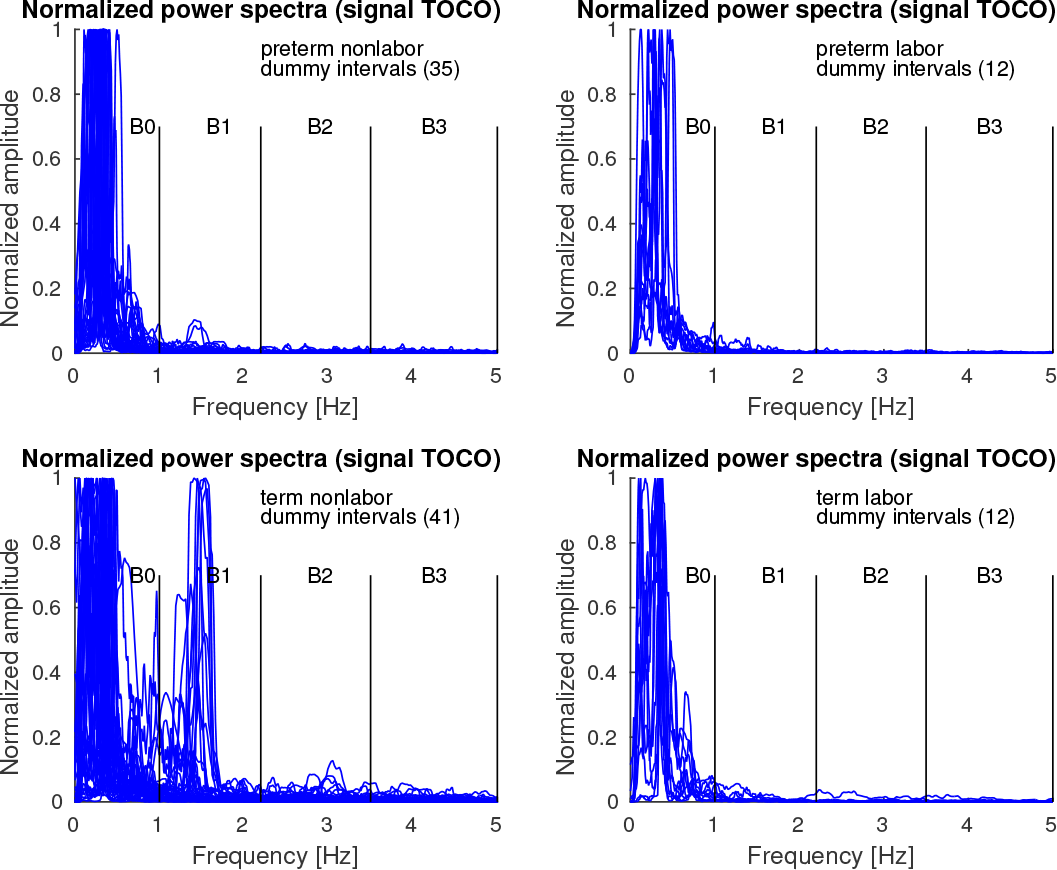
Normalized power spectra of *dummy* intervals of TOCO signal of the records of the TPEHGT DS.

**Fig 13.**
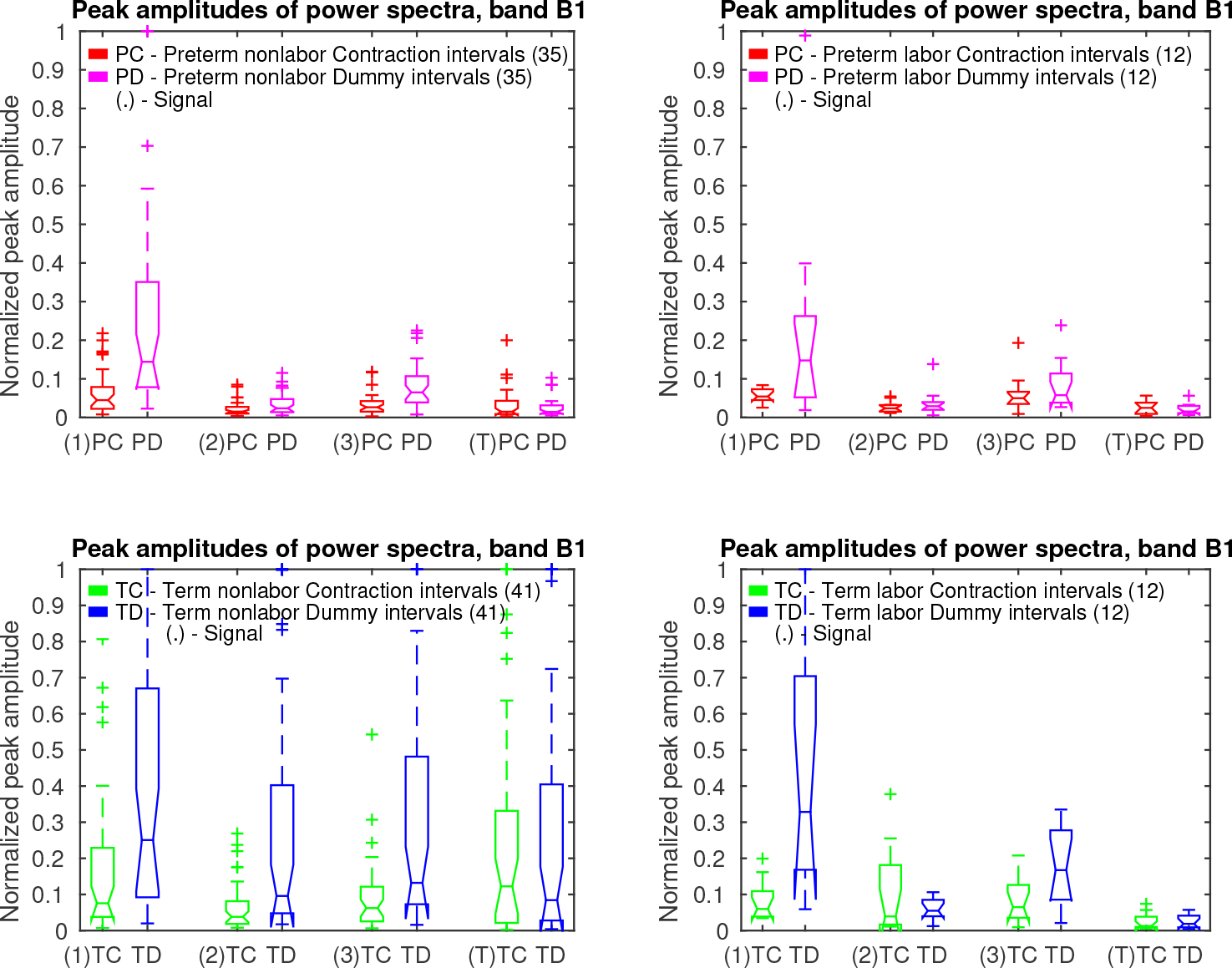
Box plots of normalized peak amplitudes, *PA*, in the frequency band B1 of signals S1, S2, S3, and TOCO, for *preterm* and *term*, nonlabor and labor, groups of *contraction* and *dummy* intervals. The line in the middle of each box is the sample median. The tops and bottoms of each box are the 25th and 75th percentiles of the samples. The wiskers represent the 10th and 90th percentiles. Crosses are outliers. Notches display the variability of the median between samples.

**Table 1.** The values of the median of the normalized peak amplitudes, *PA*, in the frequency band B1 for the groups of records.

| TPEHGT DS |  |  |  |  |  |  |  |  |
| --- | --- | --- | --- | --- | --- | --- | --- | --- |
| Group | Median $PA$, band B1, Signals S1, S2, S3, TOCO | | | | | | | |
|  | Contraction intervals |  |  |  | Dummy intervals |  |  |  |
|  | S1 | S2 | S3 | TOCO | S1 | S2 | S3 | TOCO |
| 35 *preterm* nonlabor intervals | 0.045 | 0.015 | 0.026 | 0.015 | 0.144 | 0.024 | 0.065 | 0.015 |
| 12 *preterm* labor intervals | 0.054 | 0.024 | 0.050 | 0.025 | 0.148 | 0.029 | 0.058 | 0.016 |
| 41 *term* nonlabor intervals | **0.076** | 0.038 | 0.062 | **0.123** | 0.251 | **0.096** | **0.132** | **0.084** |
| 12 *term* labor intervals | 0.060 | **0.039** | **0.065** | 0.013 | **0.328** | 0.055 | 0.167 | 0.019 |
| 41 *term* nonlabor intervals | 0.076 | 0.038 | 0.062 | 0.123 | 0.251 | 0.096 | 0.132 | 0.084 |
| 35 *preterm* nonlabor intervals | 0.045 | 0.015 | 0.026 | 0.015 | 0.144 | 0.024 | 0.065 | 0.015 |
| Average | 0.061 | (0.027) | 0.044 | (0.069) | 0.198 | (0.060) | 0.099 | (0.050) |
| Ratio | 1.689 | 2.533 | 2.385 | 8.200 | 1.743 | 4.000 | 2.031 | 5.600 |
| 41 *term* nonlabor intervals | 0.076 | 0.038 | 0.062 | 0.123 | 0.251 | 0.096 | 0.132 | 0.084 |
| Max (47 *preterm* intervals) | 0.054 | 0.024 | 0.050 | 0.025 | 0.148 | 0.029 | 0.065 | 0.016 |
| Average | 0.065 | (0.031) | 0.056 | (0.074) | 0.200 | (0.063) | 0.099 | (0.050) |
| Ratio | 1.407 | 1.583 | 1.240 | 4.920 | 1.696 | 3.310 | 2.031 | 5.250 |

The highest values of the median of the $PA$ per signal are in bold. Boxed are the highest ratios of the medians for the EHG signals and for TOCO signal. Bracketed are the averages of the medians for the signals with the highest ratios.

Fig 14 shows the overlaid normalized power spectra of the EHG signals S1, S2, S3, and TOCO signal of all 53 *non-pregnant dummy* intervals of the records of the TPEHGT DS. The major component in the frequency band B0 for the signals is the maternal respiration component. The intensity of the maternal heart activity in the frequency bands B1, B2, and B3, for the EHG signals is present, but weak. The pattern of low influence of the maternal heart in the EHG signals of non-pregnant women (Fig 14) is very similar to the pattern for *preterm* nonlabor, *preterm* labor, and *term* labor, *contraction* and *dummy*, groups of pregnant women, but not to the pattern for the *term* nonlabor group of pregnant women (Figs 7, 8, 10, and 11). In addition, there is no obvious or isolated peak in the frequency bands B1, B2, and B3, for the TOCO signal, suggesting that there is no significant mechanical activity present due the influence of the maternal heart. (S3 Fig also shows the normalized power spectra of the signals of the ninth *non-pregnant dummy* interval of the record *tpehgt-n002* from Fig 3. S1 File contains the filtered signals of the interval, while the S2 File contains the normalized power spectra of the filtered signals of the interval.) All these observations regarding *contraction* and *dummy* intervals, and *non-pregnant dummy* intervals, suggest that the uterus responds strongly to maternal heart activity for *term* pregnancies in the nonlabor phase only.

Fig 15 show box plots of normalized peak amplitudes, *PA*, in the frequency band B1 of signals S1, S2, S3, and TOCO, for all *non-pregnant dummy* intervals, and for all *preterm contraction* and *dummy* intervals, versus all *term contraction* and *dummy* intervals. Distributions of *non-pregnant dummy* intervals are quite comparable to distributions of *preterm contraction* and *dummy* intervals. The influence of the maternal heart is the strongest in signal S1. In terms of separability of *preterm* and *term* intervals, distributions of *PA* for *dummy* intervals of pregnant women offer higher separability for each signal.

**Fig 14.**
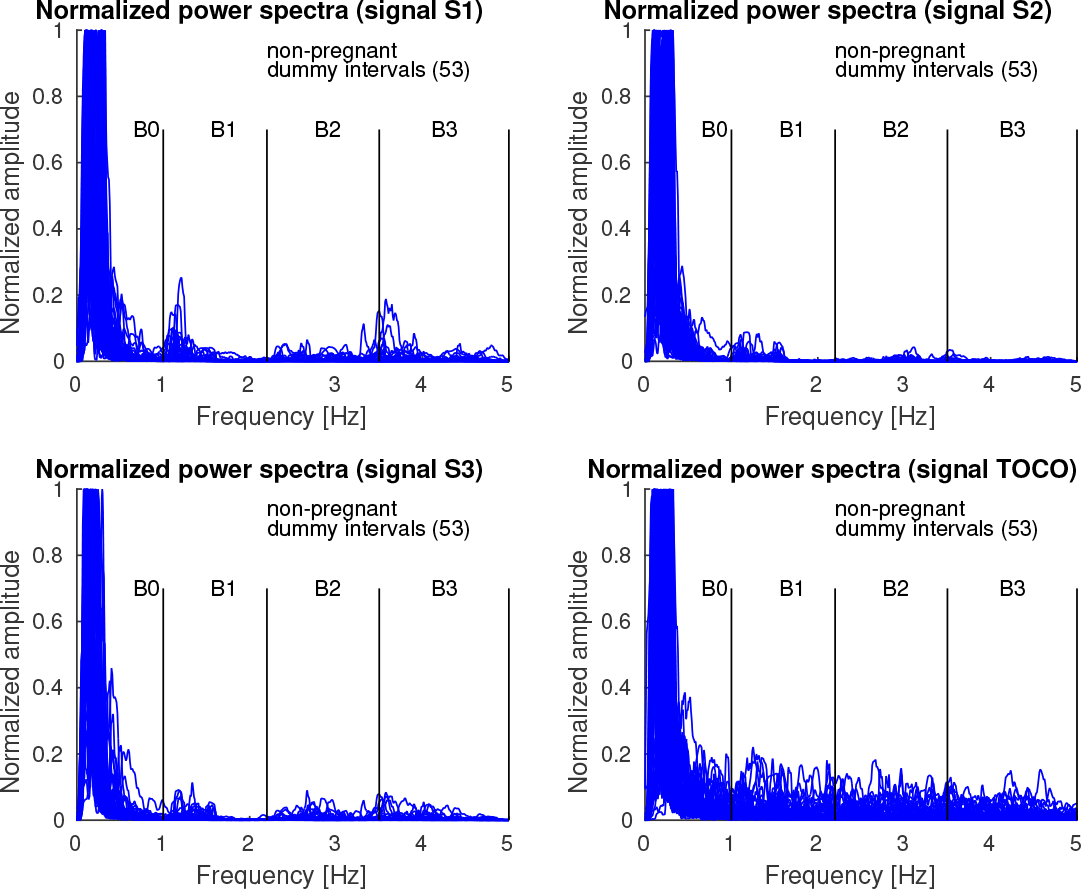
Normalized power spectra of *non-pregnant dummy* intervals of signals S1, S2, S3, and TOCO of the records of the TPEHGT DS.

**Fig 15.**
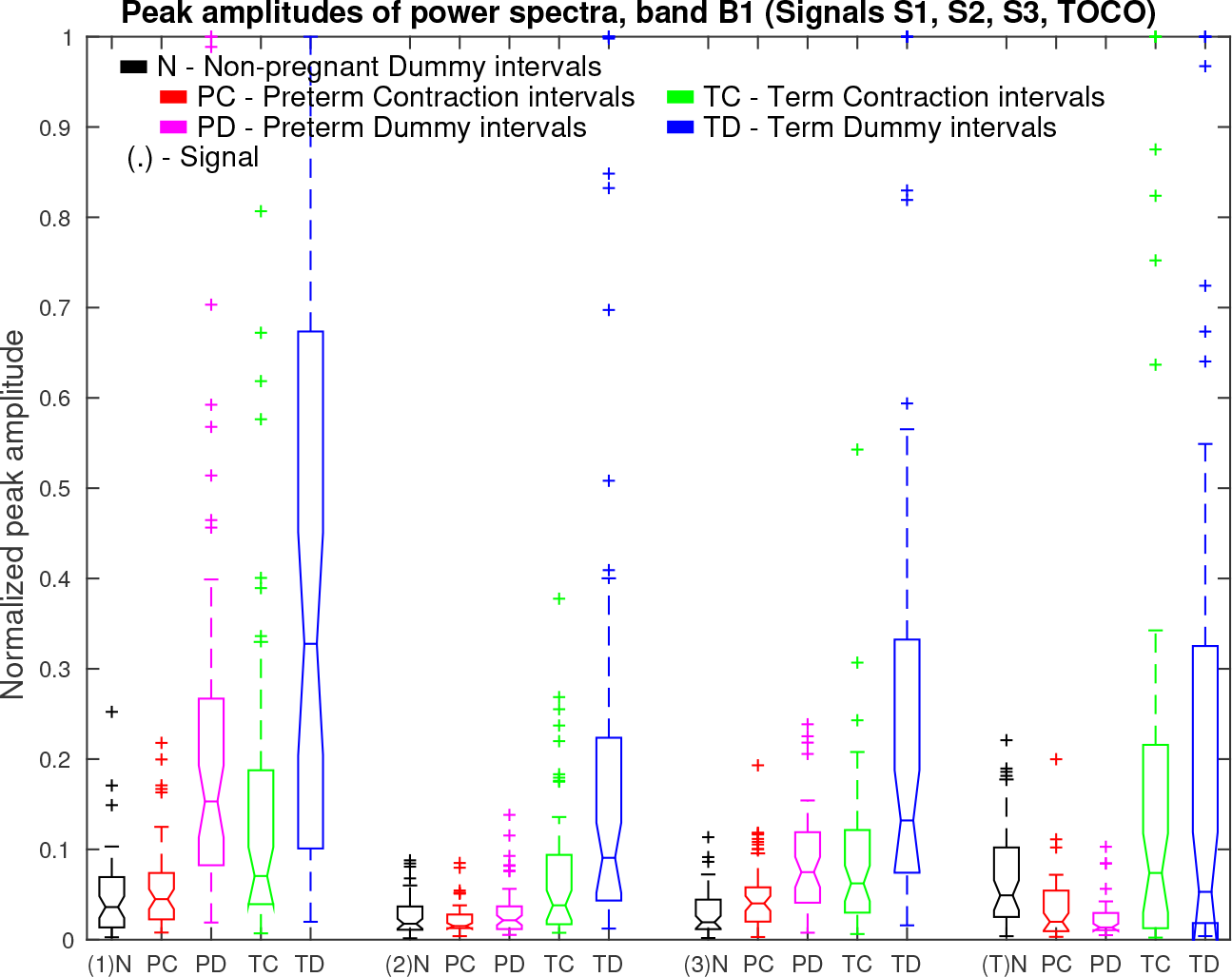
Box plots of normalized peak amplitudes, *PA*, in the frequency band B1 of signals S1, S2, S3, and TOCO, for all *non-pregnant dummy* intervals and for all *preterm* and *term*, *contraction* and *dummy* intervals. Also see caption to Fig 13.

Fig 16 shows box plots of sample entropies, *SE*, in the frequency band B1 of signals S1, S2, S3, and TOCO, for all *non-pregnant dummy* intervals, and for all *preterm contraction* and *dummy* intervals, versus all *term contraction* and *dummy* intervals. Note that the highest values of *SE* for *non-pregnant dummy* intervals are in the TOCO signal, suggesting that the maternal heart activity is barely present, or absent, in the TOCO signal for non-pregnant women. On the other hand, the low valus of *SE* for *preterm* and *term, contraction* and *dummy* intervals in the TOCO signal do suggest the presence of maternal heart activity in the frequency band B1 for pregnant women. These low values of *SE* in the TOCO signal for pregnant women are also lower than the values of *SE* in the EHG signals S1, S2, and S3, suggesting higher regularity of the TOCO signal in the frequency band B1 for pregnant women. Distributions of *SE* for *contraction* and for *dummy* intervals in signals S1, S2, S3, and TOCO of pregnant women do offer separability in the sense of classification of *preterm* versus *term* intervals. Moreover, the distribution of *SE* for *non-pregnant dummy* intervals in the TOCO signal offers very high separability between the *dummy* intervals of non-pregnant women, and *contraction* and *dummy* intervals of pregnant women.

**Fig 16.**
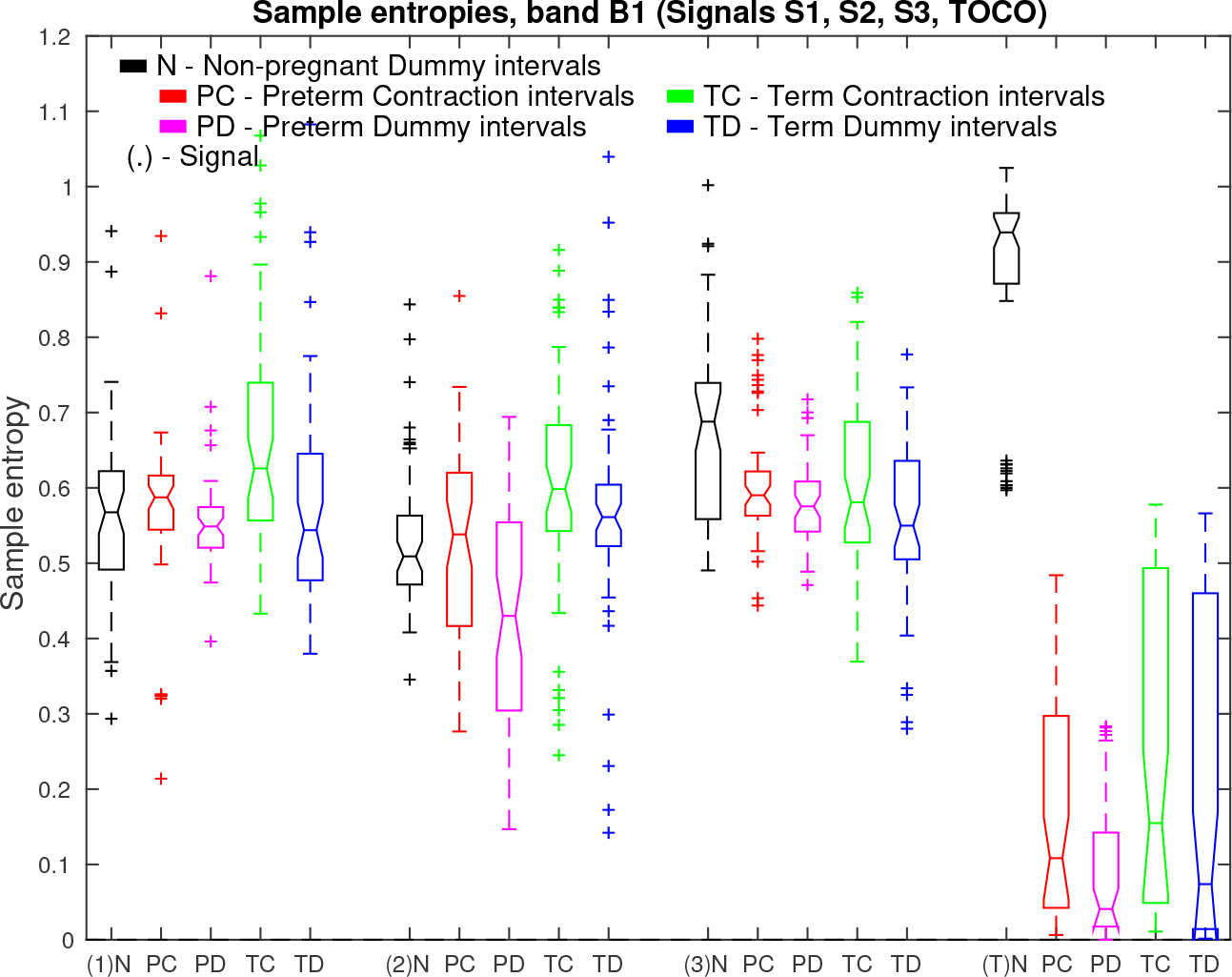
Box plots ot sample entropies, *SE*, in the frequency band B1 ot signals S1, S2, S3, and TOCO, for all *non-pregnant dummy* intervals and for all *preterm* and *term*, *contraction* and *dummy* intervals. Also see caption to Fig 13.

Finally, Fig 17 shows box plots of median frequencies, *MF*, in the frequency band B1 of signals S1, S2, S3, and TOCO, for all *non-pregnant dummy* intervals, and for all *preterm contraction* and *dummy* intervals, versus all *term contraction* and *dummy* intervals.

### Classification of *preterm* and *term* uterine records

#### Feature ranking using the TPEHGT DS

For the classification of *preterm* and *term*, *contraction* or *dummy* intervals of the TPEHGT DS, the sample entropy, *SE*, median frequency of normalized power spectrum, *MF*, and peak amplitude of normalized power spectrum, *PA*, were derived in each of the frequency bands B0, B1, B2, and B3, and for each of the input signals, given *contraction* or *dummy* intervals, resulting in 12 features per input signal per interval. In the majority of cases the maximum power spectrum component, P_max_, needed for normalization of the spectrum was found in the frequency band B0 due to maternal respiration and contractions, therefore *PA* of the frequency band B0 was omitted, resulting in 11 features per signal per interval.

**Fig 17.**
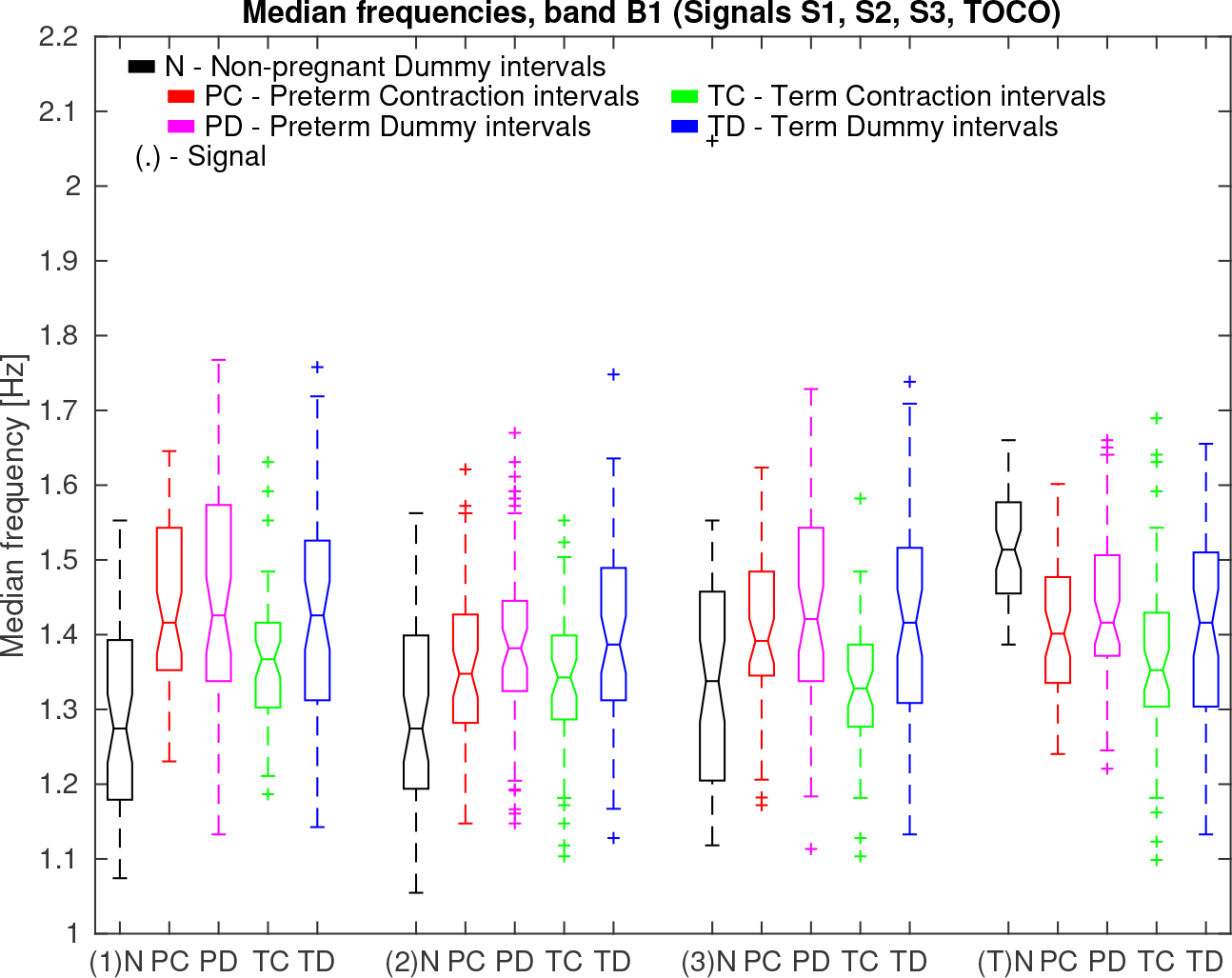
Box plots of median frequencies, *MF*, in the frequency band B1 of signals S1, S2, S3, and TOCO, for all *non-pregnant dummy* intervals and for all *preterm* and *term, contraction* and *dummy* intervals. Also see caption to Fig 13.

Table 2 summarizes the significance of each of the extracted features in each of the frequency bands for signals S2, S3, and TOCO to separate *preterm* and *term*, *contraction* and *dummy* intervals of the TPEHGT DS. The SMOTE technique to equalize the number of samples in two separate classes increased the number of samples in *preterm* minority class from 47 to 53. The values of the two-sample *t*-test, *p*, and the values of the Bhattacaryya criterion, *C*_B_, with its ranks for individual features obtained from *contraction* and *dummy* intervals, are shown. The values of the *C_B_* are also shown in Fig 18 in the form of bar charts. According to the ranks of the features, the most significant individual features for classification appear to be the *PA* features from the frequency bands B1, B2, and B3, especially those from *dummy* intervals. The differences in the ranks, if calculated for each individual signal, and for both types of intervals, using the relative entropy criterion did not differ from the ranks of the Bhattacaryya criterion until the sixth place, in each case. (S4 Fig summarizes the significance of each of the extracted features in each of the frequency bands for signals S2, S3, and TOCO, to also separate labor and nonlabor, *contraction* and *dummy* intervals of the TPEHGT DS. The SMOTE technique increased the number of samples in the labor minority class from 24 to 76.)

**Table 2.** The values of the two-sample *t*-test (*p* values), and of the Bhattacaryya criterion, *C*_B_, with its ranks per group of signals, to separate *preterm* and *term, contraction* and *dummy* intervals of the TPEHGT DS.

| TPEHGT DS |  |  |  |  |  |  |  |  |  |  |  |
| --- | --- | --- | --- | --- | --- | --- | --- | --- | --- | --- | --- |
| Signal | Contraction intervals (53 *preterm* / 53 *term*) |  |  |  |  |  |  |  |  |  |  |
|  | Band B0 |  | Band B1 |  |  | Band B2 |  |  | Band B3 |  |  |
|  | SE | MF | SE | MF | PA | SE | MF | PA | SE | MF | PA |
| S2 ($p$) | 0.138 | 0.008 | 0.004 | 0.271 | $\leq 1^{-3}$ | $\leq 1^{-3}$ | 0.028 | $\leq 1^{-3}$ | $\leq 1^{-3}$ | 0.886 | $\leq 1^{-3}$ |
| S3 ($p$) | 0.139 | $\leq 1^{-3}$ | 0.631 | $\leq 1^{-3}$ | 0.002 | 0.295 | $\leq 1^{-3}$ | 0.335 | 0.554 | $\leq 1^{-3}$ | 0.013 |
| TOCO ($p$) | 0.019 | 0.634 | 0.038 | 0.046 | $\leq 1^{-3}$ | 0.014 | 0.564 | 0.003 | 0.254 | 0.210 | 0.011 |
| S2 ($C_B$) | 0.007 | 0.017 | 0.007 | $\leq 1^{-3}$ | 0.147 | 0.040 | 0.003 | 0.292 | 0.047 | $\leq 1^{-3}$ | 0.072 |
| S3 ($C_B$) | 0.005 | 0.007 | 0.009 | 0.009 | 0.054 | $\leq 1^{-3}$ | 0.015 | 0.009 | 0.003 | 0.028 | 0.085 |
| TOCO ($C_B$) | 0.007 | 0.008 | 0.015 | 0.004 | 0.276 | 0.059 | 0.004 | 0.088 | 0.038 | 0.002 | 0.041 |
| S2 ($C_B$ rank) | 15 | **9** | 16 | 21 | **2** | **7** | 19 | **1** | **6** | 22 | **4** |
| S3 | 17 | 14 | 12 | **11** | **5** | 20 | **10** | 13 | 18 | **8** | **3** |
| S2 ($C_B$ rank) | 14 | **11** | 16 | 21 | **3** | **9** | 19 | **1** | **7** | 22 | **5** |
| TOCO | 15 | 13 | 12 | 17 | **2** | **6** | 18 | **4** | **10** | 20 | **8** |

| Signal | Dummy intervals (53 *preterm* / 53 *term*) |  |  |  |  |  |  |  |  |  |  |
| --- | --- | --- | --- | --- | --- | --- | --- | --- | --- | --- | --- |
|  | Band B0 |  | Band B1 |  |  | Band B2 |  |  | Band B3 |  |  |
|  | SE | MF | SE | MF | PA | SE | MF | PA | SE | MF | PA |
| S2 ($p$) | 0.004 | 0.066 | $\leq 1^{-3}$ | 0.785 | $\leq 1^{-3}$ | $\leq 1^{-3}$ | $\leq 1^{-3}$ | $\leq 1^{-3}$ | $\leq 1^{-3}$ | 0.632 | $\leq 1^{-3}$ |
| S3 ($p$) | 0.952 | 0.332 | 0.128 | 0.696 | $\leq 1^{-3}$ | 0.866 | 0.022 | 0.002 | 0.274 | 0.016 | $\leq 1^{-3}$ |
| TOCO ($p$) | 0.013 | 0.460 | $\leq 1^{-3}$ | 0.227 | $\leq 1^{-3}$ | 0.068 | 0.554 | $\leq 1^{-3}$ | 0.118 | 0.193 | $\leq 1^{-3}$ |
| S2 ($C_B$) | 0.005 | 0.003 | 0.016 | $\leq 1^{-3}$ | 0.326 | 0.048 | 0.039 | 0.473 | 0.064 | 0.018 | 0.188 |
| S3 ($C_B$) | $\leq 1^{-3}$ | 0.003 | 0.035 | $\leq 1^{-3}$ | 0.178 | 0.016 | 0.006 | 0.226 | 0.016 | 0.009 | 0.232 |
| TOCO ($C_B$) | 0.006 | 0.006 | 0.055 | 0.004 | 0.399 | 0.009 | 0.005 | 0.124 | 0.022 | $\leq 1^{-3}$ | 0.048 |
| S2 ($C_B$ rank) | 17 | 18 | 12 | 22 | **2** | **8** | **9** | **1** | **7** | **11** | **5** |
| S3 | 20 | 19 | **10** | 21 | **6** | 14 | 16 | **4** | 13 | 15 | **3** |
| S2 ($C_B$ rank) | 18 | 20 | 13 | 22 | **3** | **8** | **10** | **1** | **6** | 12 | **4** |
| TOCO | 16 | 15 | **7** | 19 | **2** | 14 | 17 | **5** | **11** | 21 | **9** |

Those $p$ values $\leq 1^{-3}$ and ranks of the first 11 features according to $C_B$ per group of signals are in bold.

**Fig 18.**
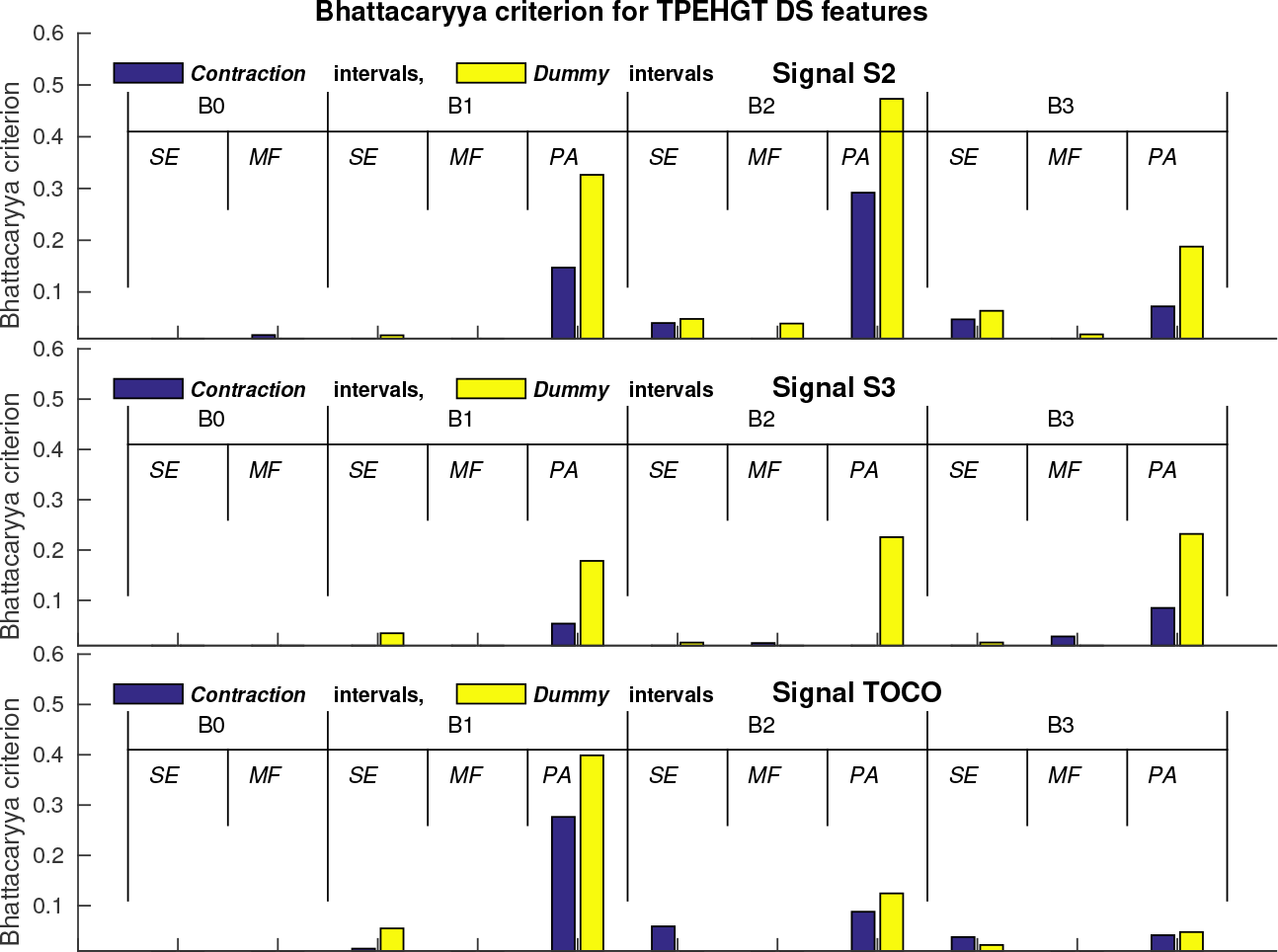
Values of the Bhattacaryya criterion, *C*_B_, for individual features to separate *preterm* and *term*, *contraction* and *dummy* intervals of the TPEHGT DS.

#### Feature selection and classification of *contraction* and *dummy* intervals of the TPEHGT DS

Inspired by the high individual separation ability of the normalized peak amplitude of the power spectrum, *PA*, in the frequency band B1, *PA*_B1_, for EHG signal S2, and for TOCO signal, reflecting the influence of maternal heart activity (Tables 1, 2, and Fig 18), we first tested its individual classification performance. Table 3 shows the classification performance results obtained for *preterm* and *term*, nonlabor, and for *preterm* and *term*, *contraction* and *dummy*, intervals of the TPEHGT DS. For *preterm* and *term* nonlabor groups, if using both signals, *dummy* intervals showed higher classification accuracy, i.e., *CA*=76.83%, than *contraction* intervals. If considering all *preterm* and *term* intervals, *contraction* intervals yielded higher performance, i.e., *CA*=74.53%.

**Table 3.** Classification performance results obtained for *preterm* and *term*, nonlabor, and for *preterm* and *term*, *contraction* and *dummy*, intervals of the TPEHGT DS.

| TPEHGT DS |  |  |  |  |  |  |  |  |
| --- | --- | --- | --- | --- | --- | --- | --- | --- |
| QDA ($PA_{B1}$)<br>10-folds | 41 *preterm* nonlabor / 41 *term* nonlabor | | | | | | | |
|  | *Contraction* intervals |  |  |  | *Dummy* intervals |  |  |  |
| [%] | *Se* | *Sp* | *CA* | *AUC* | *Se* | *Sp* | *CA* | *AUC* |
| Signal S2 | 90 | 39 | 64.63 | 65.66 | 95 | 54 | **74.39** | **78.42** |
| Signal TOCO | 95 | 51 | 73.49 | 76.40 | 93 | 51 | 71.95 | 76.43 |
| Signals S2 and TOCO | 93 | 59 | 75.61 | 77.71 | 90 | 63 | **76.83** | **82.70** |
| QDA ($PA_{B1}$)<br>10-folds | 53 *preterm* / 53 *term* | | | | | | | |
|  | *Contraction* intervals |  |  |  | *Dummy* intervals |  |  |  |
| [%] | *Se* | *Sp* | *CA* | *AUC* | *Se* | *Sp* | *CA* | *AUC* |
| Signal S2 | 92 | 40 | 66.04 | 65.42 | 96 | 43 | **69.81** | **75.30** |
| Signal TOCO | 96 | 42 | 68.87 | 70.64 | 94 | 40 | 66.98 | 72.61 |
| Signals S2 and TOCO | 94 | 55 | **74.53** | **76.82** | 92 | 49 | 70.75 | 79.98 |

The highest *CA* and *AUC* per combination of signals and intervals are in bold.

To further investigate possibilities for the efficient classification between *preterm* and *term*, *contraction* and *dummy* intervals, and between labor and nonlabor, *contraction* and *dummy* intervals of the records of the TPEHGT DS, i.e., four classification tasks, the EHG signals S2 and S3 were used due to their orthogonality, and the TOCO signal carrying mechanical information of maternal heart activity. The EHG signal S3 was chosen, since it was recorded from the lower, stretchable part of the uterus, and closer to the cervix (the bottom electrodes, Fig 1). In order to test the classification ability of the TOCO signal, the four classification tasks were performed in two variants, using the EHG signals S2 and S3, and using the EHG signal S2 and TOCO signal.

We encountered a relatively large number of potential features that can be used for classification, i.e., 11 features per signal per interval. Using two signals yields 22 features per interval. To select the relevant features, the SFS method was used. Fig 19 shows the average values of the misclassification errors, MCE, obtained on the training sets of the TPEHGT DS for the four classification tasks with two variants.

Table 4 shows the selected features with their ranks for the variety of classification tasks after using the frequency-based feature-aggregation and feature-selection procedure. The best individual features may not have a high rank, or may not even be selected. Among the four classification tasks with two variants, the most frequently selected feature, and with the highest rank (three times the first place and twice the second) appear to be the sample entropy of signal S2 in the frequency band B2, i.e., the presence or absence of the second harmonic component of the maternal heart rate. The second most selected feature is the median frequency of signal S2 in the frequency band B2 (twice the first place and three times the third). The third most selected feature is the median frequency of signal S2 in the frequency band B0 (once the first place and twice the second). The fourth feature is the sample entropy of signal S2 in the frequency band B3, i.e., the presence or absence of the third harmonic component of the maternal heart rate (twice the second place and once the third). Regarding the TOCO signal, sample entropies also played an important role. Sample entropies were selected in the frequency band B0 (once the first place and once the second), in the frequency band B1 (once the third place), and in the frequency band B2 (once the second place).

**Fig 19.**
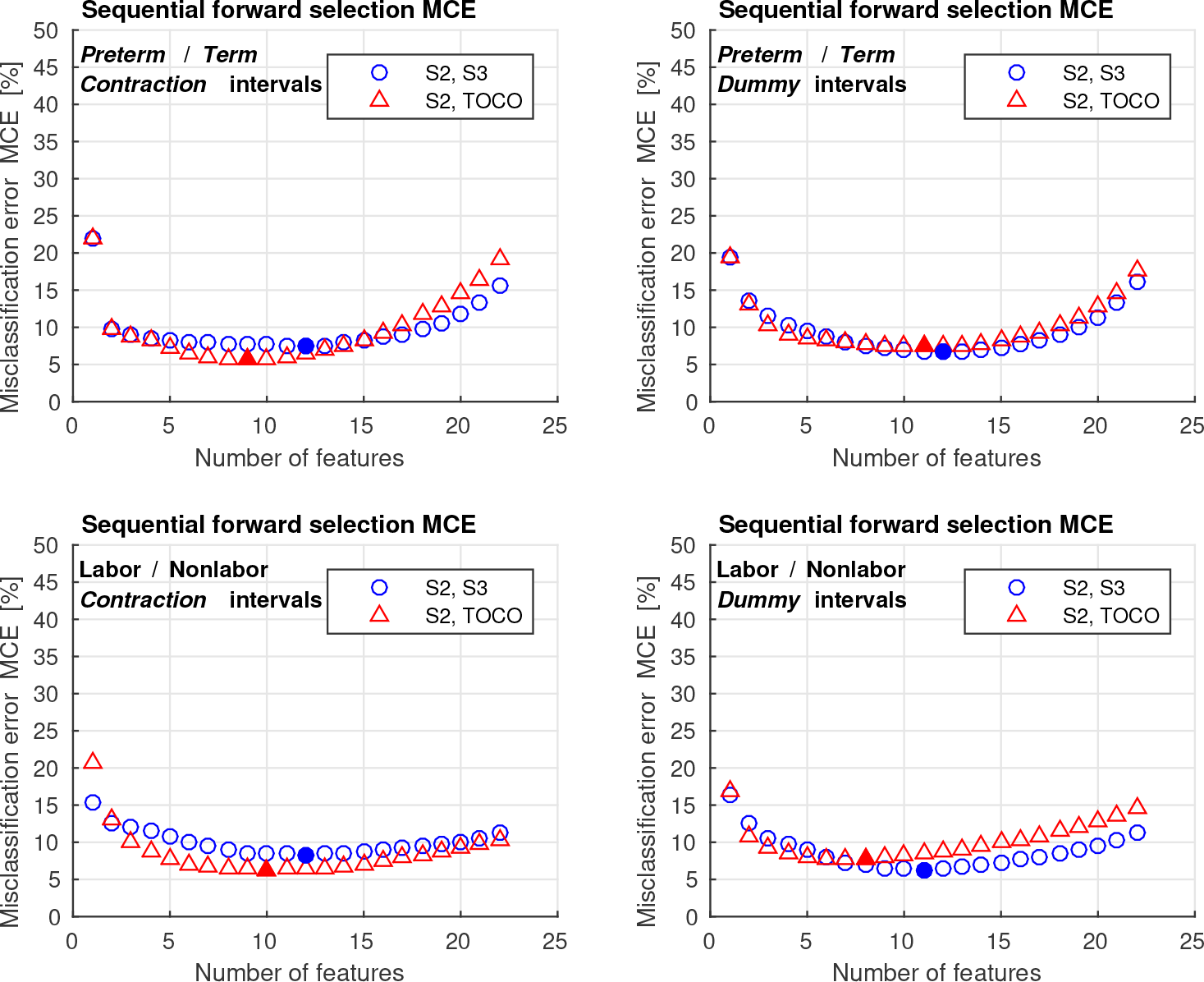
Average values of the misclassification errors, MCE, obtained on the training sets of the TPEHGT DS using the SFS method.

Table 5 shows the classification performance results obtained for *preterm* and *term*, *contraction* and *dummy* intervals, and for the labor and nonlabor, *contraction* and *dummy* intervals of the records of the TPEHGT DS. Basically, our goal here was to evaluate different aspects like: selection of intervals (*contraction* or *dummy*), selection of signals (EHG, or EHG and TOCO), and their influence on the classification capability.

Tables 5A and 5B summarize classification performances, considering the classification ability of signals S2 and S3, or signals S2 and TOCO, to classify *preterm* and *term, contraction* and *dummy* intervals. *Dummy* intervals provided higher classification accuracies than *contraction* intervals for signals S2 and S3, *CA*=88.79% versus *CA*=88.68%, and for signals S2 and TOCO, *CA*=91.51% versus *CA*=90.57%. Moreover, for *dummy* intervals, the EHG signal S2 in combination with the TOCO signal provided higher classification accuracy, *CA*=91.51%, than in combination with the EHG signal S3, *CA*=88.79%.

Another important aspect is the classification ability of signals S2 and S3, or signals S2 and TOCO, for the task of classifying between labor and nonlabor, *contraction* and *dummy* intervals. Tables 5C and 5D summarize these classification performances. Again, *dummy* intervals provided higher classification accuracy than *contraction* intervals for signals S2 and S3, *CA*=93.42% versus *CA*=88.82%, but performance was equal for signals S2 and TOCO, for both types of intervals, *CA*=92.11%. This time, for *dummy* intervals, the combination of the EHG signals S2 and S3 provided higher classification accuracy, *CA*=93.42%, than the combination of the EHG signal S2 and TOCO signal, *CA*=92.11%.

**Table 4.**
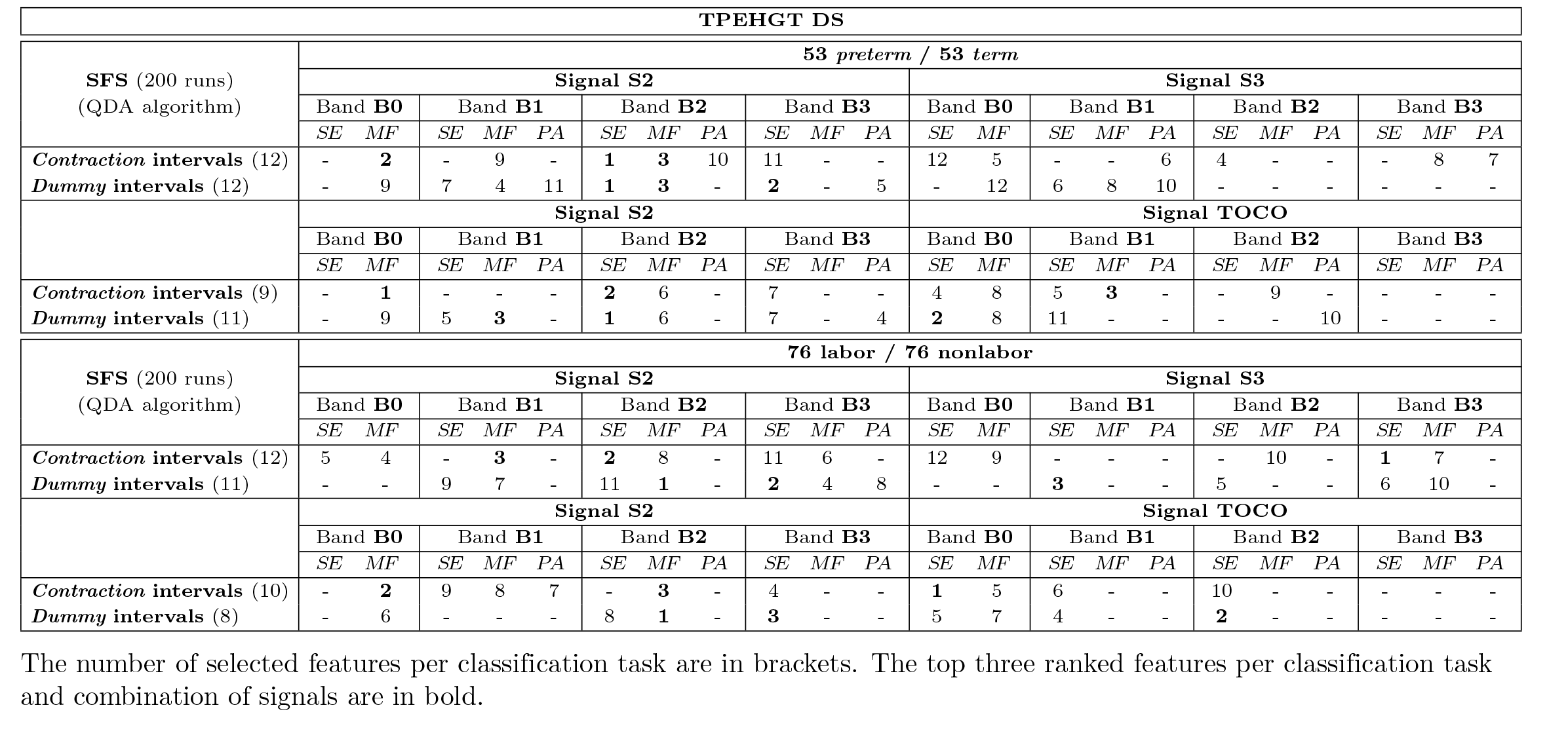
Selected features and their ranks to classify between *preterm* and *term*, and labor and nonlabor, *contraction* and *dummy* intervals of the TPEHGT DS.

**Table 5.** Classification performance results obtained for *preterm* and *term, contraction* and *dummy* intervals, and for nonlabor and labor, *contraction* and *dummy* intervals of the TPEHGT DS.

| TPEHGT DS |  |  |  |  |  |  |  |  |  |  |  |  |  |  |  |  |
| --- | --- | --- | --- | --- | --- | --- | --- | --- | --- | --- | --- | --- | --- | --- | --- | --- |
| 10-folds | A) Signals S2, S3 (53 *preterm* / 53 *term*) |  |  |  |  |  |  |  | B) Signals S2, TOCO (53 *preterm* / 53 *term*) |  |  |  |  |  |  |  |
|  | *Contraction intervals* |  |  |  | *Dummy intervals* |  |  |  | *Contraction intervals* |  |  |  | *Dummy intervals* |  |  |  |
|  | *Se* | *Sp* | *CA* | *AUC* | *Se* | *Sp* | *CA* | *AUC* | *Se* | *Sp* | *CA* | *AUC* | *Se* | *Sp* | *CA* | *AUC* |
| QDA<sub>(SFS)</sub> (12,12,9,11) | 89 | 89 | 88.68 | 94.04 | 87 | 91 | **88.79** | **95.85** | 89 | 92 | 90.57 | **95.65** | 91 | 92 | **91.51** | 95.56 |
| 10-folds | C) Signals S2, S3 (76 *labor* / 76 *nonlabor*) |  |  |  |  |  |  |  | D) Signals S2, TOCO (76 *labor* / 76 *nonlabor*) |  |  |  |  |  |  |  |
|  | *Contraction intervals* |  |  |  | *Dummy intervals* |  |  |  | *Contraction intervals* |  |  |  | *Dummy intervals* |  |  |  |
|  | *Se* | *Sp* | *CA* | *AUC* | *Se* | *Sp* | *CA* | *AUC* | *Se* | *Sp* | *CA* | *AUC* | *Se* | *Sp* | *CA* | *AUC* |
| QDA<sub>(SFS)</sub> (12,11,10,8) | 83 | 95 | 88.82 | 87.79 | 88 | 99 | **93.42** | **95.02** | 87 | 97 | **92.11** | **96.14** | 87 | 97 | **92.11** | 94.61 |

The number of features per type of interval are in brackets. The highest CA and AUC per combination of intervals are in bold.

Yet another very important aspect is differentiating between records of non-pregnant and pregnant women. Classifying between *non-pregnant dummy* intervals of the records of non-pregnant women and *contraction* and *dummy* intervals of the records of pregnant women could reveal the evidence that the electro-mechanical influence of the maternal heart is significantly higher (and measurable through the EHG and TOCO signals) during pregnancy.

Table 6 shows the selected features with their ranks for the task of classifying between *non-pregnant* dummy intervals, and *preterm* and *term, contraction* and *dummy* intervals using the SFS method and frequency-based feature-aggregation and feature-selection procedure. The numbers of selected features is low for each of the four classification tasks with two variants. In all cases sample entropies from different frequency bands were selected. If using signals S2 and S3, *contraction* intervals needed larger numbers of features (six, or five, features for *non-pregnant dummy* versus *preterm*, or *term*, respectively) than *dummy* intervals (three, or two, features for *non-pregnant dummy* versus *preterm*, or *term*, respectively). If using signals S2 and TOCO, only one feature, sample entropy from the frequency band B2, or B3, of the TOCO signal, i.e., presence or absence of the second or third harmonic of the maternal heart rate, was selected for any interval type and for any classification task.

**Table 6.**
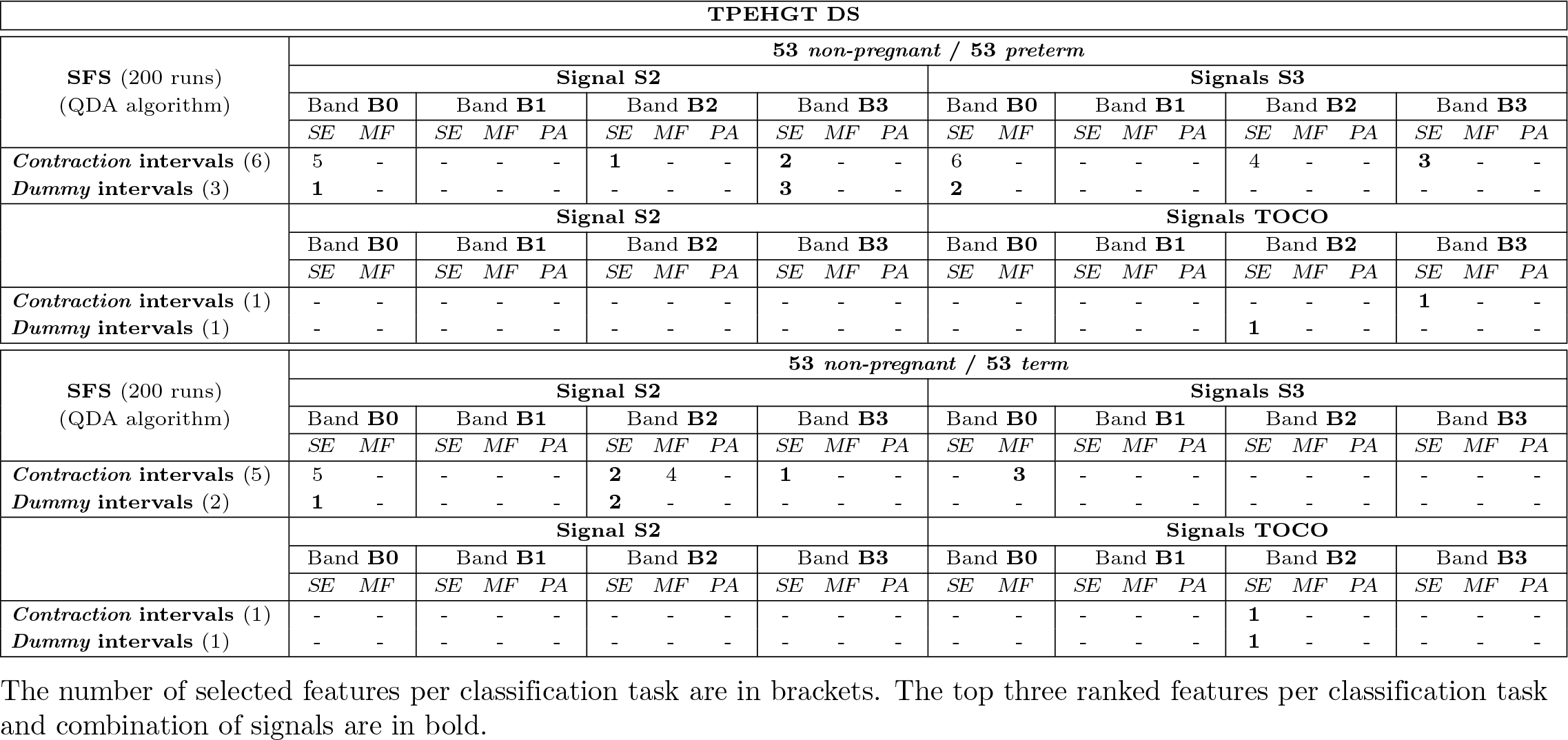
Selected features and their ranks to classify between *non-pregnant dummy* intervals, and *preterm* and *term*, *contraction* and *dummy* intervals of the TPEHGT DS.

Table 7 summarizes classification performance results for classifying *non-pregnant* dummy intervals, and *preterm* and *term, contraction* and *dummy* intervals. The only classification task where the classification accuracy was less than 100% was classifying between *non-pregnant dummy* intervals and *preterm contraction* intervals, and using the EHG signals S2 and S3, *CA* =99.06%. This result suggests that characteristics of *dummy* intervals of the records of non-pregnant women are much different from the characteristics of *contraction* and *dummy* intervals of the records of pregnant women. In addition, more features were needed for classification between *non-pregnant dummy* 27/50intervals and *preterm contraction* or *dummy* intervals than for the classification between *non-pregnant dummy* intervals and *term contraction* or *dummy* intervals, suggesting that the characteristics of the records of non-pregnant women are more similar to the characteristics of the records with *preterm* delivery than to the characteristics of the records with *term* delivery for pregnant women.

**Table 7.** Classification performance results obtained for *non-pregnant dummy* intervals versus *preterm* and *term, contraction* and *dummy* intervals of the TPEHGT DS.

| TPEHGT DS |  |  |  |  |  |  |  |  |  |  |  |  |  |  |  |  |
| --- | --- | --- | --- | --- | --- | --- | --- | --- | --- | --- | --- | --- | --- | --- | --- | --- |
| 10-folds | Signals S2, S3 (53 *non-pregnant* / 53 *preterm*) |  |  |  |  |  |  |  | Signals S2, TOCO (53 *non-pregnant* / 53 *preterm*) |  |  |  |  |  |  |  |
|  | *Contraction intervals* |  |  |  | *Dummy intervals* |  |  |  | *Contraction intervals* |  |  |  | *Dummy intervals* |  |  |  |
|  | Se | Sp | CA | AUC | Se | Sp | CA | AUC | Se | Sp | CA | AUC | Se | Sp | CA | AUC |
| QDA<sub>(SFS)</sub> (6,3,1,1) | 100 | 98 | 99.06 | 99.78 | 100 | 100 | 100 | 100 | 100 | 100 | 100 | 100 | 100 | 100 | 100 | 100 |
| 10-folds | Signals S2, S3 (53 *non-pregnant* / 53 *term*) |  |  |  |  |  |  |  | Signals S2, TOCO (53 *non-pregnant* / 53 *term*) |  |  |  |  |  |  |  |
|  | *Contraction intervals* |  |  |  | *Dummy intervals* |  |  |  | *Contraction intervals* |  |  |  | *Dummy intervals* |  |  |  |
|  | Se | Sp | CA | AUC | Se | Sp | CA | AUC | Se | Sp | CA | AUC | Se | Sp | CA | AUC |
| QDA<sub>(SFS)</sub> (5,2,1,1) | 100 | 100 | 100 | 100 | 100 | 100 | 100 | 100 | 100 | 100 | 100 | 100 | 100 | 100 | 100 | 100 |

The number of features per type of interval are in brackets. The highest CA and AUC per combination of intervals are in bold.

#### Feature ranking using the TPEHG DB

To further verify the influence of the maternal ECG on the uterus, and to further verify the classification performance of the proposed method for predicting preterm birth (Fig 4), the method was tested using publicly available TPEHG DB. The records of the TPEHG DB do not contain the TOCO signal, therefore the influence of the maternal ECG on the uterus could only be verified in the electrical sense. Moreover, there are no annotated *contraction* (or *dummy*) intervals in the TPEHG DB. Thus, the input data intervals of the records were the entire EHG records with a duration of 30 min, which are actually composed from sequences of *contraction* and non-contraction, *dummy*, intervals.

To verify the influence of the maternal ECG on the uterus in the electrical sense, we visualized a large number of records of the TPEHG DB recorded *early* and *later* in terms of spectrograms. Visual examination of the spectrograms confirmed the pattern of strong presence of the maternal heart rate for *term* records, but not for *preterm* records, being recorded *early* or *later.* S5 Fig and S6 Fig show the EHG signals S2 and S3, and the spectrograms of EHG signals S2 and S3, of a *preterm* record (delivery in 30th week) and of a *term* record (delivery in 39th week). Both records were recorded *early*, in the 25th week of pregnancy. The spectrograms of *term* record show strong influence of the maternal heart rate, while the influence is weak, or barely present, for *preterm* record.

For the classification of the entire *preterm* and *term* EHG records, the sample entropy, *SE*, median frequency, *MF*, and peak amplitude, *PA*, of the normalized power spectrum were derived, in each of the frequency bands B0, B1, B2, and B3, and for each of the EHG signals, S1, S2, and S3, of the database. Due to the normalization of each power spectrum, the *PA* from the frequency band B0 was omitted, resulting in 11 features per signal per record. The ADASYN technique, used in order to balance the representation of data distribution in two separate classes, increased the number of samples in *preterm* minority class for *early* records form 19 to 140, and for *all* records from 38 to 256. Table 8 summarizes the significance of each of the extracted features in each frequency band for signals S1, S2, and S3 to separate *preterm* and *term, early* and *all*, records of the TPEHG DB. The values of the two-sample *t*-test, *p*, and the values of the Bhattacaryya criterion, *C*_B_, with its ranks for individual features obtained from *early* and *all* EHG records, are shown. The values of the *C*_B_ for *all* EHG records are also shown in Fig 20 in the form of bar charts. According to the ranks of the features, the most significant individual features for classification between *preterm* and *term* records of the TPEHG DB again appear to be the *PA* features from the frequency bands B1, B2 and B3. The highest separability between *preterm* and *term* records for the *PA* features in the frequency bands B1, B2, and B3 confirms strong influence of the maternal ECG on the uterus. The differences in the ranks, if calculated for each individual signal, for *early* records or for *all* records, using the relative entropy criterion, did not differ from the ranks of the Bhattacaryya criterion until the fifth place in each case.

#### Feature selection and classification of *preterm* and *term* EHG records of the TPEHG DB

The SFS method was employed to select the relevant features. Due to their orthogonality, the EHG signals S2 and S3 were used to classify between *early preterm* and *term* records, providing 22 candidate features. However, for the task of classifying *all* records of the database, a much lower minimum of the misclassification error function (and consequently higher classification accuracy) was obtained if using all three EHG signals, providing 33 candidate features. Fig 21 shows the average values of the misclassification errors, MCE, obtained on the training and test sets for both classification tasks. With respect to the obtained minima of the MCE functions for the training sets, all 22 available features were selected to classify between *early preterm* and *term* records (a minimum of 0.066 at 22nd feature), and the first 32 features to classify between *all preterm* and *term* records (a minimum of 4.368 at 32nd feature). Table 9 shows the selected features with their ranks for both classification tasks after using the frequency-based feature-aggregation and feature-selection procedure. For the task of classifying between *early preterm* and *term* records, the first two, fourth, fifth, seventh, and eighth, selected features were the sample entropies from all four frequency bands, while for the task of classifying between *all preterm* and *term* records the first six, ninth, and tenth selected features were again the sample entropies from all four frequency bands. The second most frequently selected feature was median frequency, and then the peak amplitude of the normalized power spectrum.

**Table 8.** The values of the Bhattacaryya criterion, *C*_B_, with its ranks per group of signals to separate *preterm* and *term, early* and *all* EHG records of the TPEHG DB.

| TPEHG DB |  |  |  |  |  |  |  |  |  |  |  |
| --- | --- | --- | --- | --- | --- | --- | --- | --- | --- | --- | --- |
| Signal | Early records (140 preterm / 143 term) |  |  |  |  |  |  |  |  |  |  |
|  | Band B0 |  | Band B1 |  |  | Band B2 |  |  | Band B3 |  |  |
|  | SE | MF | SE | MF | PA | SE | MF | PA | SE | MF | PA |
| S2 (*p*) | 0.206 | 0.156 | 0.161 | 0.622 | 0.031 | 0.044 | 0.127 | 0.837 | 0.192 | 0.004 | $\leq 1^{-3}$ |
| S3 (*p*) | $\leq 1^{-3}$ | 0.006 | $\leq 1^{-3}$ | 0.582 | $\leq 1^{-3}$ | $\leq 1^{-3}$ | 0.327 | $\leq 1^{-3}$ | 0.163 | 0.003 | $\leq 1^{-3}$ |
| S2 ($C_B$) | **0.011** | **0.036** | 0.005 | 0.002 | **0.021** | 0.002 | 0.004 | **0.014** | 0.005 | **0.010** | **0.011** |
| S3 ($C_B$) | **0.011** | **0.021** | 0.002 | 0.002 | **0.033** | 0.003 | 0.005 | **0.083** | 0.003 | **0.012** | 0.007 |
| S2 ($C_B$ rank) | **10** | **2** | 14 | 22 | **5** | 21 | 16 | **6** | 15 | **11** | **8** |
| S3 | **9** | **4** | 19 | 20 | **3** | 18 | 13 | **1** | 17 | **7** | 12 |

| Signal | All records (256 preterm / 262 term) |  |  |  |  |  |  |  |  |  |  |
| --- | --- | --- | --- | --- | --- | --- | --- | --- | --- | --- | --- |
|  | Band B0 |  | Band B1 |  |  | Band B2 |  |  | Band B3 |  |  |
|  | SE | MF | SE | MF | PA | SE | MF | PA | SE | MF | PA |
| S1 (*p*) | $\leq 1^{-3}$ | 0.049 | 0.551 | 0.011 | $\leq 1^{-3}$ | $\leq 1^{-3}$ | $\leq 1^{-3}$ | 0.011 | $\leq 1^{-3}$ | 0.384 | $\leq 1^{-3}$ |
| S2 (*p*) | $\leq 1^{-3}$ | 0.025 | $\leq 1^{-3}$ | $\leq 1^{-3}$ | 0.109 | $\leq 1^{-3}$ | 0.059 | 0.395 | $\leq 1^{-3}$ | 0.413 | $\leq 1^{-3}$ |
| S3 (*p*) | $\leq 1^{-3}$ | $\leq 1^{-3}$ | 0.009 | $\leq 1^{-3}$ | 0.006 | 0.019 | $\leq 1^{-3}$ | 0.002 | 0.007 | 0.885 | $\leq 1^{-3}$ |
| S1 ($C_B$) | $\leq 1^{-3}$ | 0.003 | $\leq 1^{-3}$ | $\leq 1^{-3}$ | **0.066** | 0.002 | $\leq 1^{-3}$ | **0.222** | **0.005** | **0.004** | **0.033** |
| S2 ($C_B$) | $\leq 1^{-3}$ | 0.002 | **0.004** | $\leq 1^{-3}$ | $\leq 1^{-3}$ | 0.003 | $\leq 1^{-3}$ | **0.017** | **0.006** | $\leq 1^{-3}$ | $\leq 1^{-3}$ |
| S3 ($C_B$) | $\leq 1^{-3}$ | **0.009** | 0.003 | $\leq 1^{-3}$ | **0.030** | $\leq 1^{-3}$ | $\leq 1^{-3}$ | **0.066** | 0.002 | 0.002 | $\leq 1^{-3}$ |
| S1 ($C_B$ rank) | 30 | 12 | 23 | 29 | **2** | 17 | 26 | **1** | **9** | **10** | **4** |
| S2 | 31 | 18 | **11** | 22 | 25 | 13 | 27 | **6** | **8** | 19 | 32 |
| S3 | 24 | **7** | 14 | 21 | **5** | 28 | 20 | **3** | 15 | 16 | 33 |

Those $p$ values $\leq 1^{-3}$ and the ranks of the first 11 features according to $C_B$ per group of signals are in bold.

**Fig 20.**
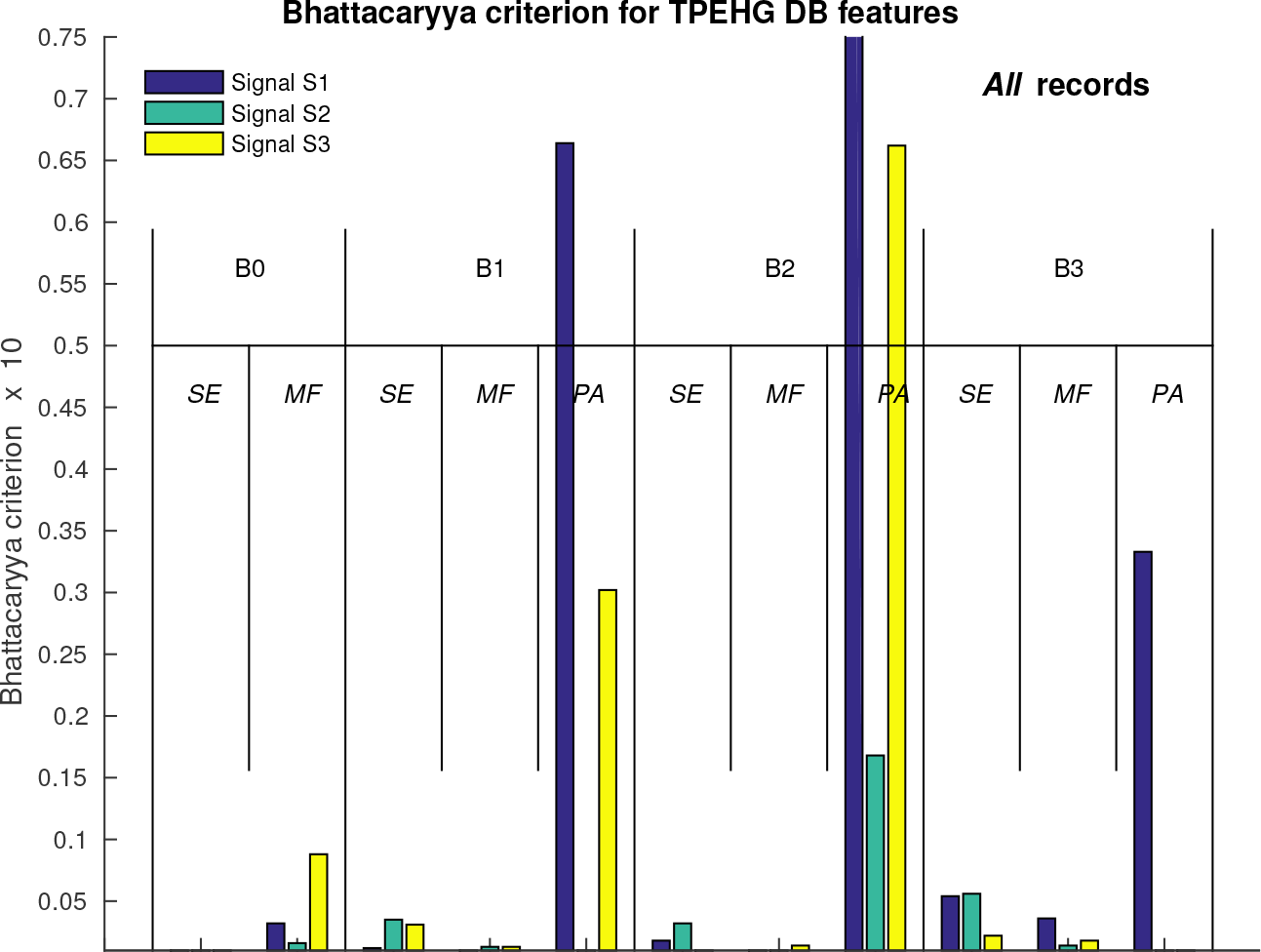
The values of the Bhattacaryya criterion, *C*_B_, for individual features to separate *all preterm* and *term* EHG records of the TPEHG DB.

**Fig 21.**
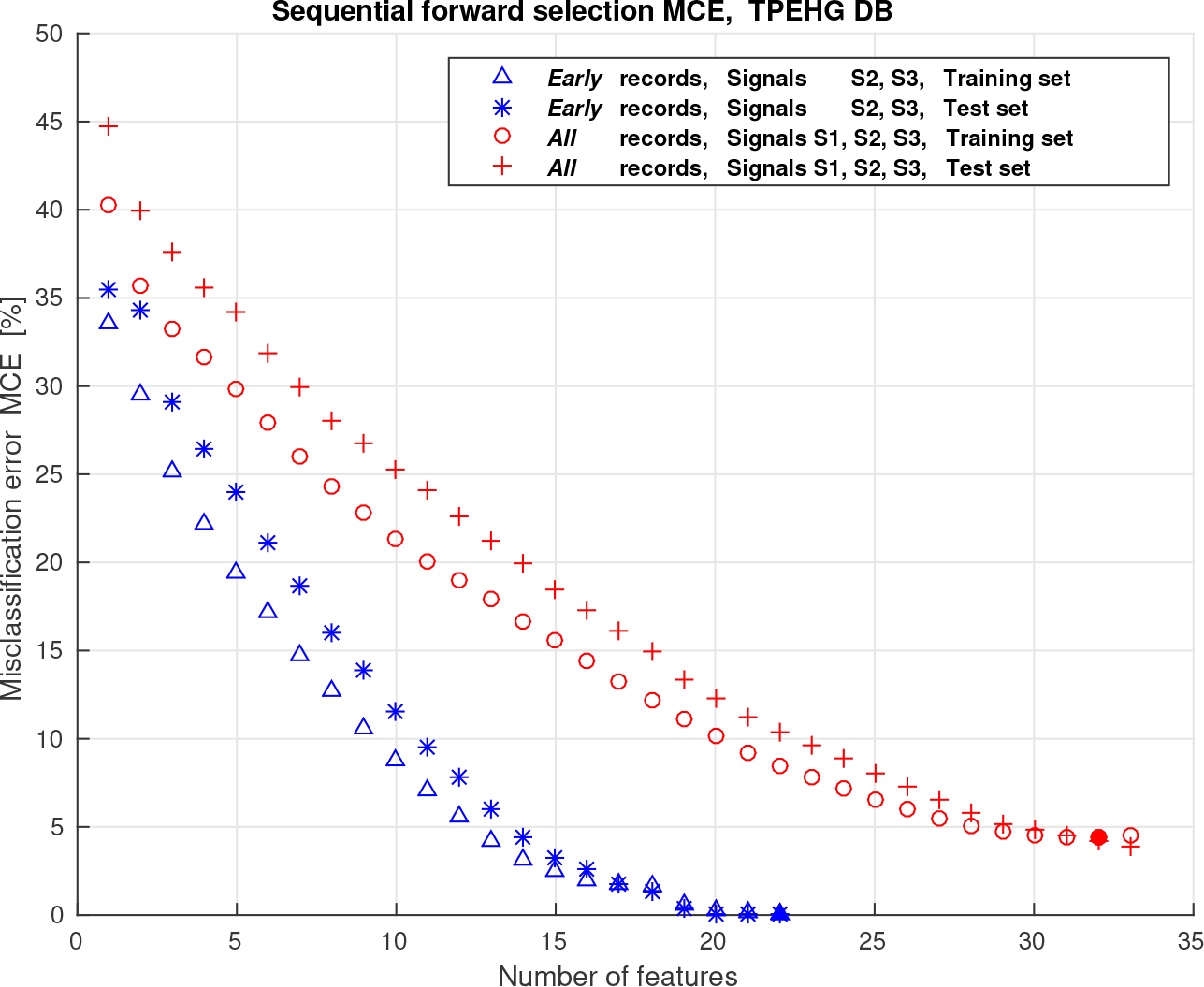
Average values of the misclassification errors, MCE, obtained on the training and test sets of the TPEHG DB using the SFS method.

**Table 9.** Selected features and their ranks to classify between *preterm* and *term, early* and *all* EHG records of the TPEHG DB.

| TPEHG DB |  |  |  |  |  |  |  |  |  |  |  |
| --- | --- | --- | --- | --- | --- | --- | --- | --- | --- | --- | --- |
| SFS (200 runs)<br>(QDA algorithm) | *Early records (140 preterm / 143 term)* |  |  |  |  |  |  |  |  |  |  |
|  | Band **B0** |  | Band **B1** |  |  | Band **B2** |  |  | Band **B3** |  |  |
|  | *SE* | *MF* | *SE* | *MF* | *PA* | *SE* | *MF* | *PA* | *SE* | *MF* | *PA* |
| **Signal S2** | **7** | **3** | **5** | 22 | 16 | **4** | 13 | 14 | **8** | **9** | 17 |
| **Signal S3** | **1** | 15 | **2** | **6** | 20 | **10** | 19 | 18 | 12 | **11** | 21 |
| SFS (200 runs)<br>(QDA algorithm) | *All records (256 preterm / 262 term)* |  |  |  |  |  |  |  |  |  |  |
|  | Band **B0** |  | Band **B1** |  |  | Band **B2** |  |  | Band **B3** |  |  |
|  | *SE* | *MF* | *SE* | *MF* | *PA* | *SE* | *MF* | *PA* | *SE* | *MF* | *PA* |
| **Signal S1** | **5** | 24 | 19 | 16 | 30 | **4** | 25 | 32 | **1** | **7** | 29 |
| **Signal S2** | 13 | 15 | **3** | 18 | 26 | 17 | 22 | 31 | **9** | 12 | 28 |
| **Signal S3** | **2** | **11** | **6** | **8** | 27 | 14 | 21 | - | **10** | 20 | 23 |

The first 11 selected features are in bold.

Table 10 summarizes the classification performance results obtained for *preterm* and *term* EHG records recorded *early*, and for *all preterm* and *term* EHG records of the TPEHG DB. For *early* records, and cross-validation with five- and ten-folds, the classification performances obtained were *CA* =100% (*AUC* =100%) and *CA* =100% (*AUC* =100%). For *all* records, and cross-validation with five- and ten-folds, the classification performances obtained were *CA* =95.75% (*AUC* =99.17%) and *CA* =96.33% (*AUC*=99.44%), respectively. Fig 22 shows classification accuracies obtained for *early* and *all* EHG records for different numbers of features, while the features were ranked according to the Bhattacaryya criterion. Finally, Table 11 summarizes the classification performances obtained by this and several other studies in classifying *preterm* and *term*, *early* and *all* EHG records of the TPEHG DB.

**Table 10.** Classification performance results obtained for *preterm* and *term*, *early* and *all* EHG records of the TPEHG DB.

| TPEHG DB (*preterm/term*) |  |  |  |  |  |  |  |  |
| --- | --- | --- | --- | --- | --- | --- | --- | --- |
| QDA | *Early* records<br>(140 *preterm* / 143 *term*) |  |  |  | *All* records<br>(256 *preterm* / 262 *term*) |  |  |  |
|  | Signals S2, S3 |  |  |  | Signals S1, S2, S3 |  |  |  |
| [%] | *Se* | *Sp* | *CA* | *AUC* | *Se* | *Sp* | *CA* | *AUC* |
| 5-folds | 100 | 100 | 100 | 100 | 92.97 | 98.47 | 95.75 | 99.17 |
| 10-folds | 100 | 100 | 100 | 100 | 94.14 | 98.47 | 96.33 | 99.44 |

**Fig 22.**
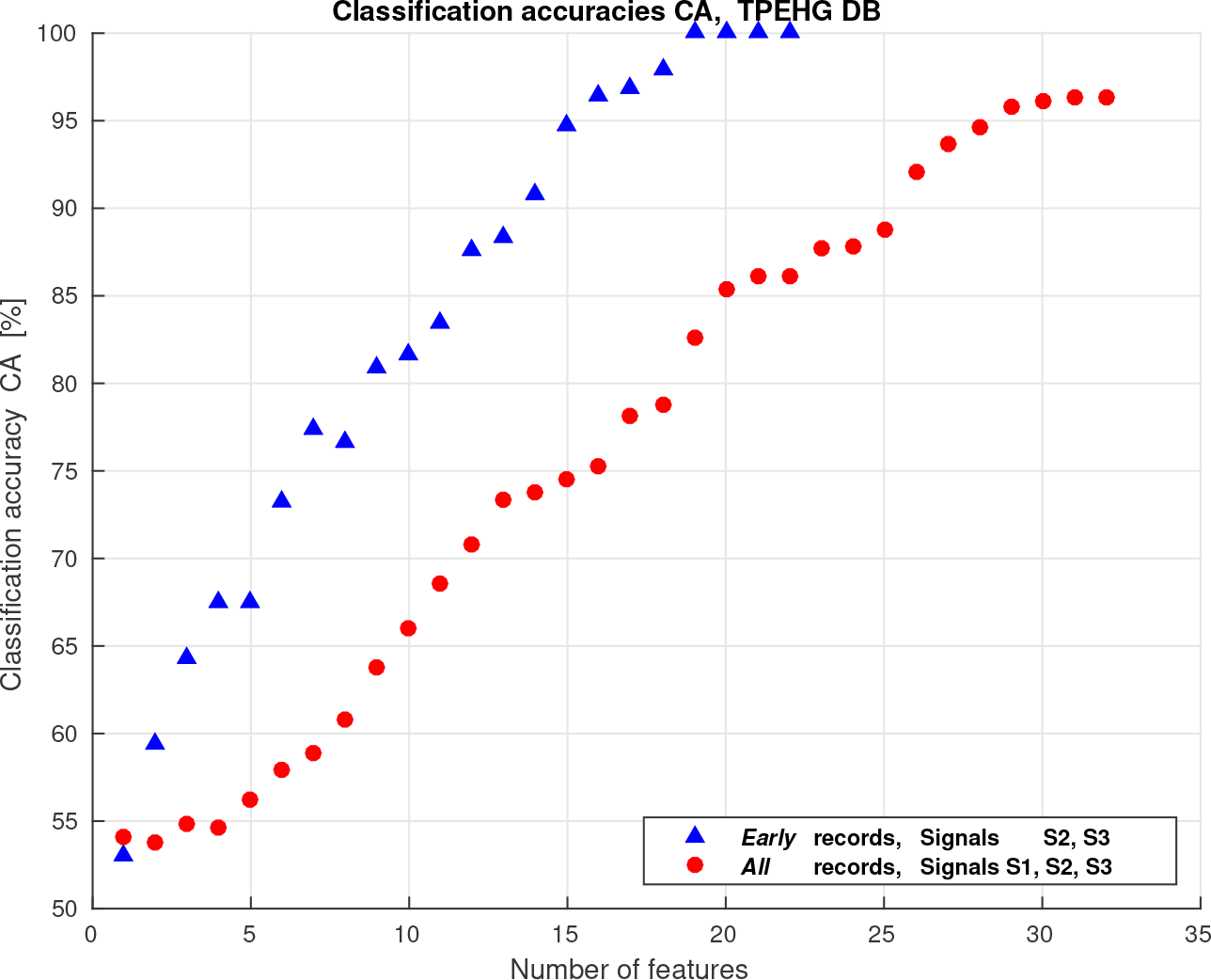
Classification accuracies for *early* and *all* EHG records of the TPEHG DB while the selected features were ranked according to the Bhattacaryya criterion.

**Table 11.** Comparison of classification performance results obtained for *preterm* and *term, early* and *all* EHG records of the TPEHG DB.

| TPEHG DB (*preterm/term*) |  |  |  |  |  |
| --- | --- | --- | --- | --- | --- |
| *Early* records |  |  |  |  |  |
| Method | Validation | $Se$ [%] | $Sp$ [%] | $CA$ [%] | $AUC$ [%] |
| Smrdel et al. [33] | 5-folds | 100 | 95 | 97 | - |
| Ahmed et al. [36] | 10-folds | 94 | 99 | 96.5 | 99 |
| The proposed method | 5-folds | 100 | 100 | 100 | 100 |
| *All* records |  |  |  |  |  |
| Method | Validation | $Se$ [%] | $Sp$ [%] | $CA$ [%] | $AUC$ [%] |
| Fergus et al. [29] | 5-folds | 97 | 90 | - | 95 |
| Hussain et al. [30] | 60% / 40% | 89 | 91 | 90 | - |
| Idowu et. al. [31] | leave-one-out | 97 | 86 | - | 94 |
| Smrdel et al. [33] | 5-folds | 96 | 79 | 87 | - |
| Ren et al. [34] | 10-folds | - | - | - | 98.6 |
| Fergus et al. [32] | 5-folds | 91 | 84 | - | 94 |
| Ahmed et al. [36] | 10-folds | 92 | 98 | 94.9 | 99 |
| Acharya et al. [35] | 5-folds | 91.39 | 97.33 | 94.47 | - |
|  | 10-folds | 95.08 | 97.33 | 96.25 | - |
| The proposed method | 5-folds | 92.97 | 98.47 | 95.75 | 99.17 |
|  | 10-folds | 94.14 | 98.47 | 96.33 | 99.44 |

## Discussion

### Measuring the influence of the maternal heart on the uterus

The influence of the maternal ECG signal on the uterine EHG activity is known [5,37]. In order to assess the intensity of the influence of the maternal ECG on the uterus in the electrical sense, it is necessary to measure the maternal heart’s influence within the EHG signals. A possible concern is whether the very specific spectral peak in the frequency band B1 for the EHG signals is present because of the maternal heart rate or some other electrophysiological phenomenon. The maternal heart is close to the uterus and produces a permanent, and the strongest, electrical signal in the body. The ratio between the amplitudes of the ECG and EHG signals is approximately 20, so it is likely that the maternal heart rate component with higher harmonics will be present. Considering *term* nonlabor uterine records of the TPEHGT DS, or *term early* EHG records of the TPEHG DB, a strong activity in the frequency band B1 for the EHG signals is present permanently and throughout the records (Figs 6 and S6 Fig), likely due to the maternal heart rate. In fact, according to the literature, within the EHG signals the following components can be measured [37]: the electrical activity of uterine bursts, the maternal respiration component, the maternal heart rate component with higher harmonics, and the EHG signal artefacts. No other electrical activity, from human organs that could be measured via EHG signals, was reported [37]. Another aspect is the influence of the maternal heart on the uterus in the mechanical sense. In this study, we showed that the influence of the maternal heart on the uterus in the mechanical sense is actually measurable via frequency domain analysis of the TOCO signal in the frequency region above 1.0 Hz, what is an important finding. Spectrograms (Fig 6) and power spectra of the TOCO signal (Figs 9, 12, 13, 15, and Table 1) for *term* nonlabor records of the TPEHGT DS clearly revealed the presence of a strong spectral component in the frequency band B1 which is due to the maternal heart rate.

In terms of the peak amplitudes of the normalized power spectra in the frequency bands B1, B2, and B3, the maternal heart’s influence seems to always be present in the EHG signals recorded on the abdomen of pregnant (Figs 5-8, 10, 11, 13, 15, S5 Fig, S6 Fig, and Table 1) and of non-pregnant women (Figs 14 and 15). The maternal heart’s influence is the strongest in the EHG signal S1 (Figs 13 and 15). Moreover, the sample entropies of the TOCO signal in the frequency band B1 for *contraction* and *dummy* intervals (Fig 16) are lower than the sample entropies of the EHG signals S1, S2, and S3, indicating the higher regularity of the TOCO signal in the frequency band B1, or the stronger influence of the maternal heart on uterus in a mechanical sense for pregnant women. Regarding the TOCO signal, it would be expected, for pregnant and non-pregnant women, that oscilations of the heart rate can be detected on the abdomen because of mechanical transmission through several organs as well as because of wave propagation through the vascular system. This is true for pregnant women. The peak amplitudes of the normalized power spectra of the TOCO signal for *contraction* and *dummy* intervals of pregnant women did reveal the influence of the maternal heart in the frequency band B1 (Figs 6, 9, 12, 13, 15, and Table 1). However, the results on the characterization of *dummy* intervals for non-pregnant women did not fully confirm these expectations. The peak amplitudes of the normalized power spectra of *non-pregnant dummy* intervals of the TOCO signal did not reveal any significant peak in the frequency bands B1, B2, or B3, where the maternal heart rate with harmonics would be expected (Fig 14). The mechanical influence does not seem to be present in the frequency band B1 (Fig 3 and S3 Fig). Moreover, the sample entropy for *non-pregnant dummy* intervals of the TOCO signal in the frequency band B1 is high (Fig 16), and much higher than the sample entropies of the TOCO signal for *preterm* and *term*, *contraction* and *dummy*, intervals of pregnant women, thus suggesting a very low regularity of the TOCO signal in the frequency band B1, or a very weak presence of a physiologic mechanism with periodic behavior for non-pregnant women. The maternal heart rate influence is present in the EHG and TOCO signals recorded on the abdomen for pregnant women, but evidently does not seem to be present in the TOCO signal for non-pregnant women. Classification between *non-pregnant dummy* intervals, and *preterm* and *term*, *contraction* and *dummy* intervals of pregnant women clearly confirmed these observations (Table 7). The classification performances (*CA* and *AUC*) were 100%, and close to 100%, for all classification tasks, suggesting that the characteristics of EHG and TOCO signals of uterine records are much different for non-pregnant and pregnant women. Moreover, if EHG signal S2 and TOCO signal were used for classification, only one feature, the sample entropy of the TOCO signal in the frequency band B2 or B3 (the presence or absence of the second or third harmonic of the maternal heart rate) was needed (Table 6) and yielded classification performance of 100% (Table 7). The latter result suggests that differences in the characteristics of uterine records for non-pregnant and pregnant women are even more significant for the TOCO signal, or, that the influence of the maternal heart on the uterus is barely present in the mechanical sense for non-pregnant women. In conclusion, the electro-mechanical influence of the maternal heart for pregnant women is higher for *term* pregnancies than for *preterm* pregnancies, and is the highest in the *term* nonlabor phase of pregnancy (Fig 13). The low level of this electro-mechanical influence for pregnant women suggests the approaching labor, or the danger of preterm birth. The monitoring of frequency spectra and spectrograms of the EHG and TOCO signals to look for the intensity of the influence of the maternal heart on the EHG and TOCO signals may become part of clinical investigation during the preliminary assessment of the danger of preterm birth.

### A new biophysical marker for the assessment of the risk of preterm birth

Normalization of the power spectra yielded an important feature, peak amplitude of the normalized power spectrum, *PA*, estimating relative proportions of peak amplitudes in different frequency bands. The characterization of the *PA* for *contraction* and *dummy* intervals in the frequency bands B1, B2, and B3, for the EHG and TOCO signals revealed a high level of the *PA* in all signals only for *term* nonlabor group of records (Figs 7 - 13). It may be thought that the rise of the *PA* in the frequency band B0 for the EHG signals occurs when labor approaches, and this is the reason for the drop of the *PA* in the frequency band B1. However, neither peak amplitude increase, nor peak median amplitude increase, for EHG signals were confirmed for contractions in the frequency band below 1.0 Hz for women who delivered within seven days [7,17,18,55]. In our study, the boundary between nonlabor and labor groups was set at three weeks. The level of the *PA* is low when delivery is close for *preterm* and *term* labor groups of records, and it is also low in *preterm* nonlabor group of records while the delivery is still far off (Fig 13 and Table 1). The level of the *PA* was the highest in the EHG signal S1, it was lower in signal S3, and the lowest in signals S2 and TOCO, for both, *contraction* and *dummy* intervals (Figs 13, 15, and Table 1). Differences in the influence of the maternal heart were the most prominent in the TOCO signal, and then in the EHG signal S2 (Table 1). The ratios between the medians of the *PA* in the frequency band B1, in the case of *term* nonlabor versus *preterm* nonlabor groups (Table 1), were the highest in the TOCO signal, and then in the EHG signal S2, for both types of intervals. They were 8.2 (TOCO signal) and 2.533 (signal S2) for *contraction* intervals, and were 5.6 (TOCO signal) and 4.0 (signal S2) for *dummy* intervals. The EHG signal S2 is a sign of electrical activity of the uterus along the vertical direction (the left electrodes, Fig 1). Since tocodynamometer, measuring mechanical uterine pressure, is attached at the top of the fundus, the TOCO signal can be regarded as measuring mechanical activity of the uterus also along the vertical direction. These are important observations. According to Table 1 (see bracketed average values), the threshold for the *PA* in the frequency band B1 for signals S2 and TOCO, to confirm term delivery, may be estimated as about 3% (signal S2) and about 7% (TOCO signal) for *contraction* intervals, and about 6% (signal S2) and about 5% (TOCO signal) for *dummy* intervals. Besides, the *PA* of the EHG signal S2, and of the TOCO signal, in the frequency band B1 have an explainable physiological mechanism in the background, i.e., influence of the maternal heart. Moreover, the most significant individual feature among all derived features for the classification between *preterm* and *term*, *contraction* and *dummy*, intervals with respect to Bhattacaryya criterion again appear to be the *PA* in the frequency bands B1, B2, and B3 (Fig 18 and Table 2). The achieved classification accuracies, using *dummy* intervals, and the individual feature *PA* from the frequency band B1 of the EHG signal S2, or of the TOCO signal, only, to classify between *preterm* nonlabor and *term* nonlabor pregnancies (Table 3), were *CA* =74.39%, or *CA* =71.95%. If using *dummy* intervals, and the individual feature *PA* from the frequency band B1 of the EHG signal S2 and of the TOCO signal, the achieved classification performances were the following: *Se*=90%, *Sp*=63%, *CA* =76.83%, and *AUC*=82.70%. Specificity is low, however, sensitivity is high. In addition, the most significant individual feature for the separation of the entire *preterm* and *term* EHG records of the TPEHG DB, as for *contraction* and *dummy* intervals of the TPEHGT DS, again appears to be the *PA* in the frequency bands B1, B2, and B3 (Fig 20 and Table 8) reflecting the influence of the electrical activity of the maternal heart. We propose the value of the peak amplitude of the normalized power spectrum of the EHG signal S2 estimating electrical propagation along the uterus in the vertical direction, and of the TOCO signal estimating mechanical activity of the uterus in the vertical direction, in the frequency band B1, describing the electro-mechanical activity of the uterus due to the maternal heart, as a new biophysical marker for the preliminary, or early, assessment of the risk of preterm birth. Moreover, the EHG signal S2, or the TOCO signal, could be used individualy for the preliminary assessment of the risk of preterm birth. Recently, according to literature, external tocography was thought to be an unpromising approach for predicting the risk of preterm birth [10,13]. However, this study brings back the importance of external tocography in predicting preterm birth. The TOCO signal carries important information about the mechanical influence of maternal heart activity during pregnancy, as well as the TOCO signal proved to be equally useful for the variety of classification tasks in combinantion with the EHG signals as the EHG signals are alone (Tables 5 and 7).

### Classification performance of the proposed method with respect to *contraction* and *dummy* intervals

Classifying between *preterm* and *term* uterine records is a very important factor for the efficient predicting of preterm birth. Several studies addressed this task using uterine contractions. In classifying between *preterm* nonlabor (38 patients), *preterm* labor (13 patients), *term* nonlabor (59 patients), and *term* labor (75 patients) groups (using peak frequency of the power spectrum up to 1.0 Hz, burst duration, number of bursts per unit time, total burst activity, and artificial neural network), the reported percentages of correct classifications were 71%, 92%, 86%, and 79%, respectively [18]. A recent study on evaluating the applicability of the EHG records for the early detection and classification of *preterm* and *term* birth during pregnancy included 20 women between the 24th and 28th week of pregnancy with threatened *preterm* labor [56]. The women were divided into two groups: those delivering within seven days and those delivering after more than seven days. To distinguish particular patterns for *preterm* and *term* EHG records, the analysis of the signals was performed using the combination of the recurrence quantification analysis and principal component analysis. The reported classification accuracy using the support vector machine classifier was 83.32%. Another study on evaluating the applicability of the EHG records in diagnosing threatening premature labor included three groups of pregnant women [16]. The first group was composed from 27 patients (from 27th to 40th week of pregnancy) with no symptoms of threatening *preterm* labor, the second group from 27 patients (from 23rd to 36th week of pregnancy) with the symptoms of threatening *preterm* labor, and the third group from 14 patients in the first labor phase of full time pregnancy. The classification performances in terms of AUC, using time domain (amplitude and area under contraction curve) and frequency domain (contraction power, median and peak frequency of power spectrum) features of uterine contractions derived from the EHG signals, and using support vector machine classifier, were 84.21% (groups one and two), 82.35% (groups one and three), and 57.14% (groups two and three). These results indicated that the properties of the contractions of the first labor phase group are similar to the properties of the contractions of the threatening *preterm* labor group. This finding is in accordance with the results of this study. Characteristics of *contraction* (and *dummy*) intervals are very similar for *term* labor intervals, and both, nonlabor and labor *preterm* intervals, but not for *term* nonlabor intervals. In our study, using the TPEHGT DS, we primarily addressed the problem of classifying between *preterm* and *term* uterine records using *contraction* and *dummy* intervals, as is the main objective of researchers when using the publicly available TPEHG DB database for the task of classifying between the entire *preterm* and *term* EHG records. The performance results achieved in classifying *preterm* and *term contraction* intervals of the TPEHGT DS, using the QDA classifier, and using signals S2 and S3, or, signals S2 and TOCO, in terms of AUC, were 94.04% (Table 5A), or, 95.65% (Table 5B), respectively. If using *dummy* intervals instead, for the same classification tasks, the performances were again quite comparable, and slightly higher on average. The performances obtained in the classification of *preterm* versus *term dummy* intervals, and using signals S2 and S3, or, signals S2 and TOCO, in terms of *AUC*, were 95.85% (Table 5A), or, 95.56% (Table 5B), respectively.

Moreover, several excellent studies addressed the problem of classifying between pregnancy (nonlabor) and labor uterine EHG records. Using uterine contractions of the EHG records, and the frequency band of 0.34-1.0 Hz, the reported performance in predicting preterm delivery within seven days (using peak frequency of the power spectrum and propagation velocity), in terms of *AUC* was 96% [10]. Considering the use of uterine contractions of EHG records, and the frequency band spreading below and above 1.0 Hz, examples of the highest reported performances in classifying between pregnancy and labor contractions in terms of correct classification were 88.72% (using Lyapunov exponent, variance entropy, wavelets related features, the QDA classifier, and 133 pregnancy and 133 labor contractions) [20]; in terms of *AUC* were 85% (using non-linear correlation coefficient, 174/115 pregnancy/labor contractions) [25], 84.2% (using time reversibility, 174/115 pregnancy/labor contractions) [21], 99% (using time reversibility, 30/30 pregnancy/labor contractions) [22]; and in terms of *CA* were 88.00% [26] (using intrinsic mode functions and the EMD method, and 150/150 pregnancy/labor contractions [57]). The performances obtained in this study (Table 5) in classification of labor versus nonlabor *contraction* intervals are quite comparable. Classification performances using the QDA classifier, and signals S2 and S3, or signals S2 and TOCO, in terms of AUC, were 87.79% (Table 5C), or 96.14% (Table 5D), respectively. If using *dummy* intervals, the classification performances obtained are again quite comparable, and slightly higher on average than those using *contraction* intervals. The classification performances in terms of *AUC* for signals S2 and S3, or signals S2 and TOCO, were 95.02% (Table 5C), or 94.61% (Table 5D), respectively. However, it is difficult to expose the best features, or the best feature selection method, or the highest classification performance, or the best performing classifier, or to compare the performances obtained, since the research groups (considering the classification of pregnancy and labor *contraction* intervals) developed and evaluated their methods using their own datasets, and the number of *contraction* intervals per dataset differed.

*Dummy* intervals, and the features obtained from the entire frequency spectrum available, up to 5.0 Hz, appear to be as suitable for the classification between *preterm* and *term*, and between nonlabor and labor, uterine records, as are *contraction* intervals. Even though *dummy* intervals are not suitable for use with other tehniques related to *contraction* intervals, like EHG propagation analysis [9] or EHG source localization [58], the results presented in this study suggest a novel and simple clinical technique, not necessarily to look for *contraction* intervals, but using *dummy* intervals for early assessment of the risk of preterm birth.

### Classification performance of the proposed method with respect to entire EHG records

When comparing the classification performances of different methods, it is very important that the methods are evaluated using the same reference database and all available records of the reference database, and using the same performance measures. Table 11 summarizes the classification performances of published methods to classify between entire *preterm* and *term* EHG records that were evaluated using all available EHG records of publicly available TPEHG DB. The bank of four-pole band-pass Butterworth filters of the proposed method for predicting preterm birth (Fig 4) decomposing each signal into a set of subspaces of the arbitrary selected bandwidths proved to be suitable for the task of feature extraction and classification, not only between *preterm* and *term*, *contraction* and *dummy* intervals of the TPEHGT DS, but also between the entire *preterm* and *term, early* and *all* records of the TPEHG DB. In comparison to the use of a single frequency band such as 0.34-1.0 Hz [29], or 0.3-3.0 Hz and MMFE [36], and in comparison to the use of multifrequency band decomposition approaches such as the EMD method [34,35], or wavelets [35], the proposed method yielded the classification accuracy of 100% for *early* records, and quite comparable and slightly higher classification performance, *CA* =96.33%, *AUC* =99.44%, for *all* records of the TPEHG DB (Table 11). Sharp cut-offs and high attenuation in the stopbands appear to be important characteristics to separate frequency bands when the signals have non-uniform spectral content due to physiological mechanisms residing in separate frequency bands, as in the case of EHG records. Activity due to contractions is expected in the frequency band B0 during *contraction* intervals, while the activity of the maternal heart with its harmonics is expected in the frequency bands B1, B2, and B3 throughout the records. (Recall that the entire EHG records of the TPEHG DB are actually compositions of *contraction* and *dummy* intervals.) The EMD method does not provide separate spectra bands of its intrinsic mode functions. Wavelets do not necessarily provide sharp cut-off and high attenuation in the stopbands with minimum overlaps of neighbouring frequency bands, nor do the set of subspaces provide arbitrary widths of the neighbouring frequency bands.

### Advantages of selected features

Another advantage of the proposed method to predict preterm birth lies in the features being extracted from each strictly separated frequency band. The features are capable of adequately estimating the presence (sample entropies), position (median frequencies), and intensity (peak amplitudes of the normalized power spectrum) of the physiological mechanisms residing in separate frequency bands. The most frequently selected features, and those with the highest rank using the SFS and frequency-based feature-aggregation and feature-selection procedure, for any of the classification task of this study, were the sample entropies (Tables 4, 6, and 9), followed by the median frequencies. Considering classification between *preterm* and *term*, labor and nonlabor, *contraction* and *dummy* intervals, the most significant features (Table 4) appear to be the sample entropies of the EHG signal S2 from the frequency bands B2 and B3 (the second and third harmonic of the maternal heart rate). Considering classification between *non-pregnant dummy* intervals, and *preterm* and *term, contraction* and *dummy* intervals, the most significant features (Table 6) again appear to be the sample entropies of the EHG signal S2 from the frequency bands B2 and B3, and from the TOCO signal in the frequency band B2. This result suggests that the presence of the second and third harmonic of the maternal heart rate is very important indicator for efficient differentiating between *preterm* and *term* deliveries. Even though the peak amplitudes of the normalized power spectra showed the highest individual classification ability, they were selected less frequently (Tables 4 and 9) or not at all (Table 6). The new proposed method for predicting preterm birth can become a novel clinical application. The method involves the analysis of uterine records containing EHG signals, or, EHG signals and the TOCO signal. Entire records may be processed fully automatically, or semi-automatically, if the phase of annotating *contraction* or *dummy* intervals is involved. In any case, it is suitable to assess the frequency content of signals using time-frequency domain visualizations using spectrograms of signals, with special attention on the frequency bands B1, B2, and B3 to quickly estimate the risk of preterm birth. Moreover, the proposed method yielded the classification accuracy of 100% for *early* records of the TPEHG DB (Table 11). This result suggests that the proposed method is suitable for predicting preterm birth already very early during pregnancy, i.e., around 23rd week of pregnancy. The highest possible classification accuracy was likely obtained due to the selected features, which directly estimate the presence and intensity of physiological mechanisms residing in strictly separated frequency bands. Another argument why the proposed method is suitable for predicting preterm birth very early during pregnancy lies in the features being extracted from frequency bands above 1.0 Hz, thus primarily estimating the influence of the maternal heart on the uterus. Since the features extracted from *dummy* intervals appear to be equally important for the classification between *preterm* and *term* records, as are the same features extracted from *contraction* intervals, the proposed method is suitable for clinical use very early during pregnancy, around 23rd week of pregnancy, when the contractions may or may not be present.

### Electrical activity of cervix

The cervix plays important role during pregnancy. The first stage of labour is the slow opening of the cervix, which happens with regular contractions of the uterus. When the cervix is fully opened (dilated), delivery follows. Cervical ripening starts early in pregnancy, in mid-gestation, with a process called softening [55]. Softening is followed by effacement and finally the dilation of the cervix. Ripening is required for the normal progression of labor. Cervical ripeness is quantified and scored according to the Bishop scoring system [59]. Unfortunately, our uterine records did not contain accompanying information about the cervix or Bishop score.

The electrical activity of the cervix is another important aspect. A previous EMG study showed that in humans the smooth muscles in the cervix act independently of those in the uterine corpus [38]. A study measuring EMG activity from the cervix reported the presence of a significant background level of electrical activity [60]. The study compared the cervical EMG activity in five women from induction through the latent phase of labour and into the active phase. During the latent phase (e.g., cervical dilation of 3 cm and cervical length of 1 cm), a significant background level of electrical activity was present between the increases in EMG activity due to bursts of cervical contractions in all cases. The amount of this background electrical activity was reduced with cervical ripening during the active phase of labor (e.g., cervical dilation of 4 cm and fully effaced) in all cases. Since the sampling frequency was 5 samples per second, and the frequency content of bursts of cervical contractions is below 1.0 Hz, this background electrical activity was likely due to the maternal ECG. This finding is in accordance with the results of this study. In *term* nonlabor case the maternal heart activity is present, while it is low or barely present in *term* labor case, and in both, nonlabor and labor, *preterm* cases. Cervical effacement and dilation could be assessed by estimating the level of normalized peak amplitude of the power spectra of the EHG and TOCO signals in the frequency bands B1, B2, and B3. A high level of this activity suggests nonlabor phase. A drop in this activity suggests labor phase, or the danger of preterm birth.

### Electro-mechanical activity of the uterus during *dummy* intervals

The modeling of electro-mechanical activity of the uterus is currently limited to uterine contractions. Several excellent studies connected to EHG propagation analysis [9], EHG source localization [58], and modeling of pregnancy contractions [61] exist.

Here we present an attempt to describe the electro-mechanical activity of a pregnant uterus within *dummy* intervals, out of contractions. Maternal heart activity, and the response of the uterus, can be described as an electrical pulse train (or waves) led into a system with non-linear transfer characteristic.

The immediate question is what properties of the uterus could cause the influence of the maternal heart rate in the electro-mechanical sense to be more or less measurable via the EHG and TOCO signals during pregnancy. We can look for the answer in those properties of uterus that change while the pregnancy progresses. There seems to be two candidates: the propagation velocity of the electrical potentials as they propagate over the uterine muscle, and the geometry and shape of the uterus and cervix (also responsible for higher harmonics). As the delivery approaches, the increased excitability of the uterine cells and increased connectivity among the cells result in the increased propagation velocity of electrical potentials [62]. It was shown that the change in propagation velocity does not appear earlier than seven days prior to delivery [10]. This study reported the propagation velocities of 11.11 ± 5.13 cm/sec and 11.31 ± 2.89 cm/sec for *preterm* and *term* nonlabor groups, and 52.56 ± 33.94 cm/sec and 31.25 ± 14.91 cm/sec for *preterm* and *term* labor groups of pregnant women, while the boundary between nonlabor and labor groups was set at seven days. The propagation velocities are practically equal for *preterm* and *term* nonlabor groups of records recorded seven or more days prior to delivery. In our study, the boundary between the nonlabor and labor groups of records of the TPEHGT DS was set at three weeks. This boundary resulted in 24 *dummy* labor intervals, i.e. 12 *dummy* intervals in *preterm* labor group and another 12 *dummy* intervals in *term* labor group. Of these 24 *dummy* intervals, only one interval relates to a pregnancy (record *tpehgt_p011*) for which the delivery appeared within one week. Therefore, we can assume that the propagation velocity for the remaining 23 *dummy* labor intervals is approximately equal to the propagation velocity of *dummy* intervals of *preterm* and *term* nonlabor groups of records. For these reasons, the propagation velocity could be excluded as a property causing changes in the influence of the maternal heart. Below, we consider the geometry and shape of the uterus and cervix only.

While the cervix is unripe and rigid, the uterus may be thought of as a closed womb with a discontinuity (cervix). Such a system responds with harmonics. Maternal ECG activity is strong, of the order of about 1 mV, while the EHG activity is of the order of about 50 *μ*V. Electrical pulses (or waves) of maternal heart activity, i.e. the electrical activity in the frequency band B1, propagate along the uterine muscle. Due to the closed womb with discontinuity the waves are reflected back, causing interference with themselves, and higher harmonics in the frequency bands B2 and B3 appear. Interference is a characteristic of waves of all types [63]. (It is beyond the scope of this study to discuss the role and intensity of each higher harmonic.) The TOCO signal is a sign of mechanical activity. In the frequency band B1, it carries information about the “vibrating” of the uterus in the vertical direction due to maternal heart activity. The presence of the electro-mechanical activity of the uterus due to the maternal heart in the frequency band B1, with higher harmonics in the frequency bands B2 and B3, was shown in this study for the EHG signals S1, S2, and S3, and for the TOCO signal in the frequency band B1 (Figs 6, 10 - 13, S6 Fig, and Table 1). The intensity of the maternal heart’s influence on the EHG and TOCO is the most prominent in *term* nonlabor phase. Due to the strong influence of maternal heart activity, the phase with a closed womb may be called the *interference phase* of pregnancy. While the cervix is unripe and rigid, maternal ECG activity is measurable via the EHG and TOCO.

While the pregnancy progresses the uterus is growing. With the approaching labor during the *term* labor phase, the intensity of the maternal heart’s influence on the EHG and TOCO diminishes together with harmonics, while during the *preterm* nonlabor and labor phases the intensity remains low throughout, or is barely present, as shown in this study (Figs 2, 5, 10 - 13, S5 Fig, and Table 1). During the ripening, the cervix effaces and slowly dilates. It means that the womb is opened. The waves caused by maternal heart activity diffract through the hole made by the effaced and dilated cervix.

Diffraction is a characteristic of waves of all types [63]. In this phase, the electro-mechanical activity in the frequency bands B1, B2, and B3 diminishes. The maternal heart’s influence on the EHG and TOCO is lower in all signals. Due to the diminished influence of the maternal heart activity, the phase with an opened womb may be called the *diffraction phase* of pregnancy. In this phase, the maternal heart’s influence on the EHG and TOCO remains the highest in the EHG signal S1, it is lower in signal S3, and the lowest in signals S2 and TOCO for *preterm* nonlabor and labor phases and for *term* labor phase (Fig 13 and Table 1). In the sense of mechanical information from the TOCO signal in the frequency band B1, the uterus is not “vibrating” any more. Since the EHG signal S2 (the left electrodes, Fig 1) estimates electrical propagation along the uterus in the vertical direction, as well as the TOCO signal estimates the intensity of mechanical activity of the uterus in the vertical direction, the drop of the maternal heart’s influence on the EHG and TOCO is likely due to the vertical diffraction of waves through the effaced and dilated cervix. Moreover, the EHG signals S1 (the top electrodes, Fig 1) and S3 (the bottom electrodes, Fig 1) estimate electrical propagation along the uterus in the horizontal direction, therefore the maternal heart’s influence in these two signals remains higher. Since the EHG signal S1 is recorded closer to the maternal heart, while the signal S3 is recorded closer to the cervix, the maternal heart’s influence remains the highest in the EHG signal S1. In conclusion, the drop in the influence of maternal heart activity is likely due the effacement and slow dilation of the cervix, and other shape changes of the cervix (shorter, and aligned with the birth canal) during the onset of labor, or during the latent phase of labor, or even earlier. This phase of weak maternal heart influence suggests the approaching labor phase, or the risk of preterm birth.

## Conclusion

The main purposes of this study were quantitative characterization of the uterine records of the TPEHGT DS, and the development and testing of a new method for predicting preterm birth. The innovations brought are: the newly developed TPEHGT DS; the establishment of a new biophysical marker for the preliminary assessment of the danger of preterm birth; confirmation of the hypothesis that the frequency region of uterine records above 1.0 Hz containing frequency components due to the influence of the maternal heart provides important features for efficient preterm delivery prediction; and the confirmation of the hypothesis that the dummy intervals are equally or even more important for predicting preterm birth than are the contraction intervals, thus suggesting a novel and simple clinical technique, rather than it being necessary to look for the contraction intervals. The most important contribution brought by this study is the improved method for predicting preterm birth. The method tested on the publicly available TPEHG DB outperformed all other currently existing methods. The method is also suitable for clinical use very early in pregnancy, around the 23rd week, when contractions may or may not be present.

Until now, other methods did not view the maternal ECG as something that needed to be considered. However, in this paper we showed that the influence of the maternal heart on the uterus in electrical and mechanical sense is an important physiological mechanism, and plays a very important role during pregnancy, and during the diagnosis of preterm birth. The characterization and classification results showed high correlation between preterm or term birth and the intensity of the influence of the maternal heart on the uterus. We believe that the findings described in this study will yield new studies, including our own, exploring the further understanding and modelling of the physiological mechanisms of the uterus that are involved during pregnancy, and their interference.

In the future, we will make efforts to provide a web server for the method proposed in this paper. In addition, we plan to develop a new, larger database that will contain the TPEHGT DS dataset, TPEHG DB database, and many new uterine records (EHG signals and TOCO signal), recorded twice given pregnancy (around 23rd and around 31st week of pregnancy), with spontaneous (*preterm* and *term*), induced, and cesarean section deliveries. This database will allow many new studies connected to the further characterization of uterine records and further development of the advanced methods for the efficient prediction of preterm birth.

## Supporting information

**S1 Fig.**
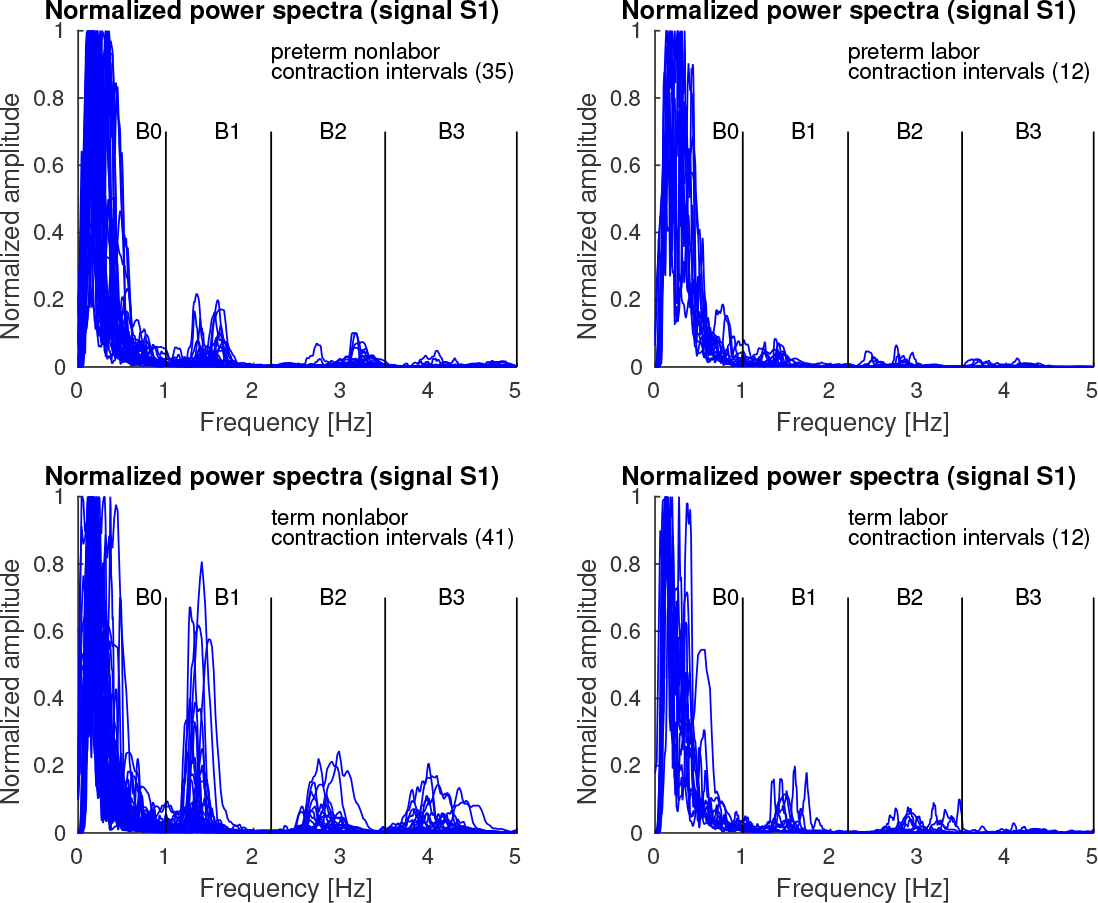
Normalized power spectra of *contraction* intervals of signal S1 of the records of the TPEHGT DS.

**S2 Fig.**
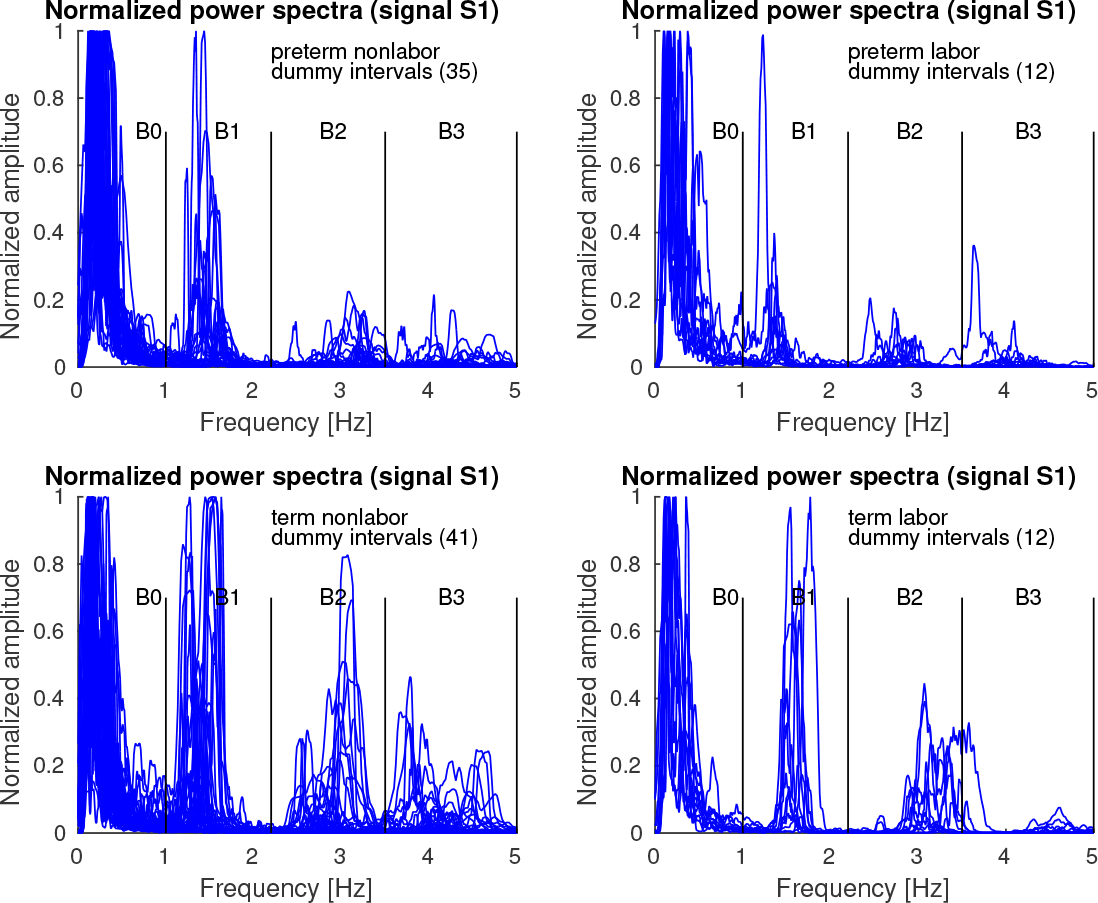
Normalized power spectra of *dummy* intervals of signal S1 of the records of the TPEHGT DS.

**S3 Fig.**
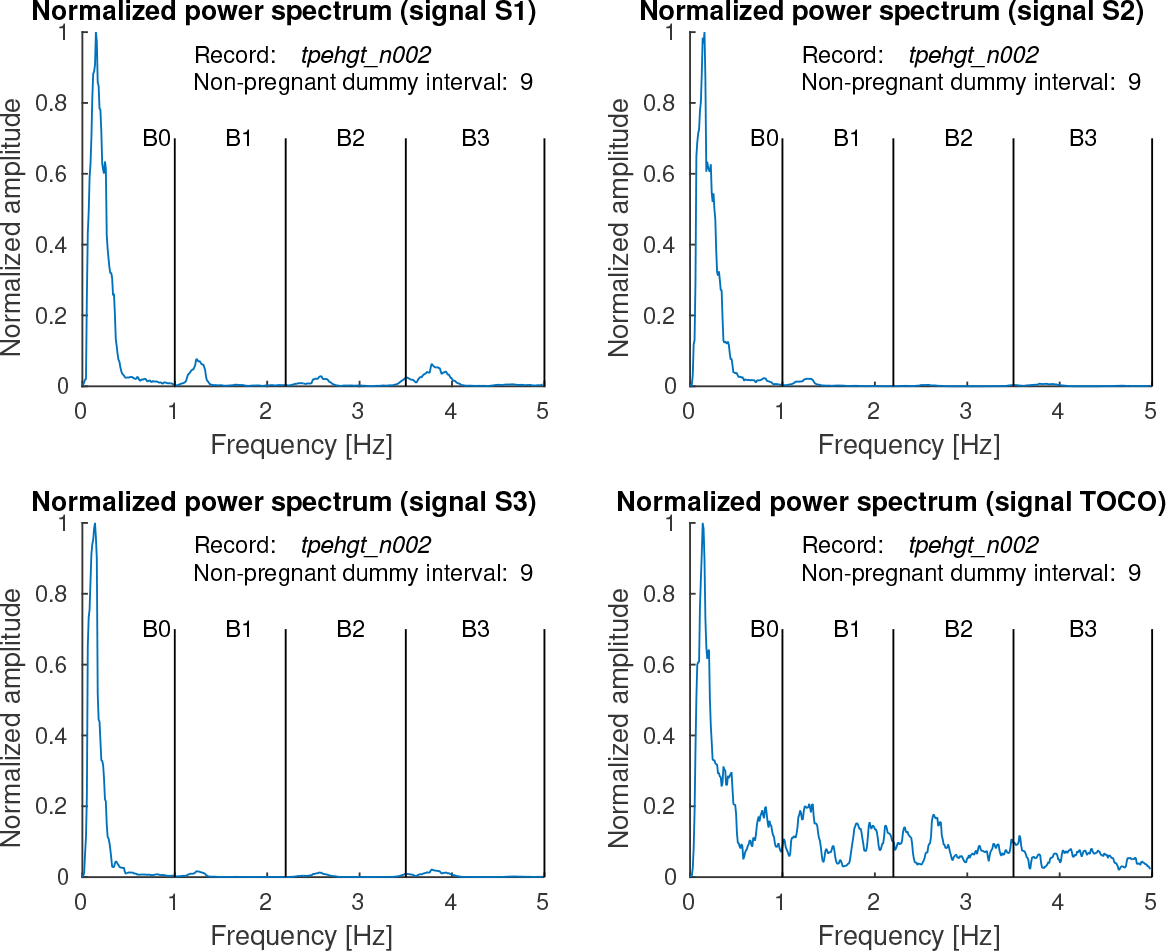
Normalized power spectra of signals S1, S2, S3, and TOCO, of the ninth *non-pregnant dummy* interval of the record *tpehgt_n002* from Fig 3.

**S1 File. Filtered signals S1, S2, S3, and TOCO, of the ninth *non-pregnant dummy* interval of the record *tpehgt_n002* from Fig 3.**

**S2 File. Normalized power spectra of filtered signals S1, S2, S3, and TOCO, of the ninth *non-pregnant dummy* interval of the record *tpehgt_n002* from Fig 3.**

**S4 Fig.**
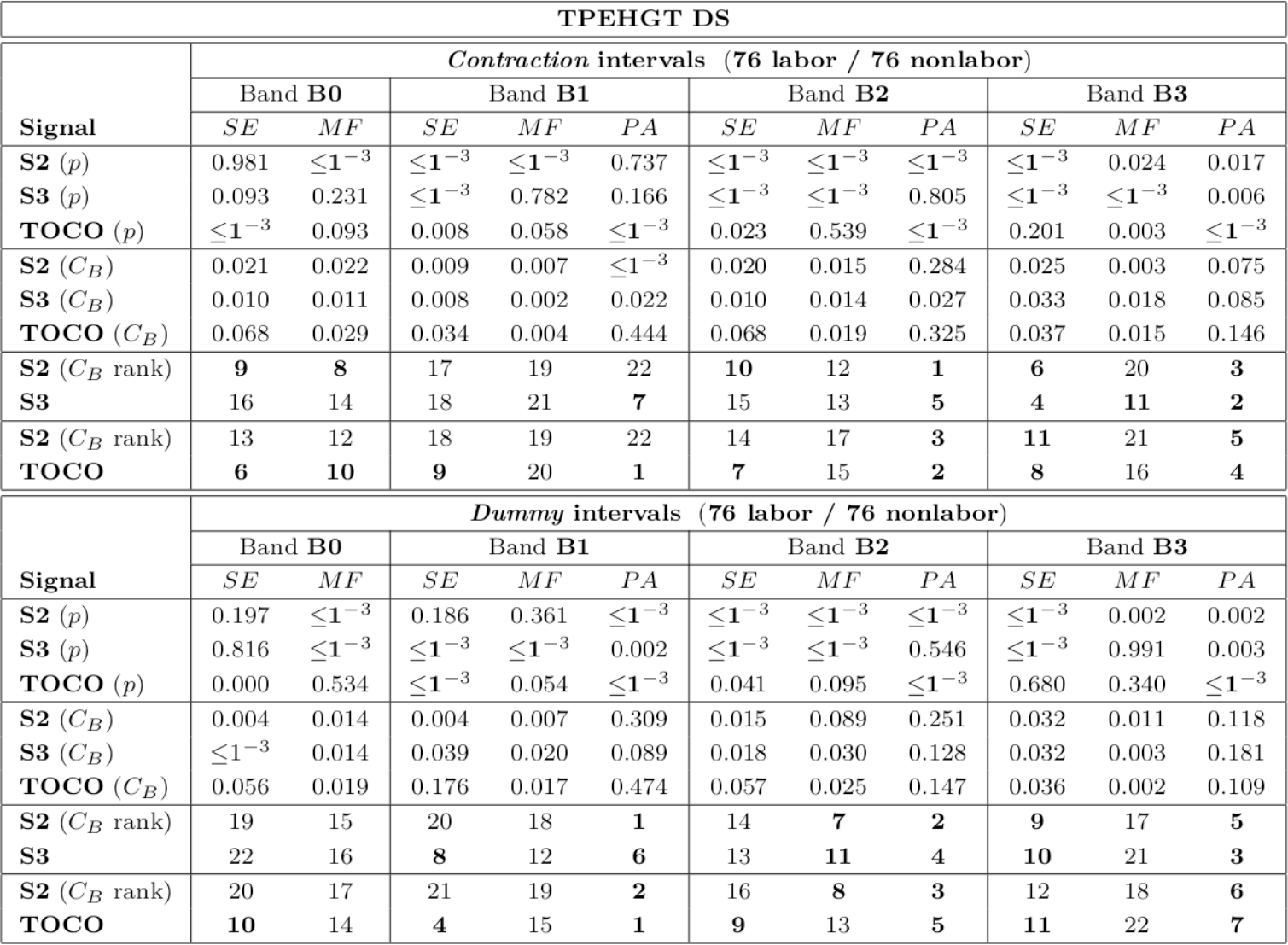
The values of the two-sample *t*-test (*p* values), and of the Bhattacaryya criterion, *C*_B_, with its ranks per group of signals, to separate labor and nonlabor, *contraction* and *dummy* intervals of the TPEHGT DS. Those *p* values ≤ 1^−3^ and ranks of the first 11 features according to *C*_B_ per group of signals are in bold.

**S5 Fig.**
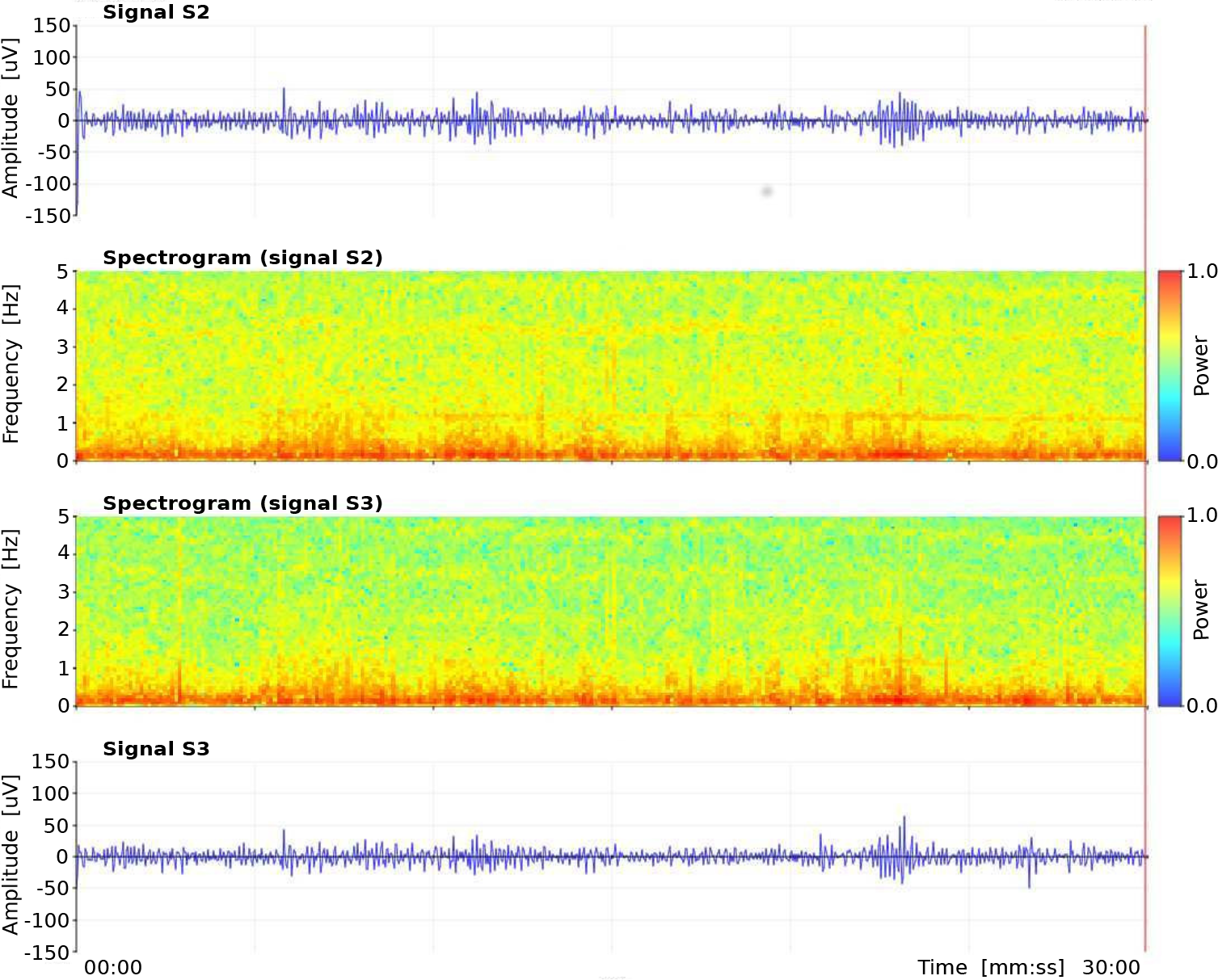
The EHG signals and their spectrograms of the record *tpehg1526* (*preterm*, recorded *early* in the 25th week, delivery in the 30th week) of the TPEHG DB. From top to bottom: EHG signal S2, spectrogram (0.0-5.0 Hz) of EHG signal S2, spectrogram (0.0-5.0 Hz) of EHG signal S3, EHG signal S3.

**S6 Fig.**
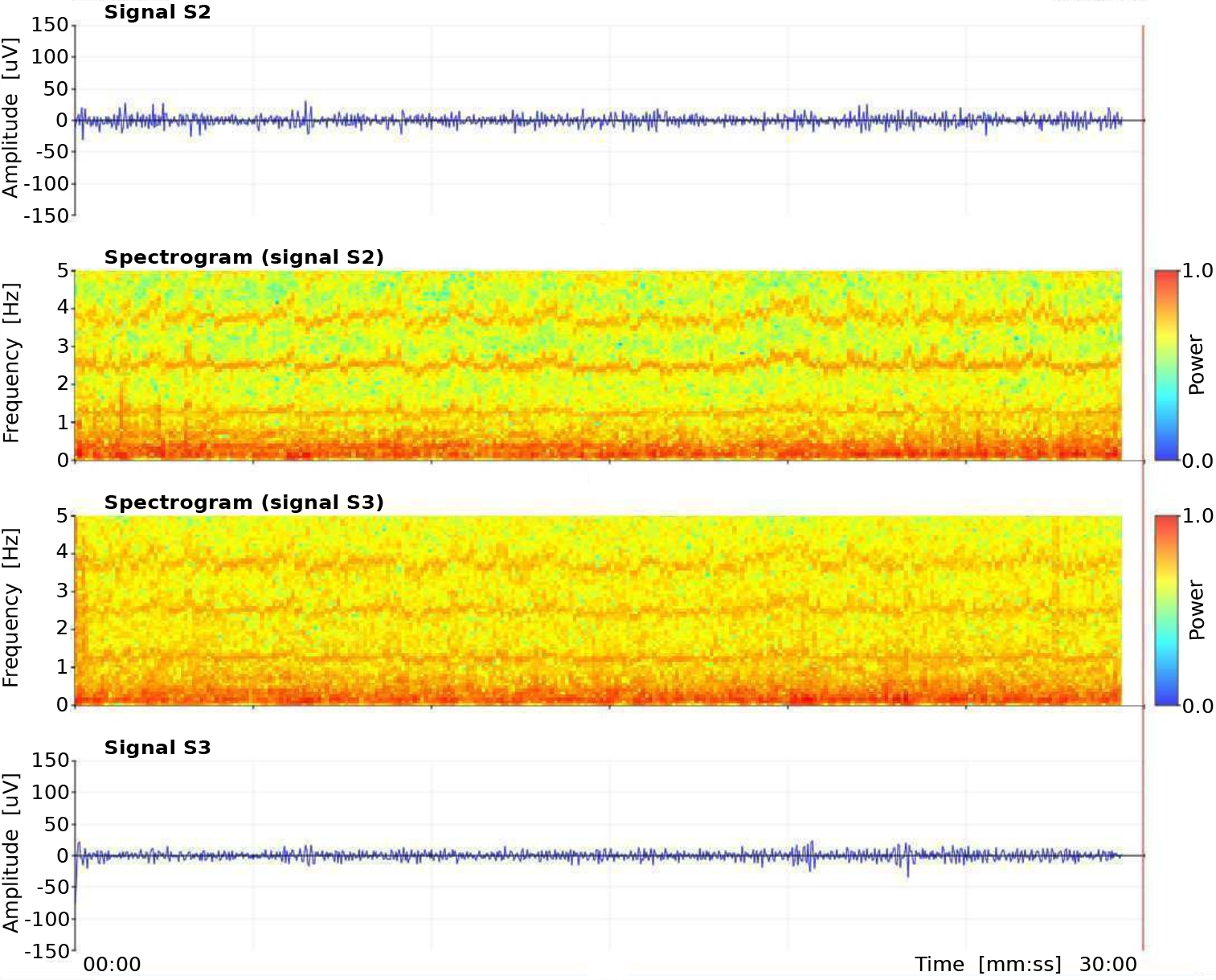
The EHG signals and their spectrograms of the record *tpehg949* (*term*, recorded *early* in the 25th week, delivery in the 39th week) of the TPEHG DB. From top to bottom: EHG signal S2, spectrogram (0.0-5.0 Hz) of EHG signal S2, spectrogram (0.0-5.0 Hz) of EHG signal S3, EHG signal S3.

